# scTensor detects many-to-many cell–cell interactions from single cell RNA-sequencing data

**DOI:** 10.1101/2022.12.07.519225

**Authors:** Koki Tsuyuzaki, Manabu Ishii, Itoshi Nikaido

## Abstract

Complex biological systems are described as a multitude of cell–cell interactions (CCIs). Recent single-cell RNA-sequencing studies focus on CCIs based on ligand–receptor (L-R) gene co-expression but the analytical methods are not appropriate to detect many-to-many CCIs.

In this work, we propose `scTensor`, a novel method for extracting representative triadic relationships (or hypergraphs), which include ligand-expression, receptor-expression, and related L-R pairs. Through extensive studies with simulated and empirical datasets, we have shown that `scTensor` can detect some hypergraphs that cannot be detected using conventional CCI detection methods, especially when they include many-to-many relationships. `scTensor` is implemented as a freely available R/Bioconductor package.

## Background

Complex biological systems and processes such as tissue homeostasis [1, 2], neurotransmission [3, 4], immune response [5], ontogenesis [6], and stem cell niches niche [7, 8] are composed of cell–cell interactions (CCIs). Many molecular biology studies have decomposed such systems into constituent parts (e.g., genes, proteins, and metabolites) to clarify their functions. Nevertheless, more sophisticated methodologies are required because CCIs essentially differentiate whole systems from functioning merely as the sum of their parts. Accordingly, micro-level measurements of such parts cannot always explain macro-level biological functions.

Previous studies have investigated CCIs using technologies such as fluorescence microscopy [9–13], microdevice-based methods such as microwells, micropatterns, single-cell traps, droplet microfluidics, and micropillars [14–22], and transcriptome-based methods [23–52]. In particular, the recent single-cell RNA-sequencing (scRNA-seq) studies have focused on CCIs based on ligand–receptor (L-R) gene co-expression. By investigating the detected cell types through scRNA-seq and the L-R pairs specifically expressed in the cell types, CCIs can potentially be understood at high resolution.

Despite their wide usage, the analytical methods based on L-R pairs are still not mature; such methods implicitly assume that CCIs consist of one-to-one relationships between two cell types and that the corresponding L-R co-expression is observed in a cell-type-specific manner. One study even removed ligand and receptor genes expressed in multiple cell types from their data matrix, assuming one-to-one CCIs [53]. In real empirical data, however, each ligand and receptor gene can be expressed across multiple cell types, and some studies have actually focused on many-to-many CCIs [25, 33, 36, 48, 54]. Such a difference between actual CCI patterns composed of real data and the hypothesis assumed by a model will cause severe bias in the detection of CCIs.

For the above reason, we propose `scTensor`, which is a novel CCI prediction method based on a tensor decomposition algorithm. Our method regards CCIs as hypergraphs and extracts some representative triadic relationships consisting of ligand-expression, receptor-expression, and related L-R pairs. The main contributions of this article are summarized as follows.

- We developed a novel simulator to model the CCIs as hypergraphs and quantitatively evaluate the performance of `scTensor` and other L-R detection methods.
- We re-implement some L-R detection methods from scratch in order to analyze the same L-R database with all of these methods and focus on only the performance of L-R detection methods, not the slight differences in data pre-processing and the L-R database used.
- We show that `scTensor`’s performance with respect to its accuracy of many-to-many CCI detection, computation time, and memory usage are superior to the other L-R detection methods.
- We describe the implementation of `scTensor` as an R/Bioconductor package to enable the reproducibility of data analyses as well as continuous maintenance and improvements. We provide some original visualization functions and a function to generate an HTML report in `scTensor` to enable detailed interpretation of the results. We have extended our framework to work with 125 species.

## Results

### CCI as a hypergraph

One of the simplest CCI representations is a directed graph, where each node represents a cell type and each edge represents the co-expression of all L-R pairs (Figure 1a, left). The direction of each edge is set as the ligand expressing cell type → the receptor-expressing cell type. Such a data structure corresponds to an asymmetric adjacency matrix, in which each row and column represents a ligand-expressing cell type and receptor-expressing cell type, respectively. If some combinations of cell types are regarded as interacting, the corresponding elements of the matrix are filled with 1 and otherwise 0. If the degree of CCI is not a binary relationship, weighted graphs and corresponding weighted adjacent matrices may also be used.

**Figure 1.**
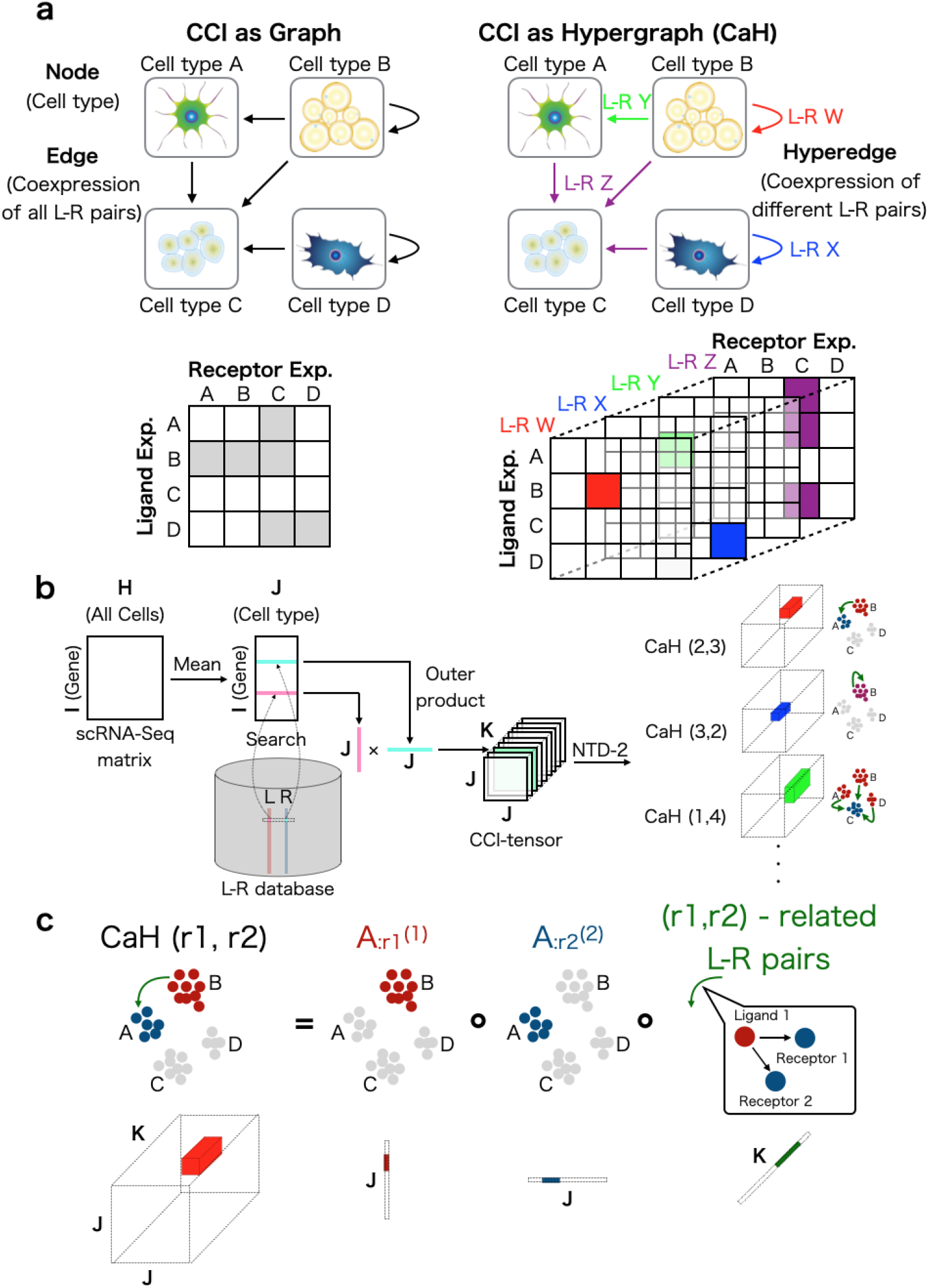
Cell–cell interaction (CCI) as a hypergraph (CaH). (a) Previous scRNA-seq studies have regarded CCIs as graphs, and the corresponding data structure can be expressed as an adjacency matrix (left). In this work, CCIs are regarded as context-aware edges (hypergraphs), and the corresponding data structure is a tensor (right). (b) The CCI-tensor is generated by users’ scRNA-Seq matrices, cell-type labels, and ligand–receptor (L-R) databases. NTD-2 is used to extract `CaH`s from the CCI-tensor. (c) Each `CaH`(r1,r2) is equal to the outer product of three vectors. 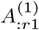 represents the ligand expression pattern, 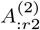 represents the receptor expression pattern, and *G*_*r*1,*r*2_,: represents the patterns of related L-R pairs.

The previous analytical methods are categorized within this approach [23, 24, 26–34, 36, 40, 43–46, 48, 49, 51, 52, 55].

The drawback of using an adjacency matrix to describe CCIs is that multiple L-R co-expression scores are collapsed into a single value by summation or averaging. Because the average is simply a constant multiple of the sum, here we discuss only the sum. The summed value has no meaning in which L-R pairs are related to the CCI, and therefore CCIs and the related L-R pair lists cannot be detected simultaneously.

In contrast to an adjacency matrix (i.e., graph), the triadic relationship of CCIs also can be described as directed hypergraphs (i.e., CCI as hypergraph; `CaH`), where each node is a cell type but the edges are distinguished from each other by the different related L-R pair sets (Figure 1a, right). Such a context-aware edge is called a “hyperedge” and is described as multiple different adjacency matrices. The set of matrices corresponds to a “tensor”, which is a generalization of a matrix to expand its order.

### Overview of scTensor

Here we introduce the procedure of `scTensor`. Firstly, a tensor data is constructed through the following steps (Figure 1b). A scRNA-seq matrix and the cellular labels specifying cell types are supposed to be provided by users. Firstly, the gene expression values of each cell are normalized by count per median of library size (CPMED [56–58]) and logarithm transformation, for variance-stabilization, is performed to the data matrix (i.e., log_10_ (CPMED + 1)).

Next, the data matrix is converted to a cell-type-level average matrix according to the cell type labels. Combined with an L-R database, two corresponding rowvectors of an L-R pair are extracted from the matrix. The outer product (direct product) of the two vectors is calculated, and a matrix is generated. The matrix can be considered as the similarity matrix of all possible cell-type combinations for each L-R pair. Finally, for each L-R pair, the matrix is calculated, and the tensor *χ* ∈ ℝ^*J×J×K*^, where *J* is the number of cell types and *K* is the number of L-R pairs, is generated as the merged matrices. In this work, this tensor is called the “CCI-tensor”.

After the construction of the CCI-tensor, we use the non-negative Tucker2 decomposition (NTD-2) algorithm [59, 60]. NTD-2 decomposes the CCI-tensor as a core tensor 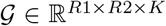, and two factor matrices ***A***^(1)^ ∈ ℝ^*J×R*1^ and ***A***^(2)^ ∈ ℝ^*J×R*2^, where *R*1 and *R*2 are the NTD-2 rank parameters. The factor matrix ***A***^(1)^ describes the *R*1 of ligand gene expression patterns in each cell type and the factor matrix ***A***^(2)^ describes *R*2 of receptor gene expression patterns in each cell type, and core tensor G describes the degree of association of all the combination (*R*1 × *R*2) of the ligand and receptor expression patterns of each L-R pair.

The result of NTD-2 is considered the sum of some representative triadic relationships. In this work, each triadic relationship is termed `CaH` (*r*1, *r*2), which refers to the outer product of three vectors, 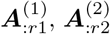, and 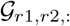, where *r*1 (1 ≤ *r*1 ≤ *R*1) and *r*2 (1 ≤ *r*2 ≤ *R*2) are the indices of the columns of the two factor matrices (Figure 1c). The `CaH`s are extracted in a data-driven way without the assumption of one-to-one CCIs. Therefore, this approach can also detect many-to-many CCIs according to the data complexity.

### Evaluation of many-to-many CCIs detection

To examine the performance of the CCI methods in terms of detecting `CaH`, we validated the CCI methods using both simulated and empirical datasets (Figure 2).

**Figure 2.**
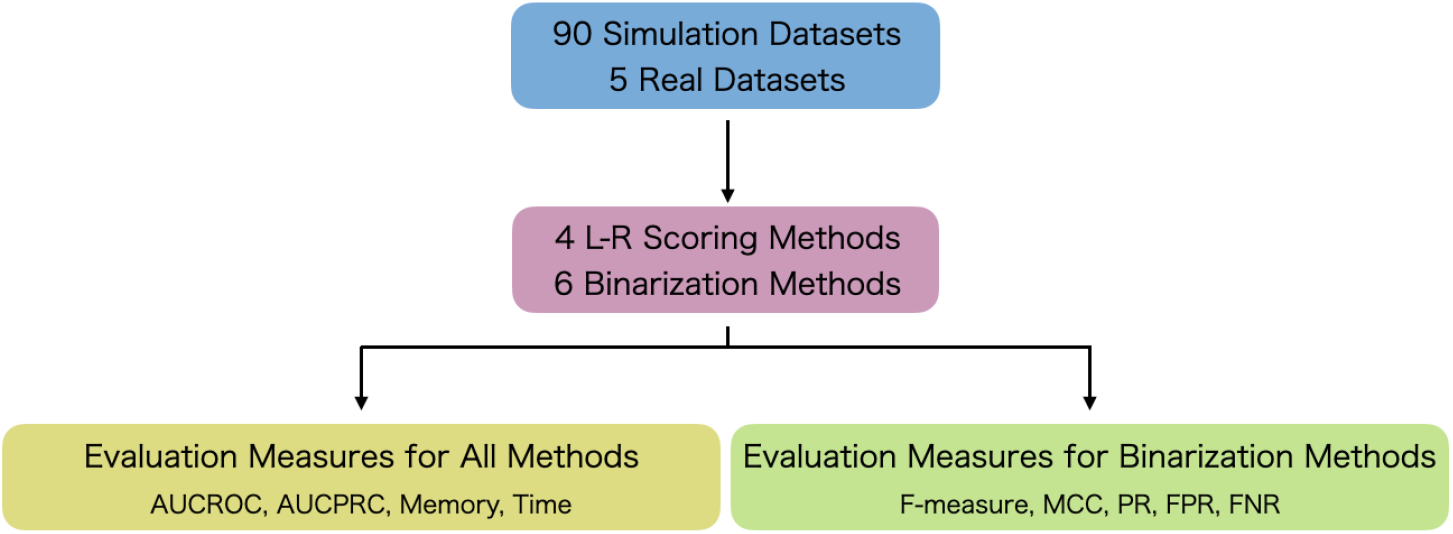
Evaluation scheme. To evaluate CCI methods, 90 simulated datasets and six real empirical datasets were prepared. Four ligand–receptor (L-R) scoring methods and six binarization methods were then evaluated. For the evaluation of these methods, area under the receiver operating characteristic curve (AUCROC), area under the precision-recall curve (AUCPRC), memory usage, and computational time were determined. For the evaluation of binarization methods, F-measure, Matthews correlation coefficient (MCC), positive ratio (PR), false positive ratio (FPR), and false negative ratio (FNR) were determined.

We first prepared 90 simulated datasets considering five numbers of cell types (3, 5, 10, 20, or 30), two CCI styles (one-to-one or many-to-many), three numbers of CCI types (1, 3, or 5), and three threshold values (E2, E5, or E10) for recognition of differentially expressed genes (DEGs). According to these conditions, ground truth CCIs were determined (Additional File 1).

Next, we prepared five real empirical datasets (`FetalKidney` [36], `GermlineFemale` [25], `HeadandNeckCancer` [54], `Uterus` [33], and `VisualCortex` [48]), each of which focused on many-to-many CCIs in their respective original papers.

There are many L-R scoring methods to quantify the degree of co-expression of ligand and receptor genes. We re-implemented four scores used in many CCI prediction methods to evaluate performance independent of software implementation. Here, we selected as the four methods `sum score` (CellPhoneDB [37], Giotto [61], CrossTalkR [62], and Squidpy [63]), `product score` (NATME [64], FunRes [65], ICELLNET [66], and TraSig [67]) `Halpern’s score`([68]), and `Cabello–Aguilar’s score` (SingleCellSignalR [69] and CellTalkDB [70]), each of which is widely used in many studies (see Additional File 2 for the simulated datasets).

To differentiate significant CCIs from non-significant CCIs, many CCI methods introduce a label permutation test with a random permutation of cell-type labels to simulate the null distribution of CCIs. This process is considered a kind of binarization (1 for significant CCIs, 0 for non-significant CCIs). For `scTensor`, binarization was realized by median absolute deviation (MAD) thresholding against each column vector in factor matrices calculated by tensor decomposition.

To quantitatively evaluate how selectively each CCI method was able to detect the ground truth CCIs before and after binarization, nine evaluation measures were introduced. Four of them were applied both before and after binarization, and the remaining five were applied to the results only after binarization.

#### scTensor could selectively detect many-to-many CCIs in simulated datasets

The area under the curve of precision-recall (AUCPR) and Matthews correlation coefficient (MCC) values of 30 datasets with an E10 threshold value are shown in Figure 3. For the details of all the evaluation results for all the conditions, see Additional Files 3 - 12. Figure 3 shows that the AUCPR values can vary among the CCI methods. When the CCI-style was one-to-one (Figure 3a, left), `scTensor` (NTD-2) achieved the highest AUCPR scores, and `Halpern’s score` obtained the second-highest AUCPR values on average. For `Halpern’s score`, however, binarization has significantly reduced the significant CCIs. This may be explained by `Halpern’s score` having the lowest FPR (Additional File 10) and the highest FNR (Additional File 11), and it suggests that `Halpern’s score` is a quite conservative method to detect one-to-one CCIs. When the CCI-style was set as many-to-many, both the previous and current versions of `scTensor` (NTD-3 and NTD-2, respectively) achieved higher AUCPR values on average (Additional File 4), compared with the other methods (Figure 3a, right).

**Figure 3.**
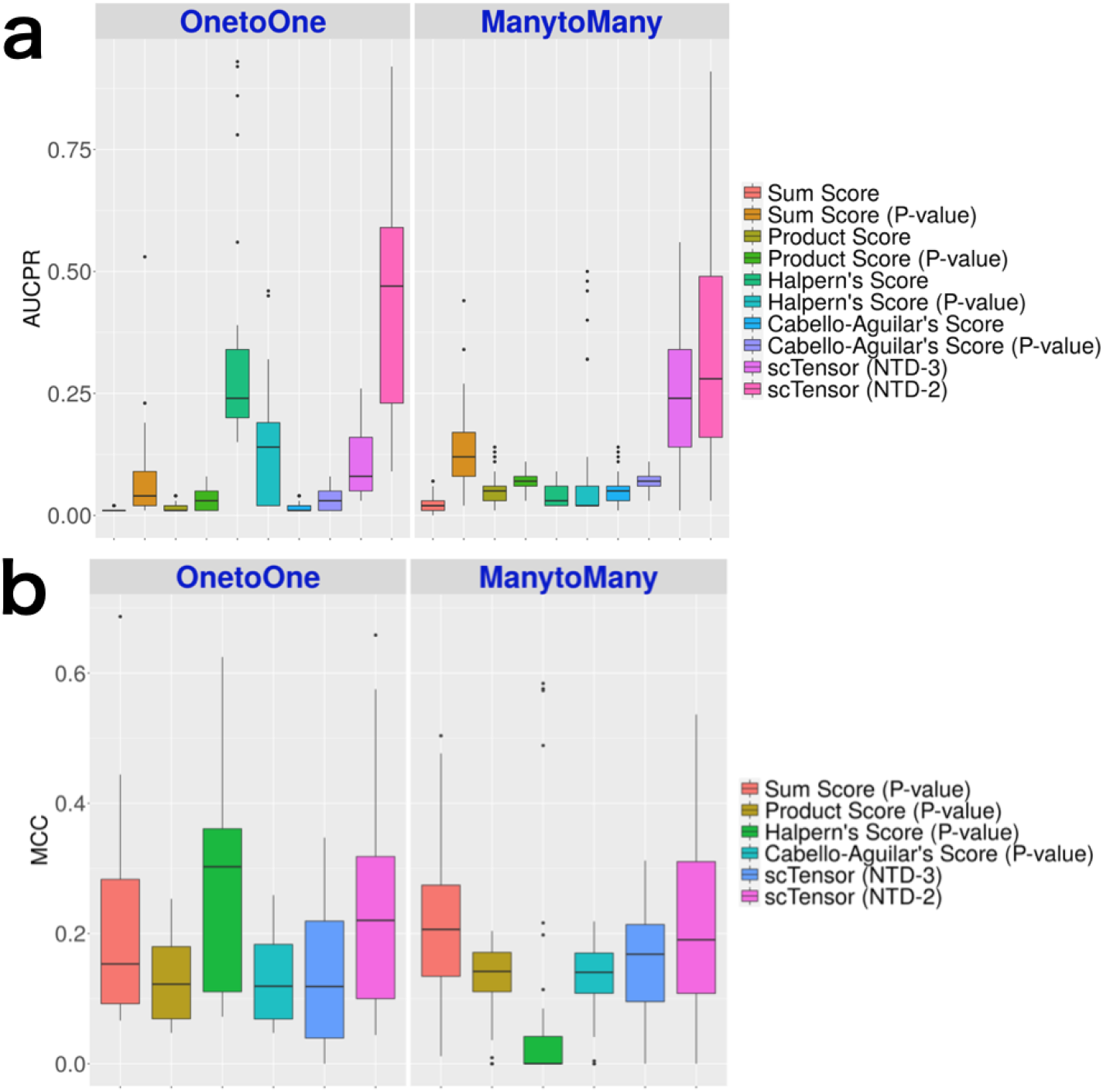
Results of simulated datasets. (a) Area under the curve of precision-recall (AUCPR) of all the methods. (b) Matthews correlation coefficient (MCC) of the binarization methods.

Figure 3b shows that the MCC values also varied among the CCI methods. When the CCI-style was one-to-one (Figure 3b, left), `Halpern’s score` achieved the highest MCC values and `scTensor` (NTD-2) obtained the second-highest values on average (Additional File 8). When the CCI-style was many-to-many, `scTensor` (NTD-2) and `sum score` obtained the highest MCC values, compared with the other methods (Figure 3b, right).

#### Characteristics of scTensor (NTD-2), Halpern’s score, and sum score

The comprehensive validation described that the three methods (`scTensor` (NTD-2), `Halpern’s score`, and `sum score`) performed better than the others under certain conditions. To further examine the characteristics and trends of each method, we aggregated the number of CCIs detected in three datasets in which each of the three methods excelled (Figure 4).

**Figure 4.**
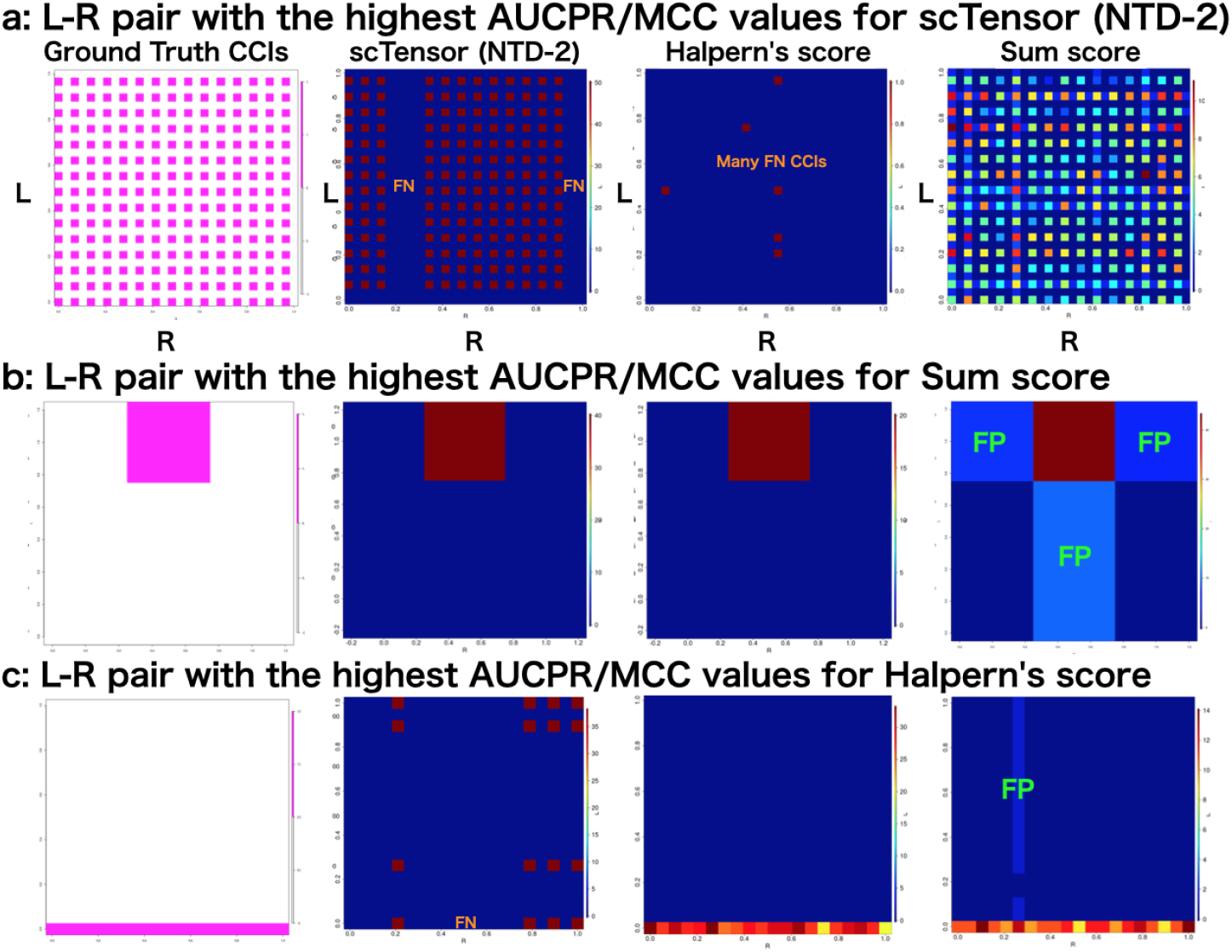
Analyses of three datasets in which each of the three methods excelled. Summary of the number of significant cell–cell interactions (CCIs) with (a) three cell types, one CCI types, one-to-one CCI style, and 1st-CCI type; (b) 20 cell types, five CCI types, many-to-many CCI style, 2nd-CCI type; and (c) 30 cell types, five CCI types, many-to-many CCI style, 5th-CCI type. The *y*-axis (L) and *x*-axis (R) indicate the ligand-expressing cell types and the receptor-expressing cell types, respectively. FN and FP indicate false negative and false positive CCIs, respectively.

`scTensor` (NTD-2): this method performed well when the CCI style was many-to-many. For example, in Figure 4a, most many-to-many CCIs could be detected. Although there were some false negative (FN) CCIs that were not detected, there were fewer false positive (FP) CCIs. In contrast, `Halpern’s score` was too conservative against this dataset and failed to detect most of the CCIs by the label permutation test. At a first glance of Figure 4a, the `sum score` appears to work well with these data, but under scrutiny at the level of individual L-R pairs, `sum score` results contain many FP and FN CCIs (Additional File 13).

`Sum score`: This method performed well when the number of cell types was small and the style of CCI was restricted to one-to-one (Figure 4b). Even though `Halpern’s score` and `scTensor` (NTD-2) were able to detect similar CCIs, `Halpern’s score` was quite conservative and contained many FN CCIs because it considered many CCIs to not be significant. For `sum score`, there seemed to be a bias toward FP CCIs. If the degree of co-expression of an L-R pair is high between two cell types, this method seems to detect FP pairs in which only one of the L-R is highly expressed. In such cases, cross-shaped patterns were observed in the heatmap in Figure 4. In our simulated datasets, this cross-shaped pattern of FP CCIs were observed more frequently in the `sum score`.

`Halpern’s score`: In most data sets, `Halpern’s score` was found to be too conservative, with many FN CCIs, but in a very specific situation, that is, when the CCI-style was one-to-all (or all-to-one), it outperformed the other methods (Figure 4c). In contrast, `scTensor` (NTD-2) inferred many FN CCIs among these data, while the `sum score` identified many FP CCIs (Additional File 13).

#### scTensor could selectively detect many-to-many CCIs in real datasets

Next, we applied these CCI methods to real empirical datasets (Table 1 and Additional Files 3 - 12). As expected from the results of simulated datasets, `scTensor` (NTD-2) outperformed the other methods on these real datasets, which contain many-to-many CCIs. Regarding AUCPR (Figure 5a) and MCC (Figure 5b) values, `scTensor` (NTD-2) achieved higher values compared with the other methods, although the difficulty of detecting the CCIs was highly dependent on the dataset (Additional Files 4 and 8). We further investigated the real empirical datasets and found that the known CCIs reported by the original papers were reproduced by `scTensor` (Table 2). Additionally, some predicted many-to-many CCIs can be considered biologically plausible because the CCIs are related to the same signaling pathways of known CCIs, although the original papers did not refer to the CCIs. These results can be interactively investigated using the HTML report generated by `scTensor` (Additional Files 13 - 17).

**Table 1.** Empirical datasets subjected to cell–cell interaction (CCI) identification.

| Name | Organisms | GEO ID | # Genes | # Cells | # Cell types | CCI-tensor size |
| --- | --- | --- | --- | --- | --- | --- |
| FetalKidney [36] | *Homo sapiens* | GSE109205 | 3204 | 4131 | 11 | 11×11×1072 |
| GermlineFemale [25] | *Homo sapiens* | GSE86146 | 2717 | 992 | 8 | 8×8×1622 |
| HeadandNeckCancer [54] | *Homo sapiens* | GSE103322 | 5244 | 5577 | 26 | 26×26×61 |
| Uterus [33] | *Mus musculus* | GSE118180 | 2566 | 6443 | 12 | 12×12×2282 |
| VisualCortex [48] | *Mus musculus* | GSE102827 | 5365 | 47209 | 30 | 30×30×274 |

**Table 2.**
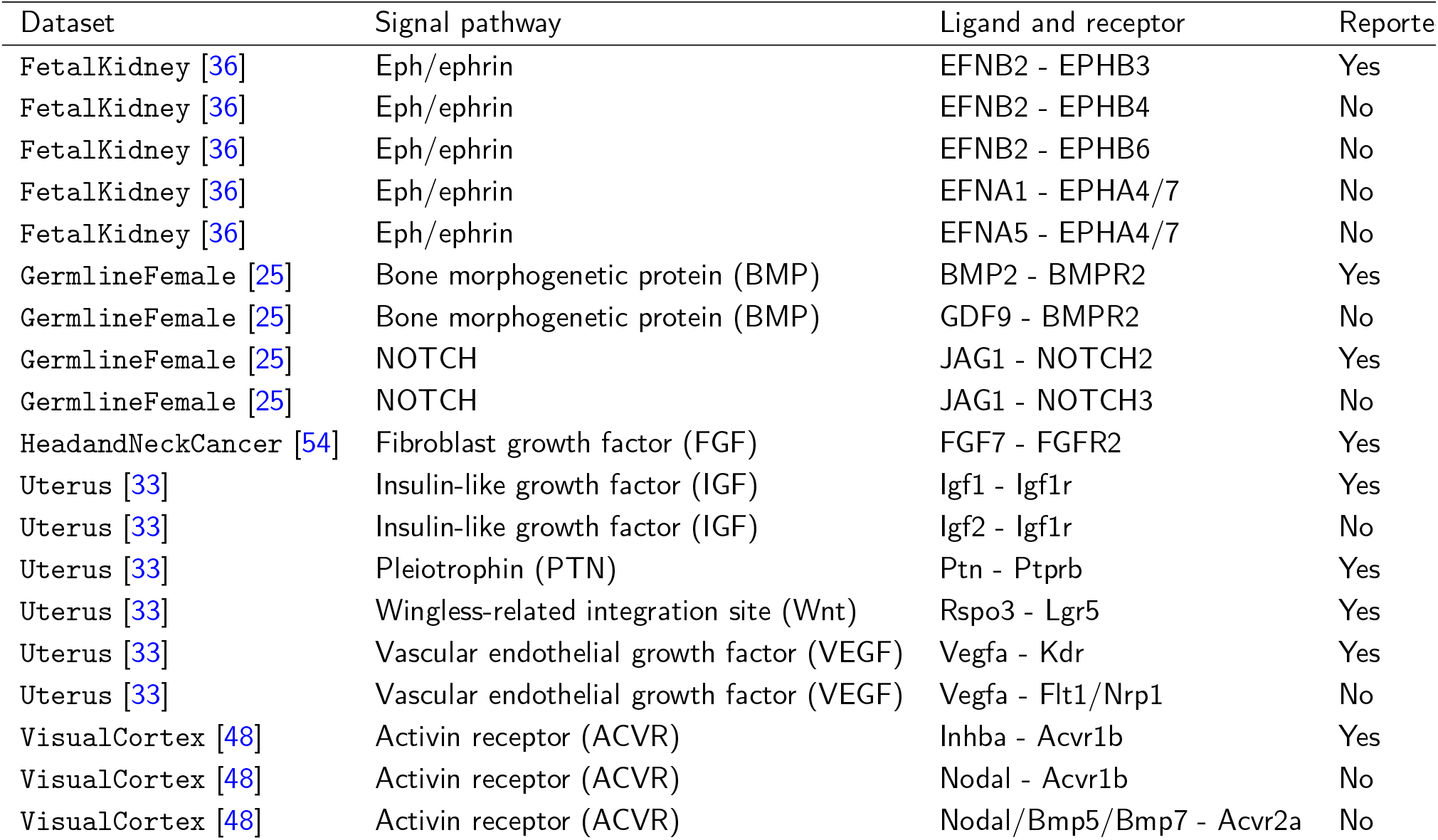
Many-to-many cell–cell interactions (CCIs) detected by only `scTensor`.

**Figure 5.**
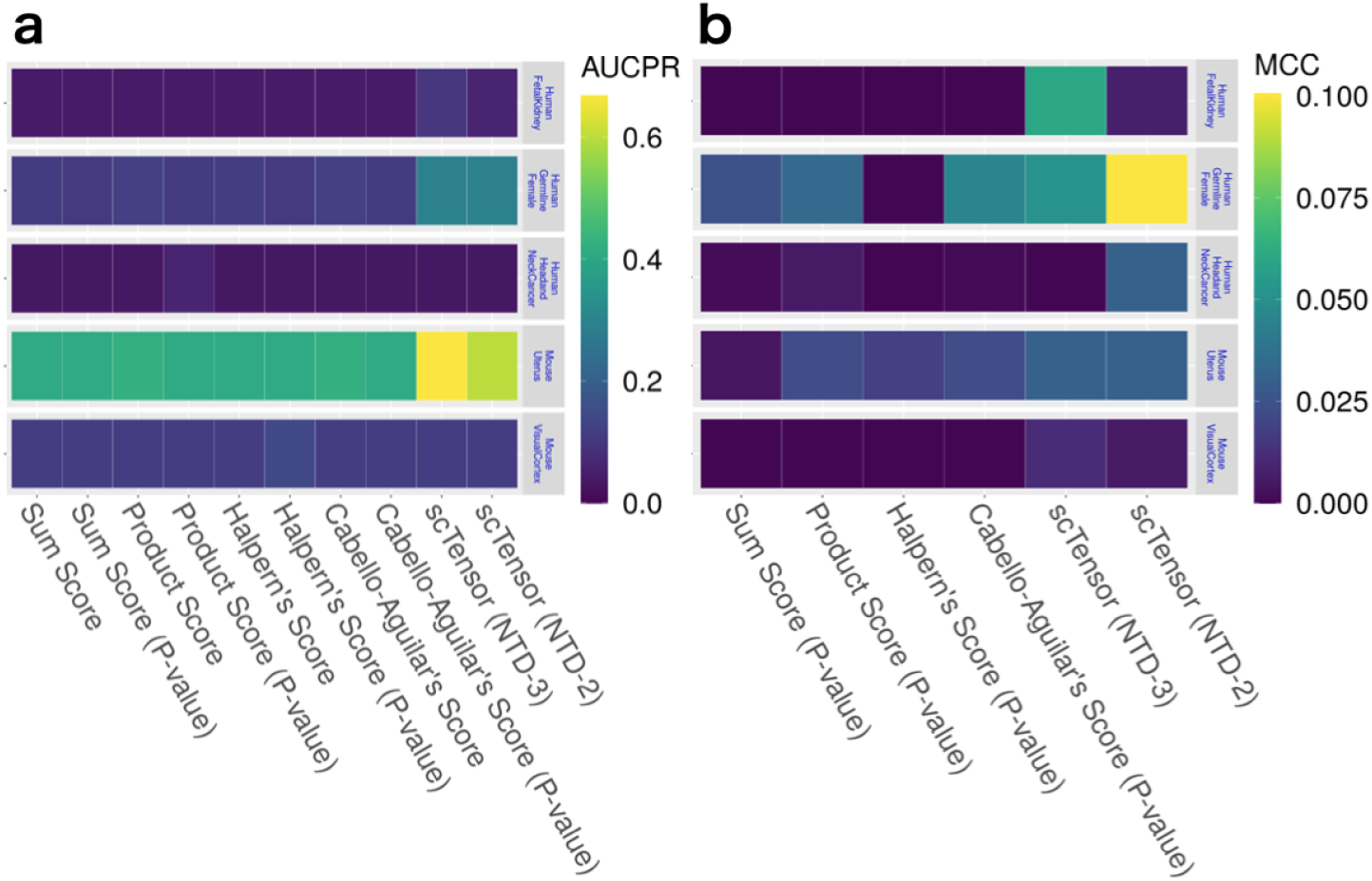
Cell–cell interaction (CCI) identification results from real empirical datasets. (a) Area under the curve of precision-recall (AUCPR) of all the methods. (b) Matthews correlation coefficient (MCC) of the binarization methods.

#### Computational complexity and memory usage

We also assessed the orders of computational complexity and memory usage of all the CCI methods (Table 3). All the L-R score methods require 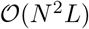 order both in the computation and in memory usage, where *N* is the number of cell types and *L* is the number of L-R pairs. The label permutation tests combining any L-R scores require 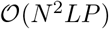 in computation, where *P* is the number of random shuffles of cell-type labels. In many cases, *P* is typically set as a large value greater than 1,000 [37, 69], making this a very time-consuming calculation.

**Table 3.** Order of calculation time and memory space for cell-cell interaction (CCI) identification.

| Method | Calculation time | Memory space |
| --- | --- | --- |
| Sum/product score | $O(N^2L)$ | $O(N^2L)$ |
| Permutation test of sum/product score | $O(N^2LP)$ | $O(N^2L)$ |
| Halpern's score | $O(N^2L)$ | $O(N^2L)$ |
| Permutation test of Halpern's score | $O(N^2LP)$ | $O(N^2L)$ |
| Cabello–Aguilar's score | $O(N^2L)$ | $O(N^2L)$ |
| Permutation test of Cabello–Aguilar's score | $O(N^2LP)$ | $O(N^2L)$ |
| Previous scTensor (NTD-3) | $O(N^2L(R1 + R2 + R3))$ | $O(N^2L)$ |
| scTensor (NTD-2) | $O(N^2L(R1 + R2))$ | $O(N^2L)$ |

In contrast, `scTensor` (NTD-2) does not perform the label permutation; instead, it simply utilizes the factor matrices after the decomposition of the CCI-tensor. Hence, the order of computational complexity is reduced to 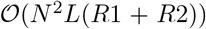, where *R*1 and *R*2 are the number of columns or “rank” parameters for the first- and second-factor matrices, respectively. The rank parameters are typically set as small numbers (e.g., 10), this leads to a substantial computational advantage compared with the label permutation test. The computation time and memory usage when analyzing the simulated and real empirical datasets show that `scTensor` has an advantage in computational complexity compared with the label permutation test (Additional Files 5 and 6).

### Implementations

`scTensor` is implemented as an R/Bioconductor package that is freely available. Both a scRNA-seq dataset and L-R database are required for `scTensor` execution. The default format for a scRNA-seq dataset is `SingleCellExperiment`, in which the gene IDs correspond to NCBI’s Gene database to allow links with other databases (Figure 6a). A scRNA-seq dataset can also be converted from a `Seurat` object. We provided instructions for this data conversion (https://bioconductor.org/packages/release/bioc/vignettes/scTensor/inst/doc/scTensor_1_Data_format_ID_Conversion.html#case-iii-umi-count).

**Figure 6.**
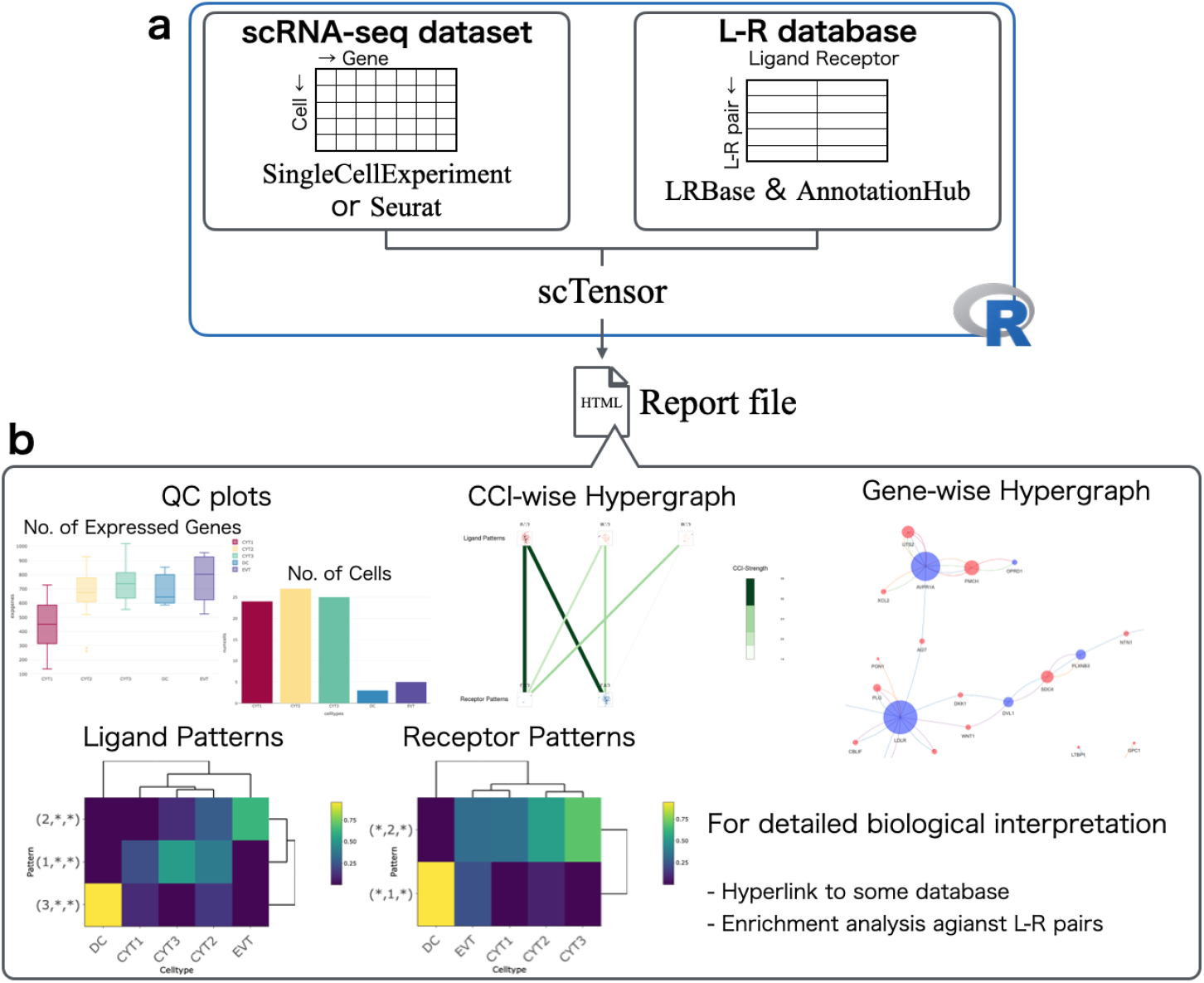
Implementation of the scTensor package. (a) `scTensor` is an R package that requires the input of both an scRNA-Seq expression matrix (`SingleCellExperiment` or Seurat) and a ligand–receptor (L-R) database (`LRBase`). The `LRBase` is retrieved from the `AnnotationHub` remote server, after which a `LRBase` object is created. (b) Using these objects, `scTensor` generates an HTML report file, and the results of cell–cell interaction (CCI) analysis can be visualized with a wide variety of plots.

`LRBase`, which is the L-R database for `scTensor`, is stored on a remote server called `AnnotationHub` and is downloaded to the user’s machine on demand, only when called by the user (Figure 6a). To extend out method to a wide range of organisms, in this work, we originally constructed and are providing the L-R lists for 125 organisms (https://github.com/rikenbit/lrbase-workflow/blob/master/samplesheet/samplesheet.csv). The details of the data processing pipeline are summarized in the README.md of lrbase-workflow (https://github.com/rikenbit/lrbase-workflow), which is a workflow for constructing the `LRBase` for each of the species. For data sustainability, we offer the data files, including older versions, on the `AnnotationHub` server. The data files are bi-annually updated in conjunction with Bioconductor updates and are provided using lrbase-workflow. Users can specify which version of the data is used for analysis, thus ensuring data reproducibility.

NTD-2 was implemented as within the function of `nnTensor` [71] R/CRAN package and internally imported into `scTensor`. `scTensor` constructs the CCI-tensor, decomposes the tensor by the NTD-2 algorithm, and generates an HTML report.

To enhance the biological interpretation of `CaH` results, we implemented some visualization functions (Figure 6a) and these plots can be interactively investigated via web browser. A wide variety of gene-wise information is included in the report and can be linked to the L-R lists through the use of other R/Bioconductor packages; the gene annotation is assigned by `biomaRt` [72] (Gene Name, Description, Gene Ontology [GO], STRING, and UniProtKB), `reactome.db` [73] (Reactome) and `MeSH.XXX.eg.db` [74] (Medical Subject Headings [MeSH]), while the enrichment analysis (also known as over-representative analysis [ORA]) is performed by `GOstats` [75] (GO-ORA), `meshr` [74] (MeSH-ORA), `ReactomePA` [76] (Reactome-ORA), and `DOSE` [77] (Disease Ontology (DO)-ORA, Network of Cancer Genes (NCG)-ORA, DisGeNET-ORA).

To validate that the detected the co-expression of L-R gene pairs is also consistently detected in the other data including tissue- or cell-type-level transcriptome data, the hyperlinks to RefEx [78], Expression Atlas [79], SingleCell Expression Atlas [80], scRNASeqDB [81], and PanglaoDB [82] are embedded in the HTML report, facilitating comparisons of the L-R expression results with the data from large-scale genomics projects such as GTEx [83], FANTOM5 [84], the NIH Epigenomics Roadmap [85], ENCODE [86], and the Human Protein Atlas [87]. Additionally, in consideration of users who might want to experimentally investigate detected CCIs, we embedded hyperlinks to Connectivity Map (CMap [88]), which provides relationships between perturbations by the addition of particular chemical compounds/genetic reagents and the resulting gene expression change.

## Discussion

In this work, we regarded CCIs as `CaH`s, which represent the triadic relationships of ligand-expressing cell types, receptor-expressing cell types, and the related L-R pairs. We implemented a novel algorithm `scTensor` based on a tensor decomposition algorithm for detecting such `CaH`s. Our evaluations using both simulated and real empirical datasets suggest that `scTensor` can detect many-to-many CCIs more accurately than the other conventional CCI methods. Additionally, the calculation time and memory usage performances of `scTensor` are also superior to those of the other CCI methods.

To extend the use of `scTensor` to a wide range of organisms, we also created multiple L-R datasets for 125 organisms. `scTensor` has been published as an R/Bioconductor package, facilitating the reproducibility of data analysis and the maintainability of datasets. We also implemented an HTML report function that simplifies checking the analysis results of `scTensor`. Like many CCI tools, `scTensor` can import an external L-R database.

In the development of many CCI tools, the authors also develop their own L-R databases and investigate the differences among various L-R databases, particulaly when comparing their method with other conventional methods [89]. This makes it difficult to distinguish whether the performance of a method is caused by differences in algorithms or databases. Although the primary CCI resources used for existing L-R tools are highly duplicated, even slight differences can influence the detection of CCIs [89]. Therefore, to separate these two comparisons and to focus only on the algorithmic differences, in this work, we compared several existing CCI algorithms, by re-implementing them and anchoring them to a common L-R database.

We were also able to examine several strengths and weaknesses of the methods other than `scTensor`. For example, `Halpern’s score` was found to be too conservative with many FN CCIs, but it was superior to the other methods with respect to the detection of one-to-all (or all-to-one) CCIs. A possible reason for this is that since the formula for this score includes the square root of the chi-square distribution with two degree of freedom (or an exponential distribution with an expected value of 2), and these distributions are known to be heavy-tailed to some extent, thus potentially inflating the number of significant L-R pairs.

The permutation test implicitly assumes that the interactions occur between very few cell types because the larger the observed L-R score is than the empirical distribution computed by label permutation, the more significant the test result is. However, if the expression levels of ligand and receptor genes are high in any cell of any cell type, the L-R scores calculated by the label permutation are will also be high, and thus, the observed value of the L-R score will be regarded as not a particularly high value in the empirical distribution; consequently, such a test result will be not significant. Therefore, detection of many-to-many CCIs by label permutation test is difficult in principle. In the extreme case of all-to-all CCIs, the current approaches (although it also includes `scTensor`) cannot avoid FN CCIs.

There are still some plans to improve `scTensor` to build on the advantages of this current framework. For example, the algorithm can be improved by utilizing acceleration techniques such as randomized algorithm/sketching methods [90], incremental algorithm/stochastic optimization [91, 92], or distributed computing on large-scale memory machines [93] for tensor decomposition, as is now available.

To reduce the memory usage of `scTensor`, we are developing `DelayedTensor` [94], which is an R/Bioconductor package to perform various tensor arithmetic and tensor decomposition algorithms based on `DelayedArray` [95], another R/Bioconductor package for handling out-of-core multidimensional arrays in R. We intend to reduce the memory usage of `scTensor` by supporting this data format.

Tensor data formats are very flexible ways to represent heterogeneous biological data structures [96], because they easily integrate supplemental information about genes or cell types in a semi-supervised manner. Such information could extend the scope of the data and thus improve the accuracy of inferences. For example, there are some attempts to use the following additional information for CCI detection as well (for more details, see Additional File 1).

- CCI inference via receptor-receptor and extracellular matrix data [51, 97].
- Consideration of multi-subunit complexes [37].
- Comparison of CCIs across multiple conditions [62, 98–105].
- Consideration of downstream transcriptional factors, target genes, and signaling pathways [69, 106–108].
- Integration with bulk RNA-Seq or other type of omics datasets [37, 64, 65, 106–109].
- Integration with pseudo-time [67, 110, 111].
- Integration with spatial transcriptome data [112].

In particular, in a recent benchmark study [112], the proximity of spatial coordinates on tissue sections measured by spatial transcriptome technology and the CCI detected by L-R data were correlated, and some studies have attempted to integrate these two kinds of datasets a single model ([112] and Additional File 1). Auxiliary Information such as the proximity in spatial coordinates can be incorporated as a regularization term to extend the tensor decomposition model [113–115].

Although it is beyond the scope of the present paper to cover all of the above-mentioned topics, considering these in the framework of tensor decomposition is a promising research direction, so we aim to continuously work on these through the development of updates and releases of `scTensor` for Bioconductor.

## Conclusion

In this work, we present and evaluate `scTensor`, a new method for detecting CCIs based on L-R co-expression in scRNA-seq datasets. We also revealed that the widely used label permutation test has a bias that impedes the detection of many-to-many CCIs and demonstrated that the proposed method is a viable alternative.

## Materials and methods

### Simulated datasets

The simulated single-cell gene expression data were sampled from the negative binomial distribution *NB* (*f_gc_m_g_, ϕ_g_*), where *f_gc_* is the fold-change (FC) for gene *g* and cell type *c*, and *m_g_* and *ϕ_g_* are the average gene expression and the dispersion parameter of the expression of gene *g*, respectively.

The *m_g_* value and gene-wise variance *v_g_* were calculated from a real scRNA-seq dataset of mouse embryonic stem cells (mESCs) measured by Quartz-Seq [116], and the gene-wise dispersion parameter *ϕ_g_* was estimated as 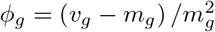.

For the determination of differentially expressed genes (DEGs) and non-DEGs, *f_gc_* values were calculated based on the non-linear relationship of FC and the gene expression level log_10_*f_gc_* = *a* exp(−*b* log_10_ (*m_g_* + 1)). To estimate the parameters *a* and *b*, we detected the DEGs using `edgeR`. By setting the threshold values (i.e., false discovery rate) of `edgeR` as 10^−2^ (E2), 10^−5^ (E5), and 10^−10^ (E10) and using the resulting DEGs, *a* and *b* values for each threshold were estimated as (0.701, 0.363), (1.907, 0.666), and (4.429, 0.814), respectively.

For genes identified as DEGs based on a threshold according to the non-linear relationship above, the estimated *f_gc_* value was used, otherwise, 1 is specified as *f_gc_*. If a ligand gene of a cell type and a receptor gene of a cell type were both DEGs, we defined the relationship between these cell types as the ground truth CCIs and used them for quantitative evaluation.

To simulate the “dropout” phenomenon of scRNA-seq experiments, we also introduced the dropout probability 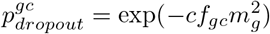, which is used in `ZIFA` [117] (default, *c*=1), and the expression values were randomly converted to 0 according to the dropout probability.

To simulate various situations, we set many different CCI tensors, considering the number of cell types (3, 5, 10, 20, or 30), the style of CCIs (one-to-one or many-to-many), the number of types of CCIs (1, 3, or 5), and the DEG threshold value (E2, E5, or E10); in total, 90 synthetic CCI tensors were generated.

### scRNA-seq real datasets

The gene expression matrix of human `FetalKidney` data was retrieved from the GEO database (GSE109205), and only highly variable genes (HVGs : http://pklab.med.harvard.edu/scw2014/subpoptutorial.html) with low *P*-values (≤ 1E-1) were extracted. The cell-type label data were provided by the authors upon our request.

The gene expression matrix of human `GermlineFemale` data was retrieved from the GEO database (GSE86146), and only HVGs with low *P*-values (≤ 1E-7) were extracted.

The gene expression matrix and the cell-type labels of human `HeadandNeckCancer` data were retrieved from the GEO database (GSE103322), and only HVGs with low *P*-values (≤ 1E-1) were extracted.

The gene expression matrix of mouse `Uterus` data was retrieved from the GEO database (GSE118180), and only HVGs with low *P*-values (≤ 1E-1) were extracted. The cell-type labels were provided by the authors upon our request.

The gene expression matrix and the cell-type labels of mouse `VisualCortex` data were retrieved from the GEO database (GSE102827), and only HVGs with low *P*-values (≤ 1E-1) were extracted.

The gene expression values of each cell are normalized by CPMED [56–58] and logarithm transformation, for variance-stabilization, is performed to the data matrix. For analyzing these real datasets, known L-R pairs in DLRP [118], IUPHAR [119], and HPMR [120] were searched in the data matrix. We defined the ground truth CCIs between two cell types if the CCIs were reported by the original studies. The L-R pairs associated with the CCIs were used for quantitative evaluation.

### scTensor algorithm

#### CCI-tensor Construction

Here, data matrix ***Y*** ∈ ℝ^*I×H*^ is the gene expression matrix of scRNA-seq data, where *I* is the number of genes and *H* is the number of cells. Matrix ***Y*** is converted into cell-type-wise average matrix ***X*** ∈ ℝ^*I×J*^, where *J* is the number of cell types. The cell-type labels are assumed to be specified by the user’s prior analysis, such as clustering or confirmation of marker gene expression. The relationship between the ***X*** and ***Y*** is described below:

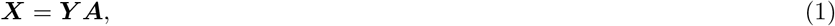

where the matrix ***A*** ∈ ℝ^*H×J*^ converts cellular-level matrix ***Y*** to cell-type-level matrix *X* and each element of ***A*** is

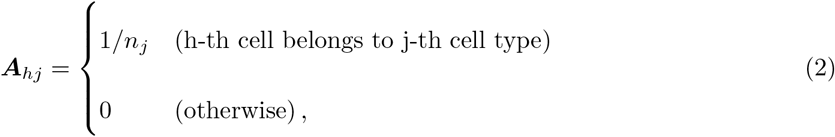

where *n_j_* is the number of cells belonging to the *j*’s cell type.

Next, we search to determine whether any L-R pairs stored in the L-R database are both in the row names of matrix *X*, and if both IDs are found, corresponding *J*-length row-vectors of the ligand and receptor genes (***x**_L_* and ***x**_R_*) are extracted.

Finally, a *J* × *J* matrix is calculated as the outer product of ***x**_L_* and ***x**_R_* and incrementally stored. The stacked *J* × *J* matrices can be considered as a three-dimensional array, which is also known as a three-order tensor. The following outer product in the *k*-th L-R pair (*L* (*k*) and *R* (*k*)) found is stored as the frontal slice (sub-tensor) of the CCI-tensor *χ* ∈ ℝ^*J×J×K*^ :

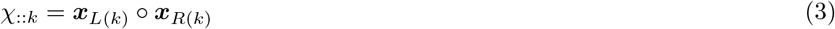

#### Non-negative Tucker3 Decomposition (NTD-3)

To extract the `CaH`s from the CCI-tensor *χ* ∈ ℝ^*J×J×K*^, we utilize NTD-3 and NTD-2, which are generalizations of non-negative matrix factorization (NMF) to tensor data [59, 60]. The NMF approximates a non-negative matrix data as the product of two lower rank non-negative matrices (also known as factor matrices). Similar to NMF, NTD-3 and NTD-2 approximate a non-negative tensor data as the product of some factor matrices and a core tensor.

To extend NMF to NTD-3, we consider iterative updating 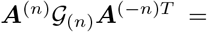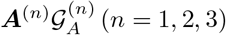, which is the matricized expression of tensor decomposition. Here ***A***^(*−n*)^ is Kronecker product of the factor matrices without ***A***^(*n*)^ and 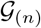 is the mode-*n* matricization of the core tensor 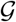. For example, if *n* = 1, these become ***A***^(2)^ ⊗ ***A***^(3)^ and 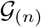, respectively. By replacing *X* in the multiplicative update rule [60] with 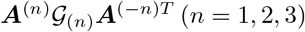, we can obtain the update rule for ***A***^(*n*)^ as follows;

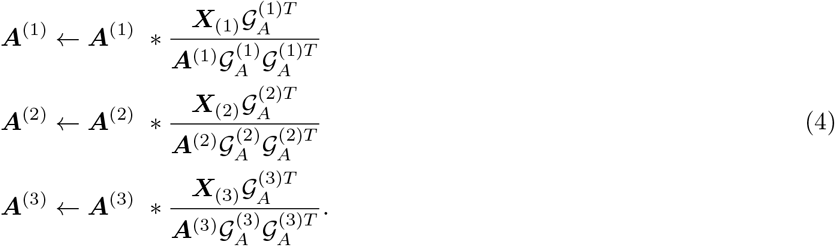

Similarly, the updating rule for core tensor 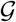 is:

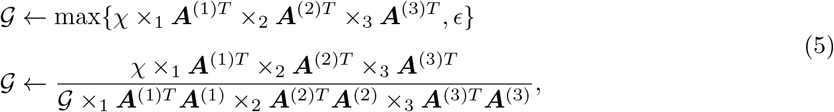

where *ϵ* is a small value included to avoid generating negative values (default value 1E-10).

#### Non-negative Tucker2 Decomposition (NTD-2)

The NTD-3 has three rank parameters to be estimated, and it requires huge search space (*R*1 × *R*2 × *R*3). Additionally, the fewer the factor matrices, the more interpretable the results are. For these reasons, we further expanded the NTD-3 into a model called the NTD-2 [60] since v1.4.0 of `scTensor`.

In NTD-2, the third factor matrix ***A***^(3)^, which is related to L-R pairs, is replaced by an identity matrix *I_K_*, where the *K* diagonal elements are all 1 and the iteration step of ***A***^(3)^ is skipped as follows:

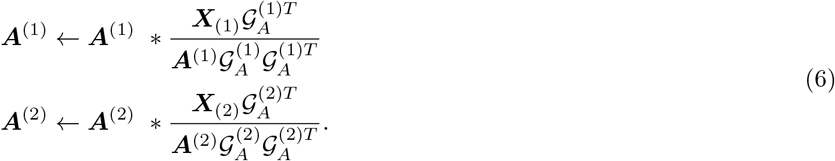

Here, 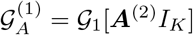 and 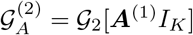.

The updating rule for core tensor 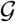 is

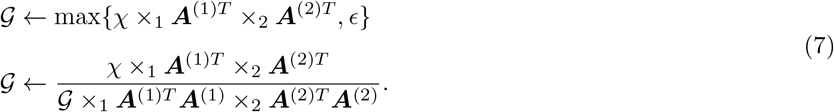

#### Rank estimation of NTD-2

To extract the CaHs, `scTensor` estimates the NTD-2 ranks for each matricized CCI-tensor (*X*^(*n*)^, *n* = 1 or 2). To be able to focus only on the dimensions that are informative and are not noisy, we used an ad hoc approach for NTD-2 rank estimation.

Because NMF is performed in each matricized CCI-tensor in `scTensor`, we estimated each rank of NMF based on the residual sum of squares (RSS) [121] as

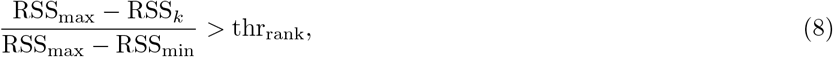

where RSS_max_ is the RSS by full rank NMF, RSS_min_ is the RSS by rank-1 NMF, RSS_k_ is the RSS by rank-*k* NMF, and thr_rank_ is the threshold value, ranging 0 to 1 (the default value is 0.8). RSS by rank-*k* NMF is calculated between a data matrix *X* and the reconstructed matrix from *W* and *H* calculated by multiplicative updating rule [60] as follows:

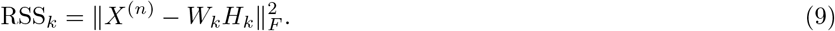

RSS by full-rank and rank-1 NMF is calculated by setting *k* as *J* and 1, respectively. With the estimated ranks 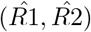, NTD-2 was performed, and only the pairs (r1,r2) with large core tensor values are selected as `CaH`s. In its default mode, `scTensor` selects `CaH`s that explain the top 20 pairs sorted by the core tensor values.

#### Binarization

To binarize each column vector of the factor matrices obtained by NTD-2, median absolute deviation (MAD), which is the median version of standard deviation (SD), was applied. Because we are only interested in the outliers of the elements of each vector in the positive direction, not the negative one, we focused only on the elements that deviate from the median in the positive direction as follows:

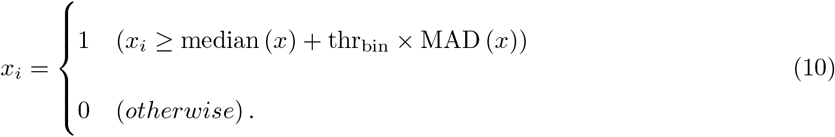

Here, MAD (*x*) is median (∥*x* − median (*x*) ∥) and thr_bin_ is the threshold value (the default value is 1.0).

### L-R scoring

Several methods have been proposed to score the degree of co-expression of a given L-R pair between two cell types, as described below.

#### Sum score

The gene expression of a ligand gene *l* can be averaged over cells belonging to the *s*-th cell type within *J* cell types as follows:

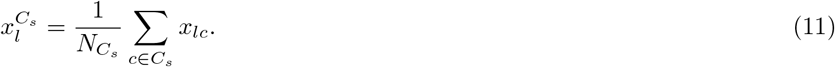

Here, *C_s_* ∈ (*C*_1_, *C*_2_, …, *C_J_*) and 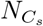 is the number of cells belonging to cell type *C_s_*.

Likewise, the gene expression of a receptor gene *r* is averaged over cells belonging to the *t*-th cell type within *J* cell types as follows:

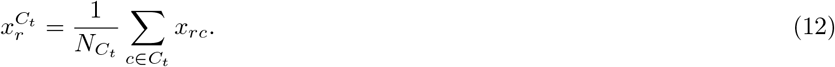

Here, *C_t_* ∈ (*C*_1_, *C*_2_, …, *C_J_*) and 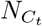 is the number of cells belonging to cell type *C_t_*.

Using these values, the `sum score` is calculated as follows:

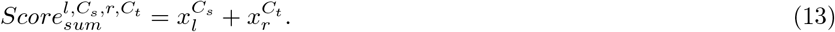

For example, some methods such as CellPhoneDB [37], Giotto [61], CrossTalkR [62], and Squidpy [63], essentially use this type of scoring (Additional File 1).

#### Product score

In some studies, the degree of co-expression is expressed as a product instead of a summation.

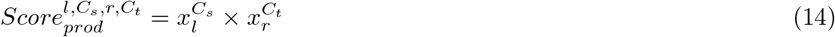

For example, some methods such as NATME [64], FunRes [65], ICELLNET [66], and TraSig [67], essentially use this type of scoring (Additional File 1).

#### Halpern’s score

Derived from the `sum score`, `Halpern et al`. proposed a score described below.

In this score, *Z*-scaling is firstly applied to both 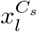 and 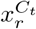 over *J* cell types as follows:

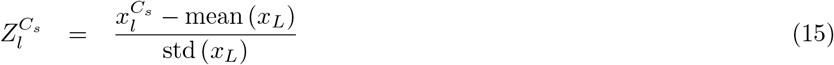

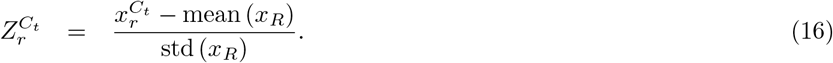

Here, 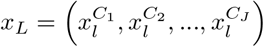 and 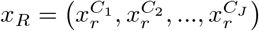. Then, the square root of the sum of squares of these values is used as the degree of co-expression as follows:

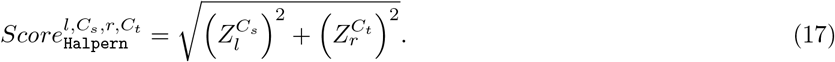

#### Cabello–Aguilar’s score

Derived from the `product score`, `Cabello–Aguilar et al`. proposed a score described as follows:

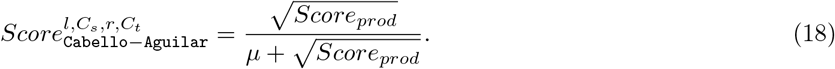

Here, *μ* is the averaged value of the normalized read count matrix and is added as a scaling factor to avoid division by zero. This score is used in SingleCellSignalR [69] and CellTalkDB [70].

### Label permutation method

To quantify the deviation of the observed scores obtained from real data, many studies employ *P*-values in a statistical hypothesis testing framework. Typically, the label permutation method is widely used to calculate *P*-values. In principle, this method can be used in combination with any L-R score as described above.

Here, we consider assigning a *P*-value to any type of 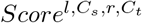 above. In this method, the cluster labels of all the cells are randomly shuffled, and a synthetic score value is calculated. Performing this process 1,000 times generates 1,000 of the values. These values are used to generate the null distribution; for a combination of cell types, the proportion of the means which are “as or more extreme” than the observed mean is calculated as the *P*-value. The label permutation test is performed as a one-tailed test; there is a focus on L-R scores with significantly higher values compared to the null distribution, and not on L-R scores with significantly lower values. Because separating significant CCIs from non-significant CCIs by hypothesis testing can be regarded as a binarization process, label permutation results were compared with the results of `scTensor` binarization.

### Quantitative Evaluation of CCIs

The CCIs detected by the various methods tested in this paper were compared with ground truth CCIs to quantitatively evaluate the performance of each method. To evaluate the results, we used the metrics below.

#### Evaluation of the scoring before and after binarization

Each CCI method uses each corresponding L-R score to quantify the degree of coexpression of a given L-R pair between two cell types. To quantitatively evaluate the performance of each score, area under the curve of receiver operating characteristic (AUCROC) and area under the curve of precision-recall (AUCPR) were used.

A receiver operating characteristic (ROC) curve is a plot of the true positive rate (TPR, or the 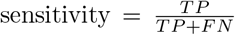) versus the false positive rate (FPR, or 1 - specificity, where 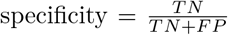) (where TP is the number of true positive CCIs, FP is the number of false positive CCIs, TN is the number of true negative CCIs, and FN is the number of false negative CCIs). The AUCROC value is the area under the ROC curve. AUCROC values range from 0 to 1, and the closer the value is to 1, the more the score indicates enrichment of the ground truch CCIs among the inferred CCIs.

A precision-recall curve is a plot of recall (i.e., sensitivity) versus precision (i.e., 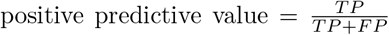). The AUCPR value is the value of the area under the precision-recall curve. AUCPR ranges from 0 to 1, and the closer the value is to 1, the more the score indicates enrichment of the ground truch CCIs among the inferred CCIs. AUCPR is known for its robustness against class imbalance, compared with AUCROC [122–124]. Hence, it seems that AUCPR is more appropriate than AUCROC because the number of significant CCIs are assumed to be less than that of non-significant CCIs in both simulated and real empirical data.

To evaluate whether the binarization was properly performed, we also applied these metrics to assess label permutation. As the label permutation test outputs *P*-values, we utilized 1 - *P*-value to quantify the degree of co-expression of elements of L-R pairs in the test. Because tensor decomposition is an unsupervised learning methods, we cannot distinguish which `CaH`s are enriched within the ground truth CCIs in advance. Additionally, we expected that `scTensor` could separate different styles of CCIs as multiple `CaH`s. Hence, we used the combination of `CaH`s from `scTensor` and ground truth CCIs with the maximum metrics values.

The calculation time and memory usage were evaluated by using the benchmark rules of Snakemake (https://snakemake.readthedocs.io/en/latest/snakefiles/rules.html?highlight=benchmark#benchmark-rules).

#### Evaluation of the scoring after binarization

Each CCI method uses a threshold value (e.g., *P*-value, or MAD for `scTensor`) to differentiate significant CCIs from non-significant CCIs. This process is considered a kind of binarization (1 for significant CCIs, 0 for non-significant CCIs), so we evaluated how well each thresholding strategy could selectively detect the ground truch CCI by comparing the metrics below.

F-measure is the harmonic mean of precision and recall and is defined as follows:

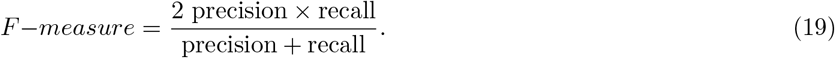

Matthews Correlation Coefficient (MCC) is a special case of Pearson correlation coefficient when two variables are both binary vectors. MCC is defined as follows:

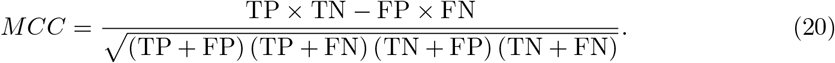

MCC is widely used for binary classification evaluation and especially known for its robustness against the class imbalance, compared with the other metrics such as accuracy, balanced accuracy, bookmaker informedness, markedness, and F-measure [125–127]. Hence, it seems that MCC is more appropriate to use than F-measure because the number of significant CCIs are assumed to be less than that of non-significant CCIs in both simulated and real empirical data.

To distinguish whether the F-measure and MCC values correspond to the number of detected CCIs or their selectivity in focusing only the ground truth CCIs, we also compared the positive rate (PR), false positive rate (FPR), and false negative rate (FNR) values of all the methods.

## Supporting information

https://zenodo.org/record/7412280/files/01_Comparison.xlsx?download=1

https://zenodo.org/record/7412280/files/02_GroundTruth.pdf?download=1

https://zenodo.org/record/7412280/files/03_AUCROC.pdf?download=1

https://zenodo.org/record/7412280/files/04_AUCPR.pdf?download=1

https://zenodo.org/record/7412280/files/05_Memory.pdf?download=1

https://zenodo.org/record/7412280/files/06_Time.pdf?download=1

https://zenodo.org/record/7412280/files/07_F-measure.pdf?download=1

https://zenodo.org/record/7412280/files/08_MCC.pdf?download=1

https://zenodo.org/record/7412280/files/09_PR.pdf?download=1

https://zenodo.org/record/7412280/files/10_FPR.pdf?download=1

https://zenodo.org/record/7412280/files/11_FNR.pdf?download=1

https://zenodo.org/record/7412280/files/12_TR.pdf?download=1

## Availability and requirements

### R packages

- `scTensor`: https://bioconductor.org/packages/devel/bioc/html/scTensor.html
- `nnTensor`: https://cran.r-project.org/web/packages/nnTensor/index.html
- `AnnotationHub`: https://bioconductor.org/packages/release/bioc/html/AnnotationHub.html
- `LRBaseDbi`: https://bioconductor.org/packages/release/bioc/html/LRBaseDbi.html
- Operating system: Linux, Mac OS X, Windows
- Programming language: R (v-4.1.0 or higher), Bioconductor version (v-3.14.0 or higher)
- License: Artistic-2.0
- Any restrictions to use by non-academics: For non-profit use only

### Snakemake workflows

- `scTensor-experiments (for the analyses conducted in this study)`: https://github.com/rikenbit/scTensor-experiments
- `lrbase-workflow (for the bi-annual updates of LRBase)`: https://github.com/rikenbit/lrbase-workflow
- Operating system: Linux, Mac OS X, Windows
- Programming language: Python (v-3.7.8 or higher), Snakemake (v-6.0.5 or higher), Singularity (v-3.8.0 or higher)
- License: MIT
- Any restrictions to use by non-academics: For non-profit use only

## Abbreviations

CCIs: cell–cell interactions
scRNA-seq: single-cell RNA sequencing
L-R: ligand and receptor
CaH: CCI as hypergraph
CMPED: count per median of library size
NTD-3: non-negative Tucker3 decomposition
NTD-2: non-negative Tucker2 decomposition
AUCPR: area under the curve of precision-recall
FPR: false positive rate
FNR: false negative rate
MCC: Matthews correlation coefficient
FP: false positive
FN: false negative
GO: Gene Ontology
MeSH: medical subject headings
ORA: over-representative analysis
DO: Disease Ontology
NCG: Network of Cancer Genes
CMap: Connectivity Map
FC: fold-change
mESCs: mouse embryonic stem cells
DEGs: differentially expressed genes
HVGs: highly variable genes
NMF: non-negative matrix factorization
RSS: residual sum of squares
MAD: median absolute deviation
SD: standard deviation
AUCROC: area under the curve
TPR: true positive rate
TP: true positive
TN: true negative
PR: positive rate

## Competing interests

The authors declare that they have no competing interests.

## Funding

This work was supported by MEXT KAKENHI Grant Number 16K16152 to KT. This work was partially supported by JST CREST grant numbers JPMJCR16G3 and JPMJCR1926 to IN.

## Authors’ contributions

KT and IN designed the study. KT designed the algorithm and benchmark test. KT retrieved and preprocessed the test data to evaluate the proposed method, implemented the source code, and performed all analyses. MI implemented the pipeline for bi-annual automatic updates of the R/Bioconductor packages. All authors have read and approved the manuscript.

## Acknowledgements

Some cell images used in figures are presented by © 2016 DBCLS TogoTV. We thank Mr. Akihiro Matsushima for their assistance with the IT infrastructure for the data analysis. We are also grateful to all members of the Laboratory for Bioinformatics Research, RIKEN Center for Biosystems Dynamics Research for their helpful advice.

## Additional Files

Additional file 1 — List of existing L-R scoring and CCI detection methods (XLSX 16.0 kB, https://zenodo.org/record/7412280/files/01_Comparison.xlsx?download=1)

Additional file 2 — Ground truth CCIs in simulated datasets (PDF 2.0 MB, https://zenodo.org/record/7412280/files/02_GroundTruth.pdf?download=1)

Additional file 3 — AUCROC values of all methods (PDF 3.7 MB, https://zenodo.org/record/7412280/files/03_AUCROC.pdf?download=1)

Additional file 4 — AUCPR values of all methods (PDF 3.8 MB, https://zenodo.org/record/7412280/files/04_AUCPR.pdf?download=1)

Additional file 5 — Memory values of all methods (PDF 3.7 MB, https://zenodo.org/record/7412280/files/05_Memory.pdf?download=1)

Additional file 6 — Computational time values of all methods (PDF 2.7 MB, https://zenodo.org/record/7412280/files/06_Time.pdf?download=1)

Additional file 7 — F-measure values of all binarization methods (PDF 2.6 MB, https://zenodo.org/record/7412280/files/07_F-measure.pdf?download=1)

Additional file 8 — MCC values of all binarization methods (PDF 2.5 MB, https://zenodo.org/record/7412280/files/08_MCC.pdf?download=1)

Additional file 9 — PR values of all binarization methods (PDF 2.4 MB, https://zenodo.org/record/7412280/files/09_PR.pdf?download=1)

Additional file 10 — FPR values of all binarization methods (PDF 2.6 MB, https://zenodo.org/record/7412280/files/10_FPR.pdf?download=1)

Additional file 11 — FNR values of all binarization methods (PDF 2.6 MB, https://zenodo.org/record/7412280/files/11_FNR.pdf?download=1)

Additional file 12 — TR of all datasets (PDF 358.5 kB, https://zenodo.org/record/7412280/files/12_TR.pdf?download=1)

Additional file 13 — Three L-R pairs in which each of the three methods excelled (PNG 237.2 kB, https://zenodo.org/record/7412280/files/13_Crossshaped.png?download=1)

Additional file 14 — HTML report of FetalKidney (ZIP 137.0 MB, https://zenodo.org/record/7412280/files/14_Human_FetalKidney.zip?download=1)

Additional file 15 — HTML report of GermlineFemale (ZIP 173.5 MB, https://zenodo.org/record/7412280/files/15_Human_Germline_Female.zip?download=1)

Additional file 16 — HTML report of HeadandNeckCancer (ZIP 189.6 MB, https://zenodo.org/record/7412280/files/16_Human_HeadandNeckCancer.zip?download=1)

Additional file 17 — HTML report of Uterus (ZIP 243.8 MB, https://zenodo.org/record/7412280/files/17_Mouse_Uterus.zip?download=1)

Additional file 18 — HTML report of VisualCortex (ZIP 319.0 MB, https://zenodo.org/record/7412280/files/18_Mouse_VisualCortex.zip?download=1)

## Notes

### Competing Interest Statement

The authors have declared no competing interest.

