## Supplementary material for "scTensor detects many-to-many cell–cell interactions from single cell RNA-sequencing data": https://zenodo.org/record/7412280/files/02_GroundTruth.pdf?download=1

### Simulated Datasets

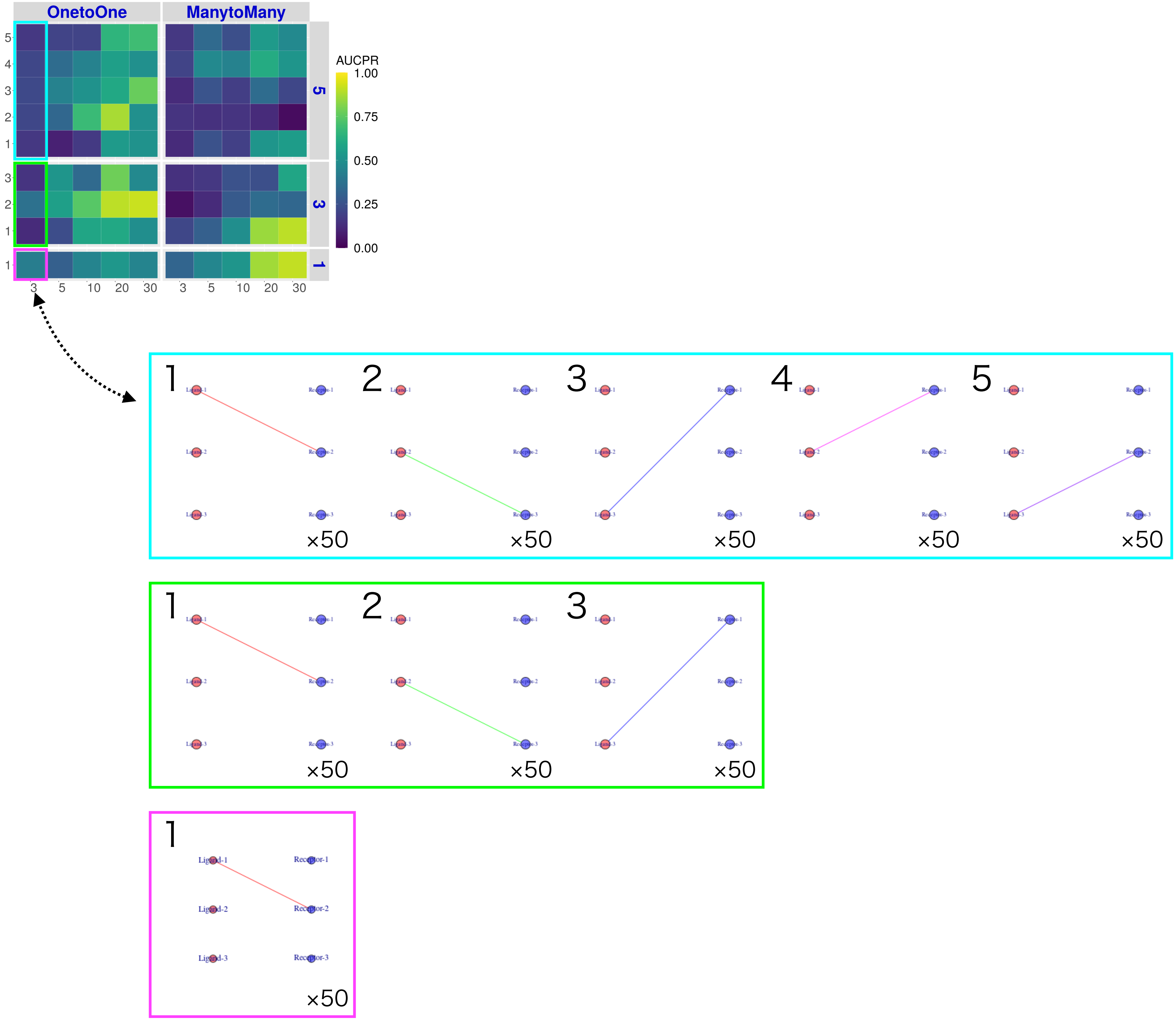

OnetoOne
ManytoMany
AUCPR
1.00
0.75
0.50
0.25
0.00
5
4
3
2
1
3
2
1
1
3 5 10 20 30
3 5 10 20 30
5
3
1
1 2 3 4 5
×50
1 2 3
×50
1
Ligand-1
Ligand-2
Ligand-3
Receptor-1
Receptor-2
Receptor-3
×50

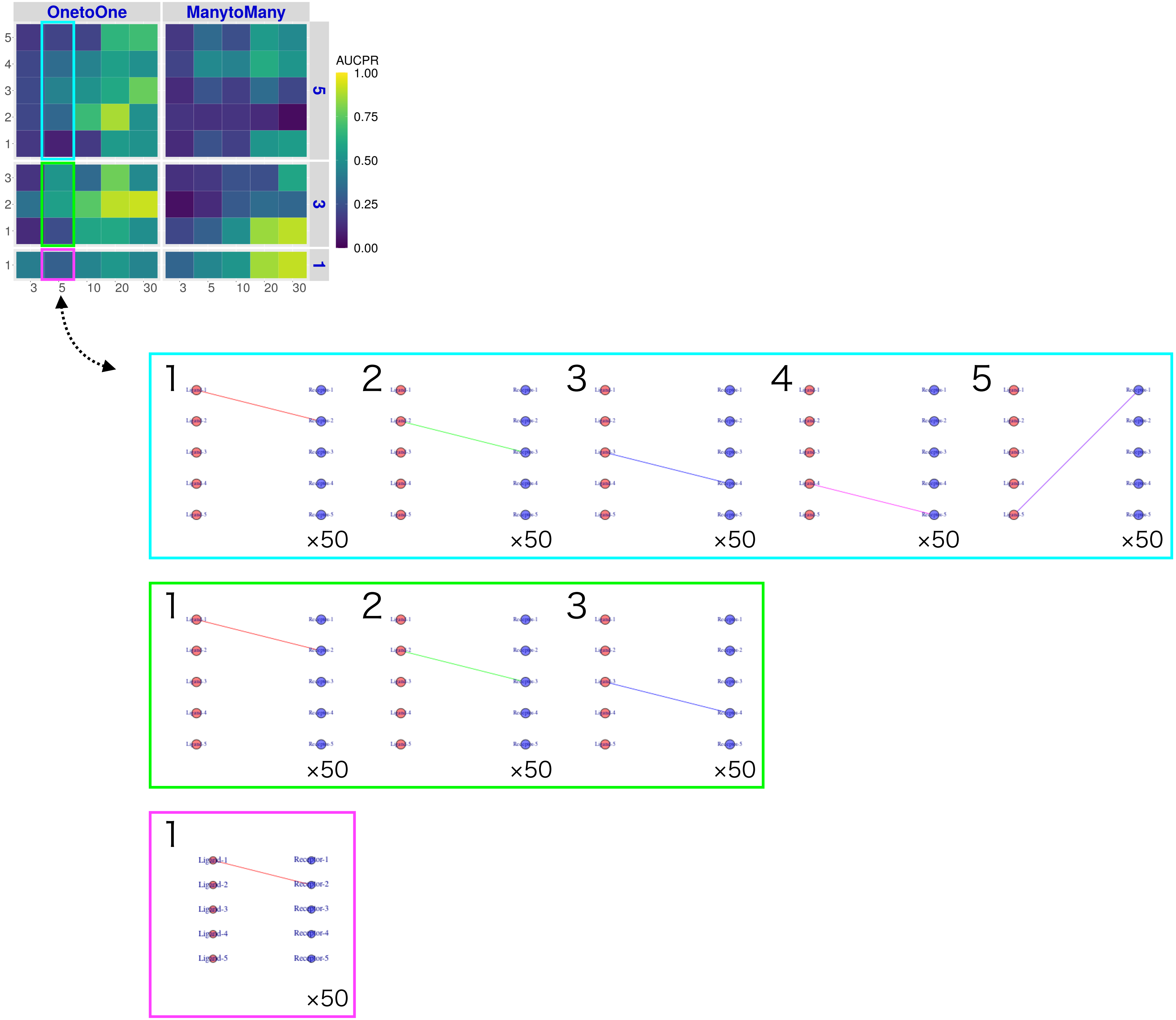
OnetoOne
ManytoMany
AUCPR
1.00
0.75
0.50
0.25
0.00
5
3
1
3
5
10
20
30
×50

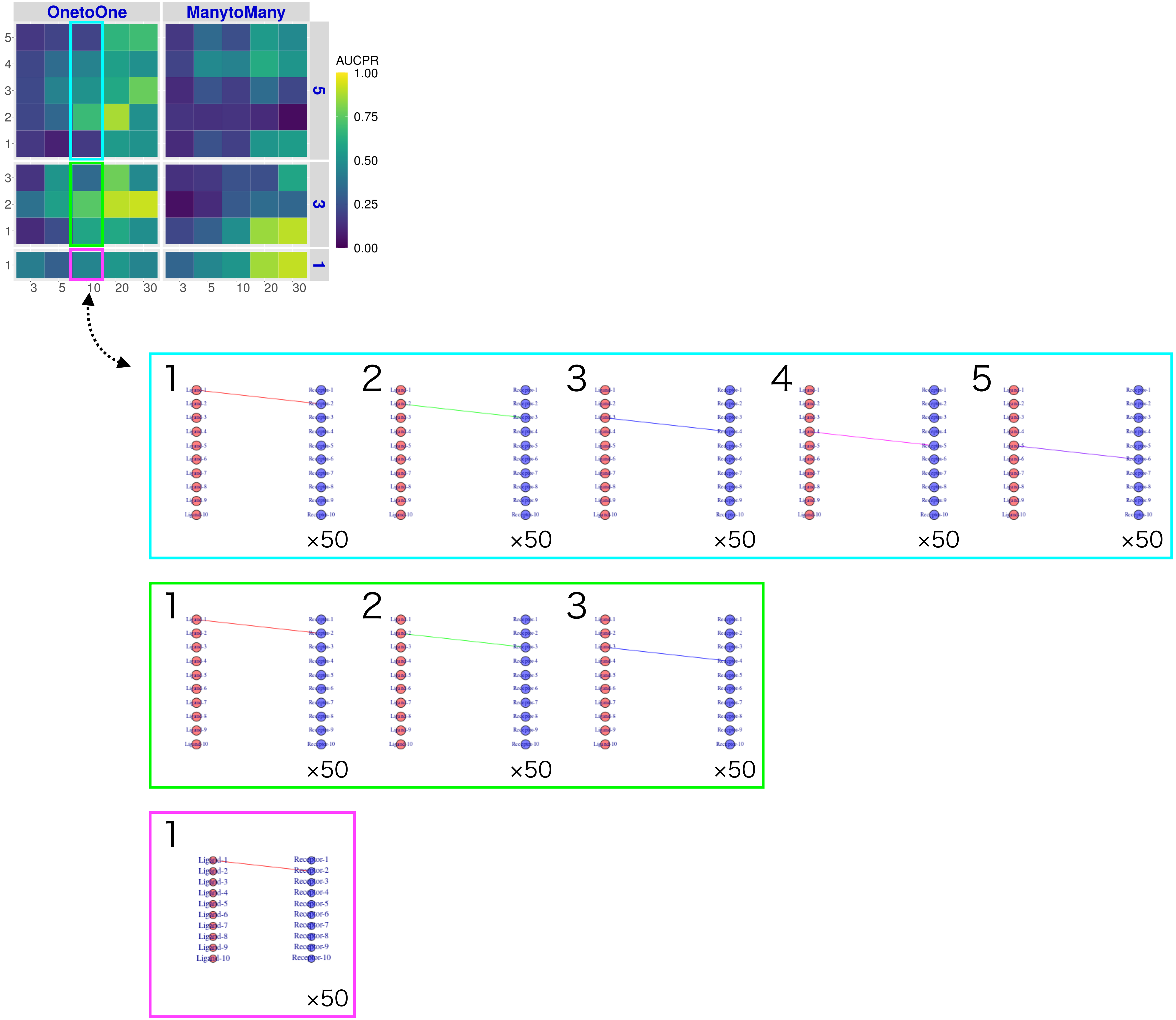
OnetoOne
ManytoMany
AUCPR
1.00
0.75
0.50
0.25
0.00
5
3
1
3
5
10
20
30
×50
Ligand-1
Ligand-2
Ligand-3
Ligand-4
Ligand-5
Ligand-6
Ligand-7
Ligand-8
Ligand-9
Ligand-10
Receptor-1
Receptor-2
Receptor-3
Receptor-4
Receptor-5
Receptor-6
Receptor-7
Receptor-8
Receptor-9
Receptor-10

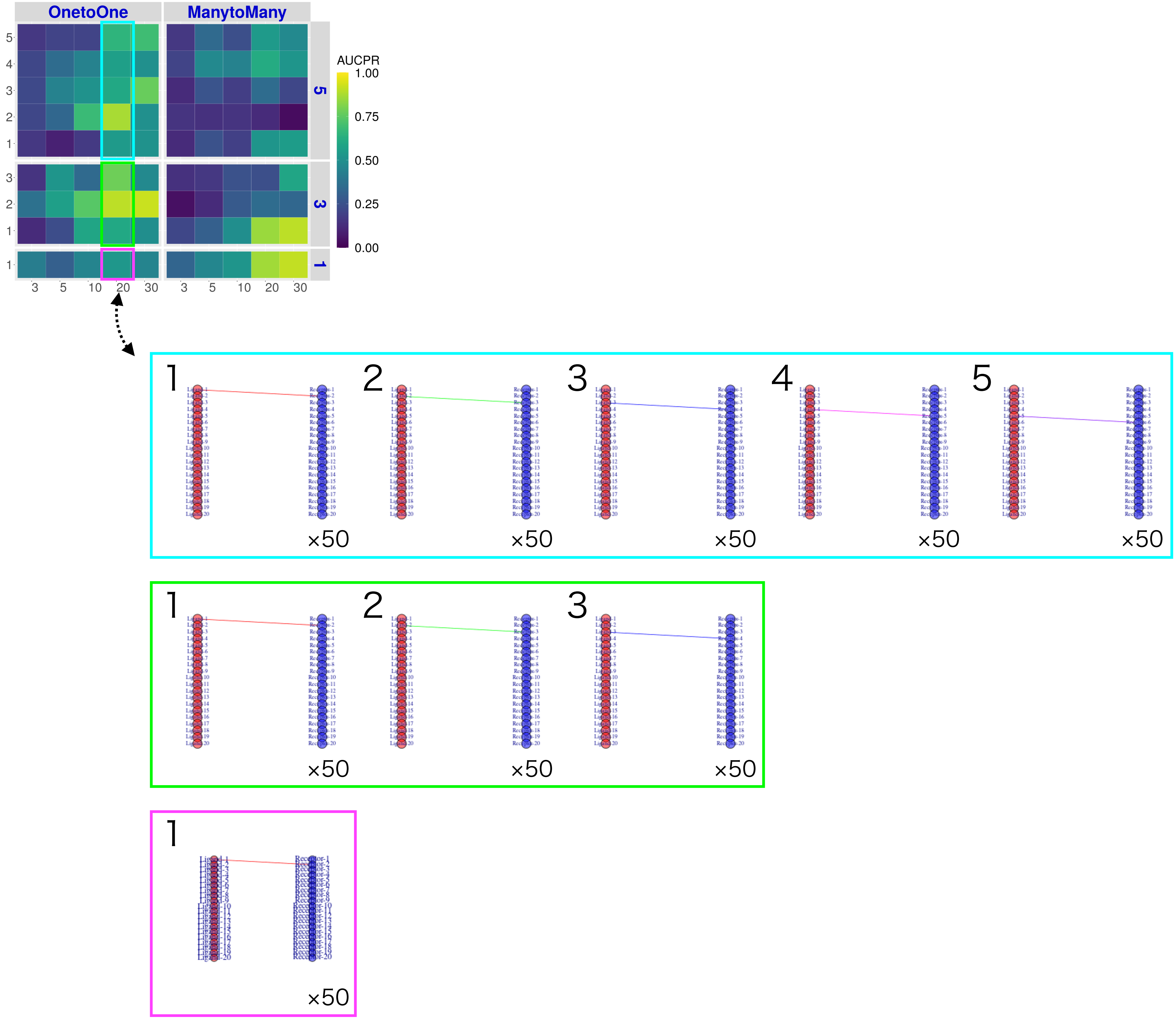

OnetoOne
ManytoMany
AUCPR
1.00
0.75
0.50
0.25
0.00
5
4
3
2
1
3
5
10
20
30
×50

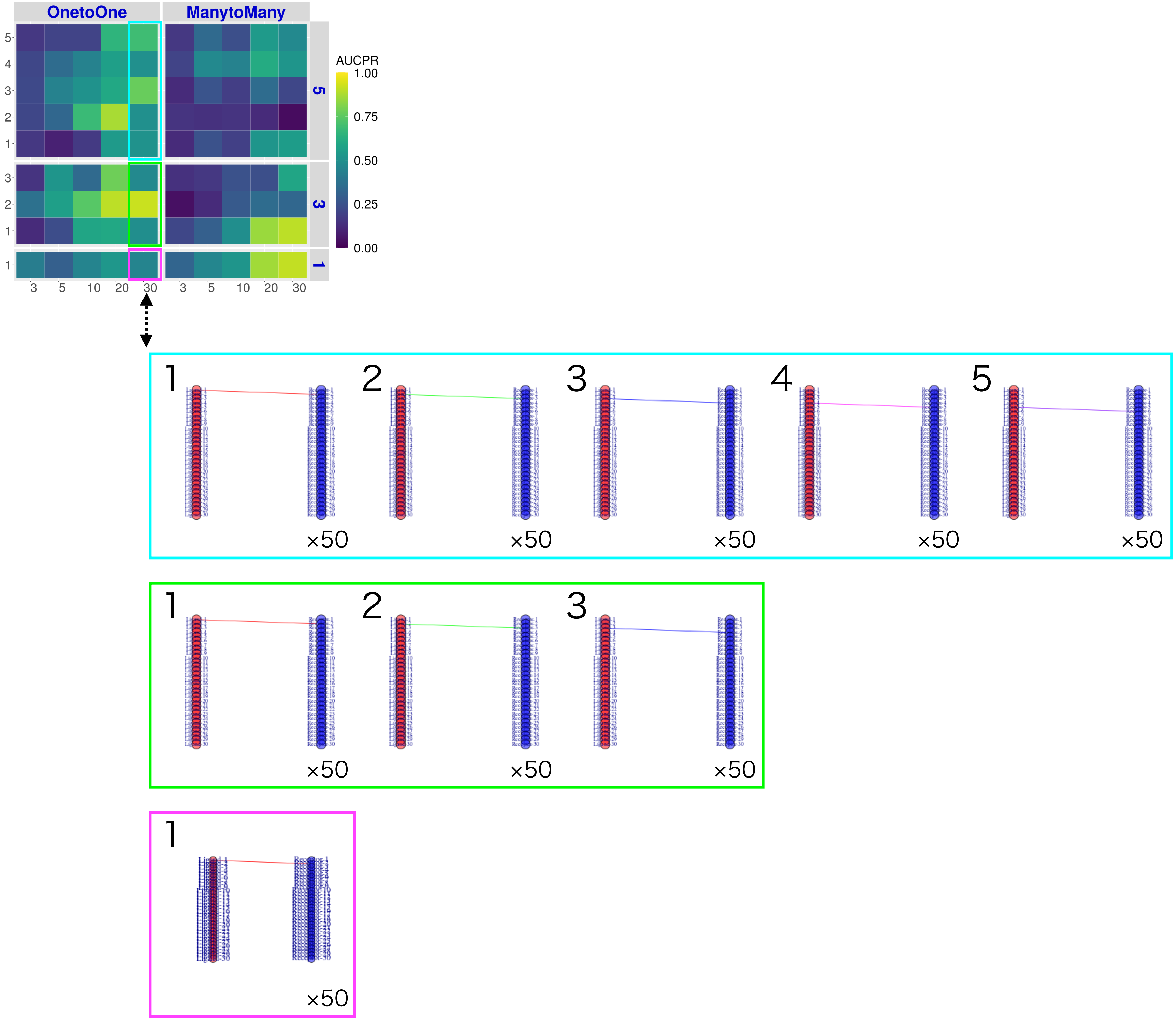
OnetoOne
ManytoMany
AUCPR
1.00
0.75
0.50
0.25
0.00
5
3
1
3 5 10 20 30
×50

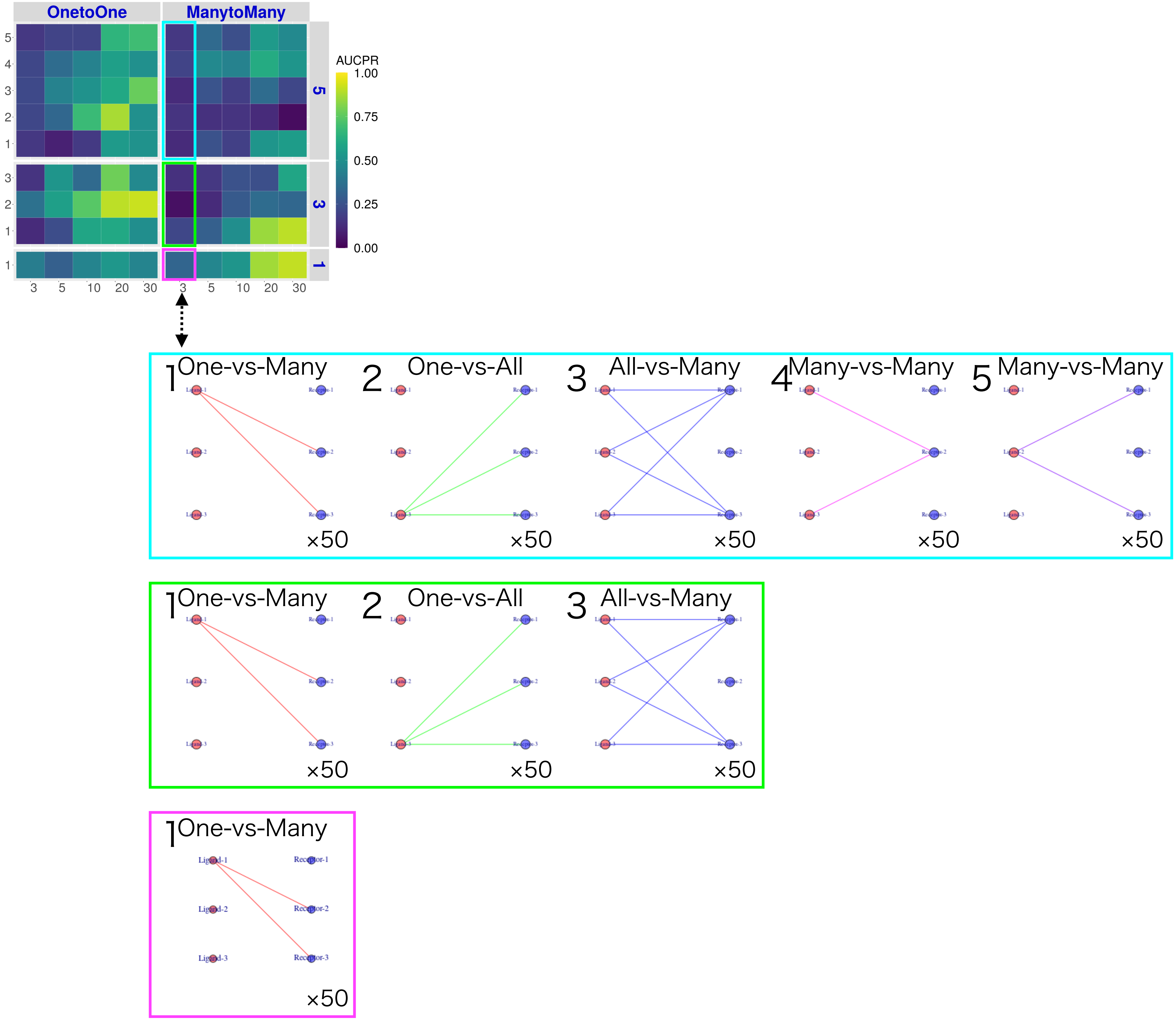
OnetoOne
ManytoMany
AUCPR
1.00
0.75
0.50
0.25
0.00
5
3
1
3
5
10
20
30
1 One-vs-Many
2 One-vs-All
3 All-vs-Many
4 Many-vs-Many
5 Many-vs-Many
×50
Ligand-1
Ligand-2
Ligand-3
Receptor-1
Receptor-2
Receptor-3

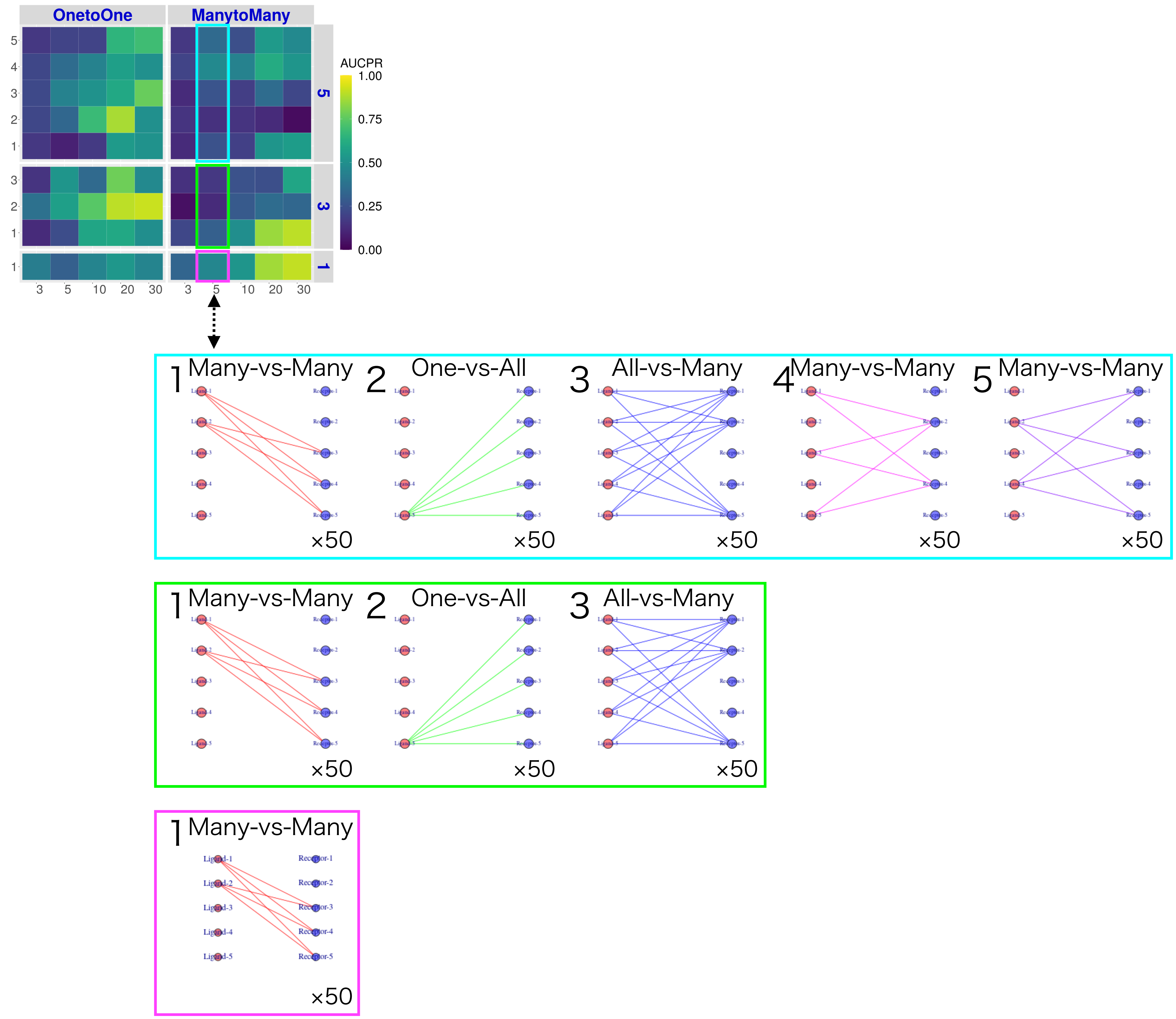
OnetoOne
ManytoMany
AUCPR
1.00
0.75
0.50
0.25
0.00
5
3
1
3 5 10 20 30
3 5 10 20 30
1 Many-vs-Many
2 One-vs-All
3 All-vs-Many
4 Many-vs-Many
5 Many-vs-Many
×50
Ligand-1
Ligand-2
Ligand-3
Ligand-4
Ligand-5
Receptor-1
Receptor-2
Receptor-3
Receptor-4
Receptor-5

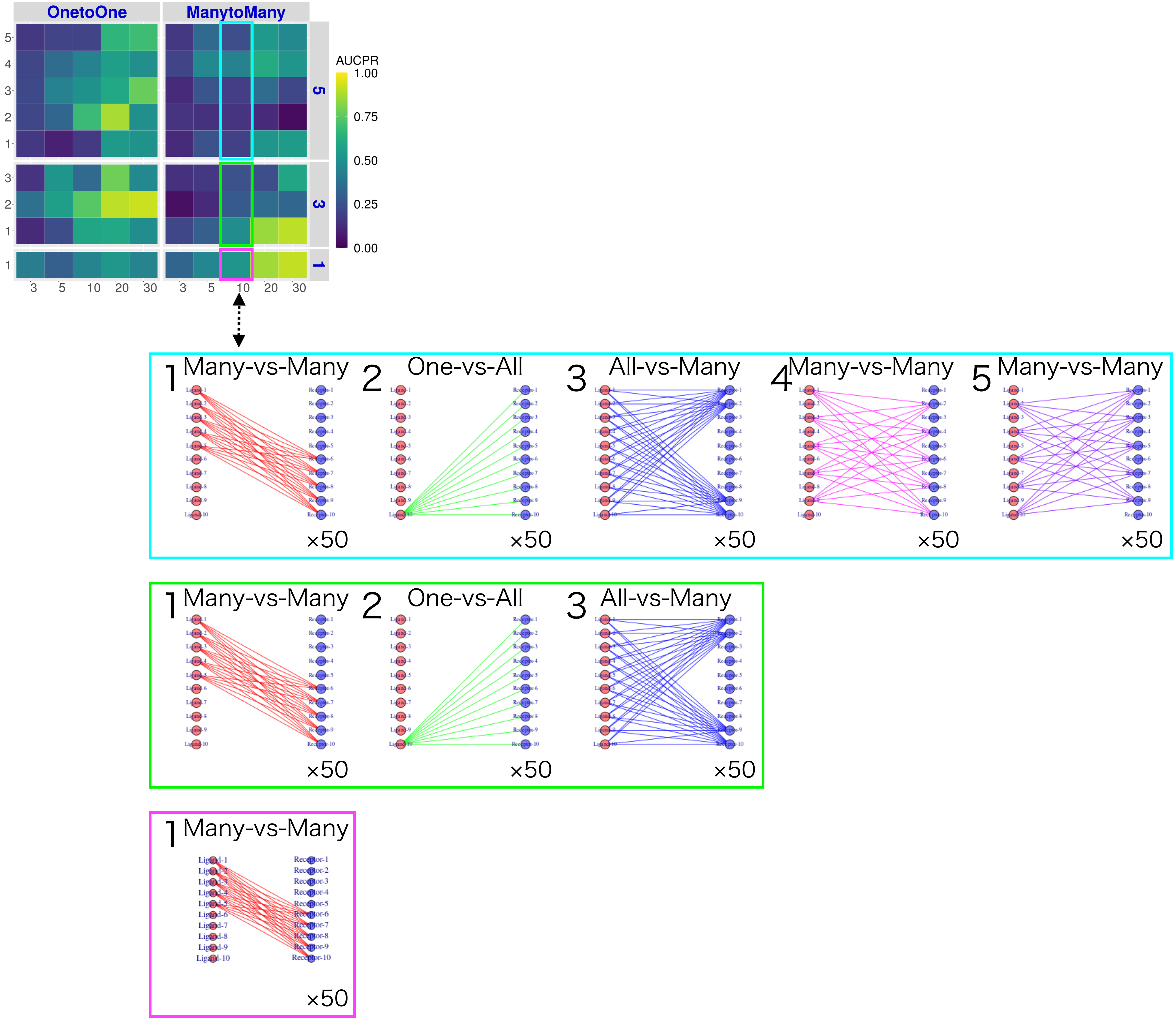

OnetoOne
ManytoMany
AUCPR
1.00
0.75
0.50
0.25
0.00
5
3
1
3
5
10
20
30
1 Many-vs-Many
2 One-vs-All
3 All-vs-Many
4 Many-vs-Many
5 Many-vs-Many
×50
1 Many-vs-Many
2 One-vs-All
3 All-vs-Many
×50
1 Many-vs-Many
×50

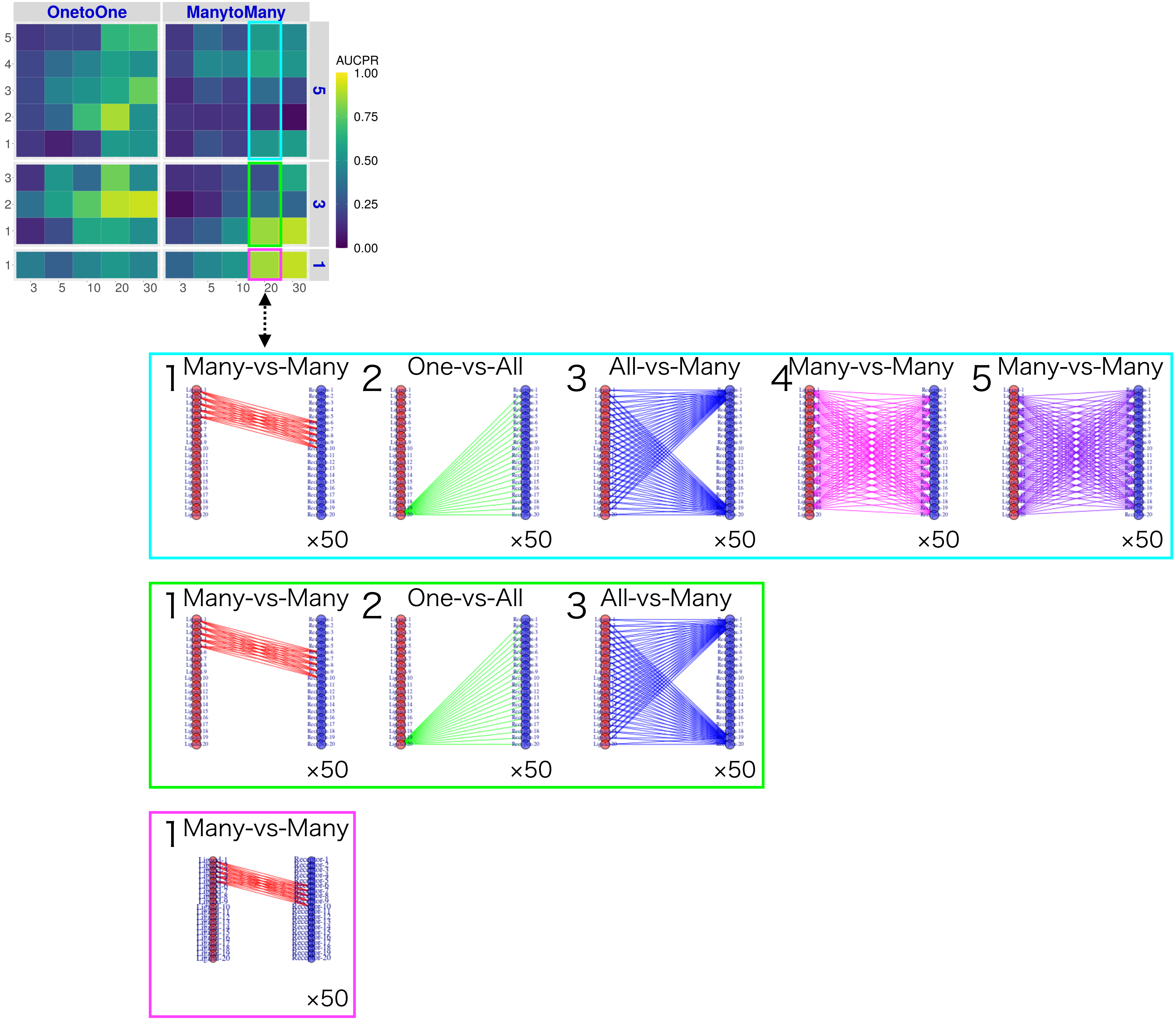

OnetoOne
ManytoMany
AUCPR
1.00
0.75
0.50
0.25
0.00
5
3
1
3
5
10
20
30
1 Many-vs-Many
2 One-vs-All
3 All-vs-Many
4 Many-vs-Many
5 Many-vs-Many
×50

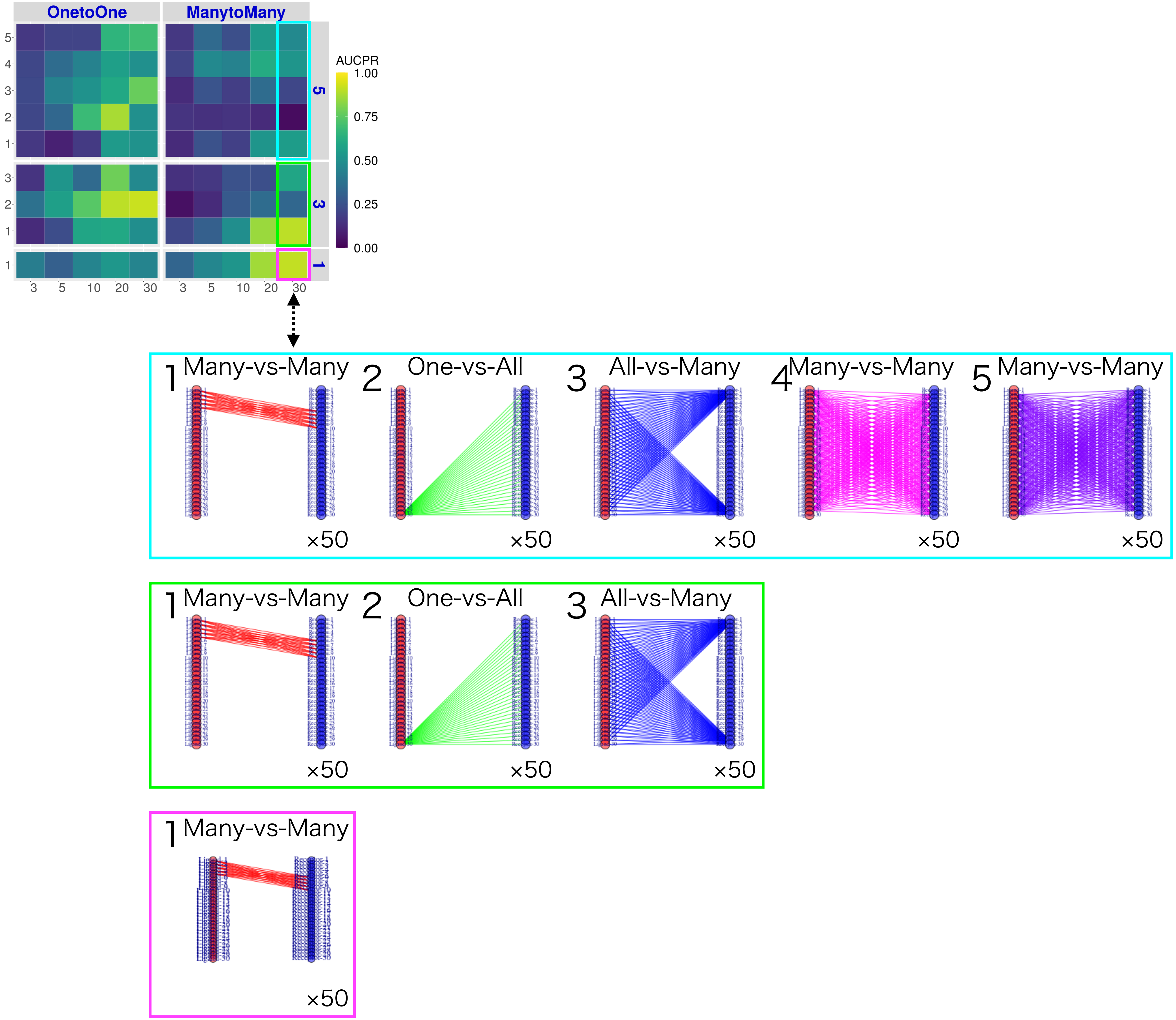

OnetoOne
ManytoMany
AUCPR
1.00
0.75
0.50
0.25
0.00
5
3
1
3
5
10
20
30
1 Many-vs-Many
2 One-vs-All
3 All-vs-Many
4 Many-vs-Many
5 Many-vs-Many
×50

### Real Datasets

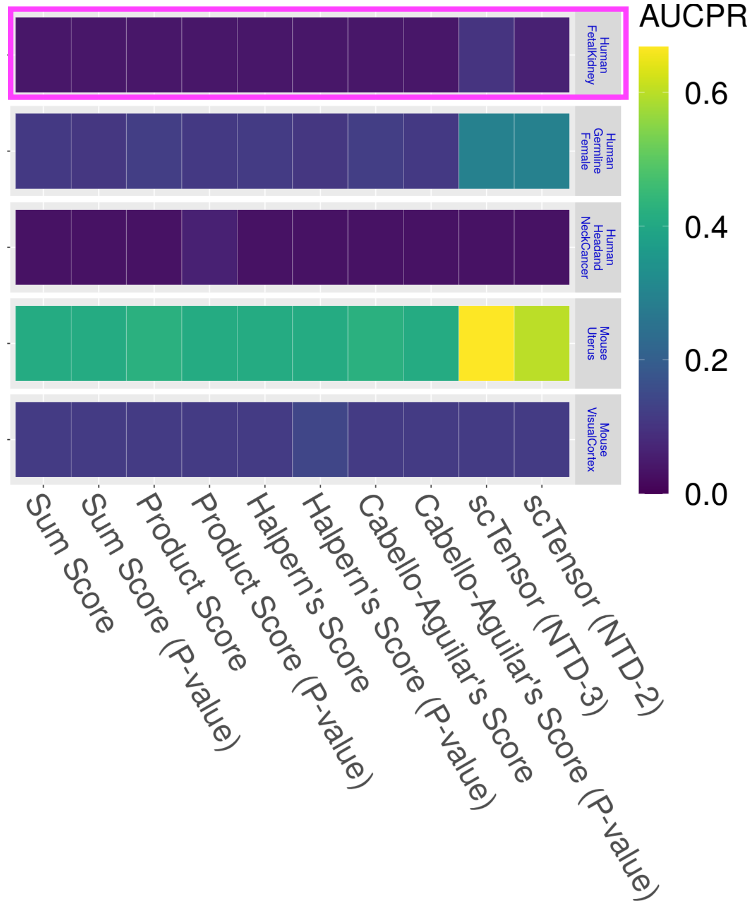
AUCPR
0.6
0.4
0.2
0.0
Human FetalKidney
Human Germline Female
Human Headand NeckCancer
Mouse Uterus
Mouse VisualCortex
Sum Score
Sum Score (P-value)
Product Score
Product Score (P-value)
Halpern's Score
Halpern's Score (P-value)
Cabello-Aguilar's Score
Cabello-Aguilar's Score (P-value)
scTensor (NTD-3)
scTensor (NTD-2)

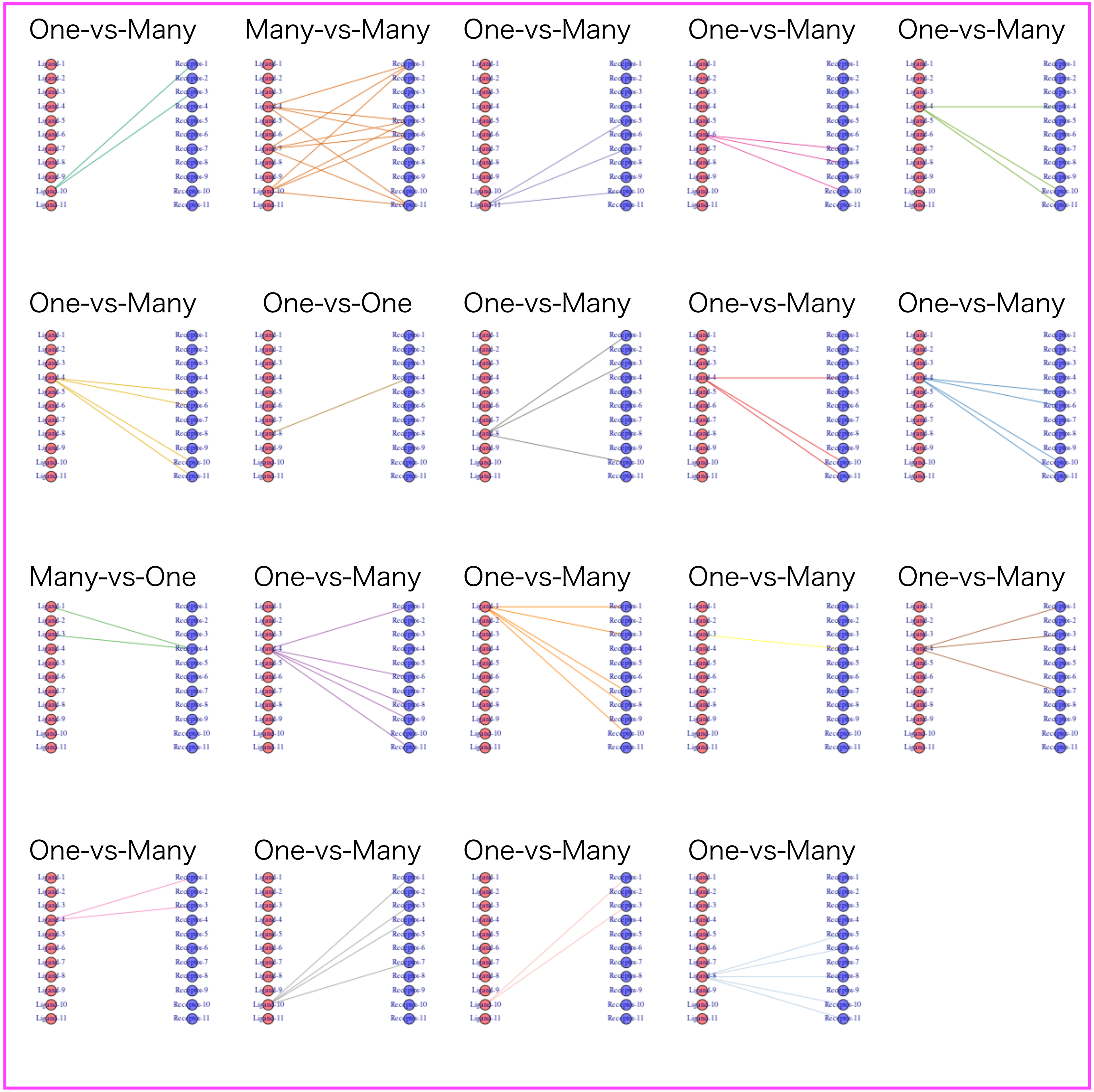
One-vs-Many
Many-vs-Many
One-vs-Many
One-vs-Many
One-vs-Many
One-vs-Many
One-vs-One
One-vs-Many
One-vs-Many
One-vs-Many
Many-vs-One
One-vs-Many
One-vs-Many
One-vs-Many
One-vs-Many
One-vs-Many
One-vs-Many
One-vs-Many
One-vs-Many

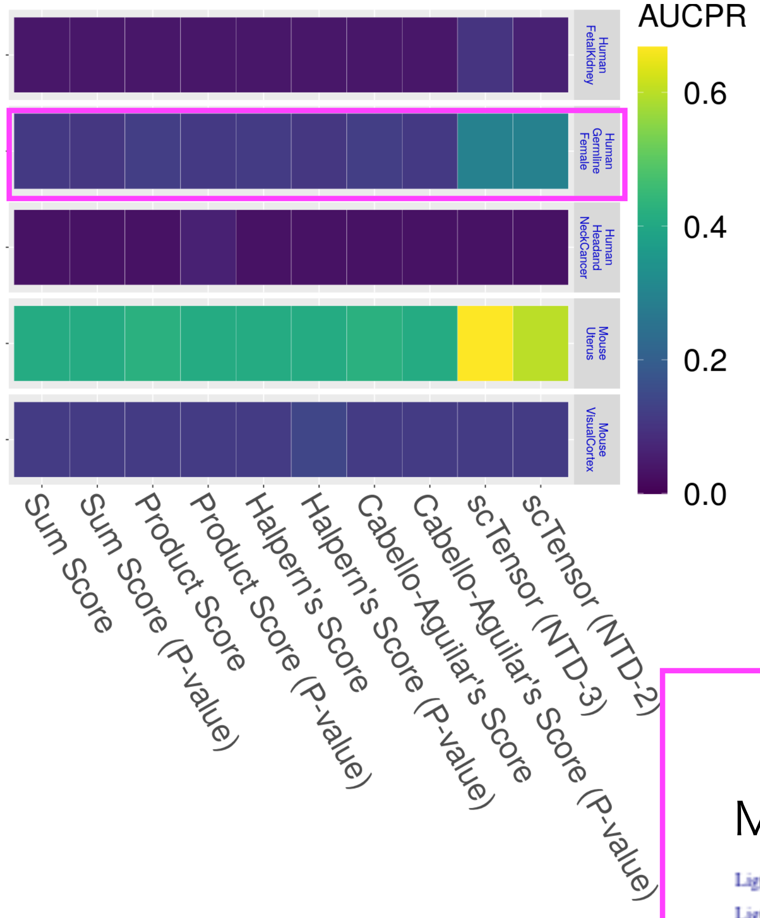
AUCPR
0.6
0.4
0.2
0.0
Human FetalKidney
Human Germline Female
Human Headand NeckCancer
Mouse Uterus
Mouse VisualCortex
Sum Score
Sum Score (P-value)
Product Score
Product Score (P-value)
Halpern's Score
Halpern's Score (P-value)
Cabello-Aguilar's Score
Cabello-Aguilar's Score (P-value)
scTensor (NTD-3)
scTensor (NTD-2)

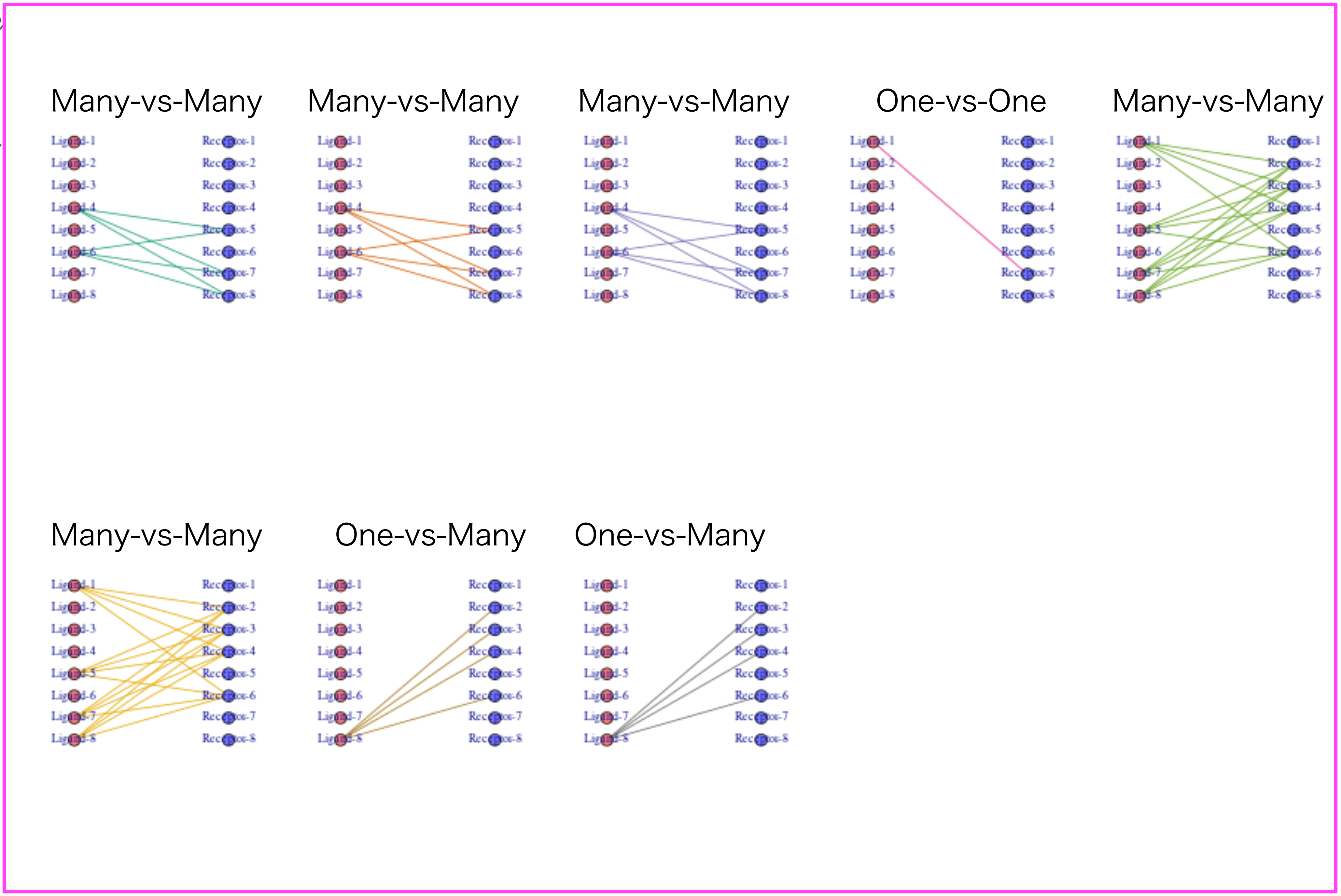
Many-vs-Many
Many-vs-Many
Many-vs-Many
One-vs-One
Many-vs-Many
Many-vs-Many
One-vs-Many
One-vs-Many

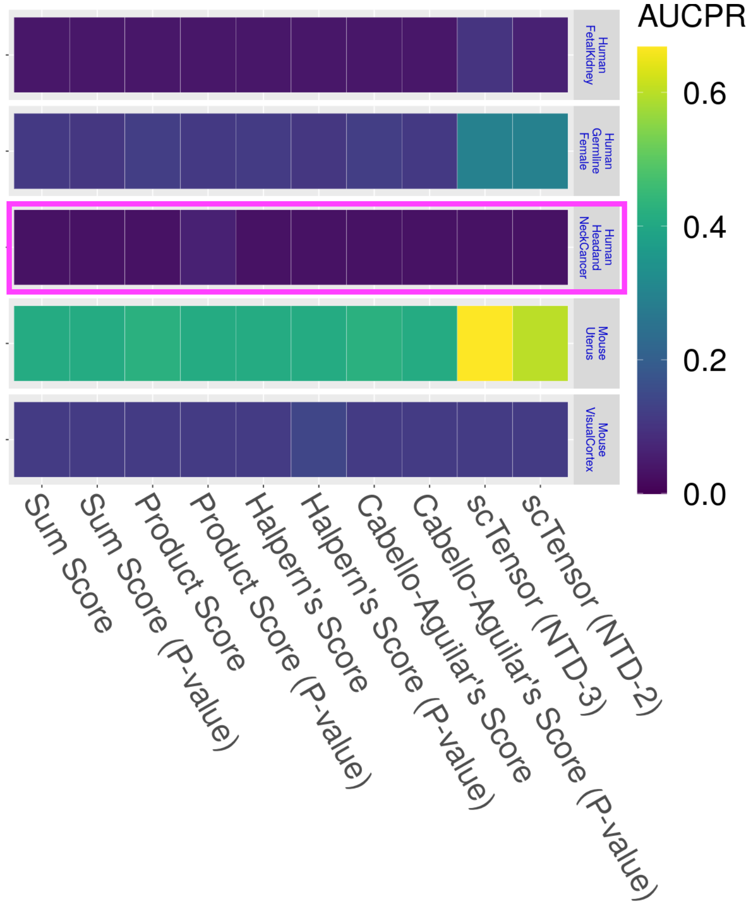
AUCPR
0.6
0.4
0.2
0.0
Human FetalKidney
Human Germline Female
Human Headand NeckCancer
Mouse Uterus
Mouse VisualCortex
Sum Score
Sum Score (P-value)
Product Score
Product Score (P-value)
Halpern's Score
Halpern's Score (P-value)
Cabello-Aguilar's Score
Cabello-Aguilar's Score (P-value)
scTensor (NTD-3)
scTensor (NTD-2)

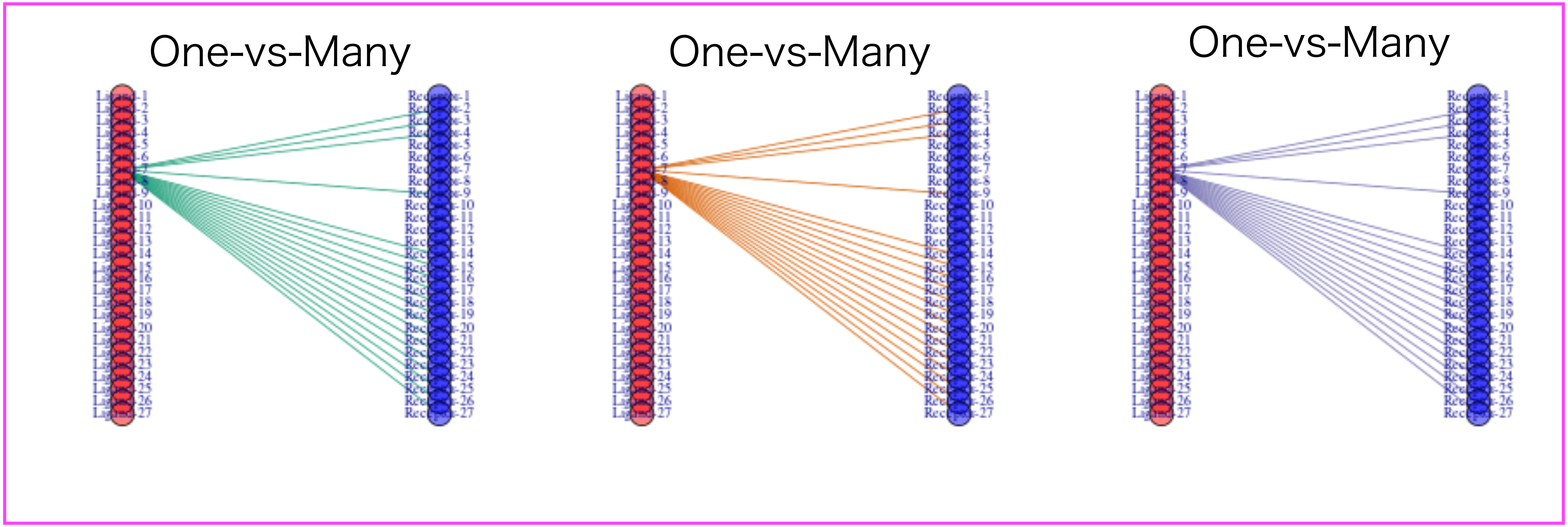
One-vs-Many
One-vs-Many
One-vs-Many

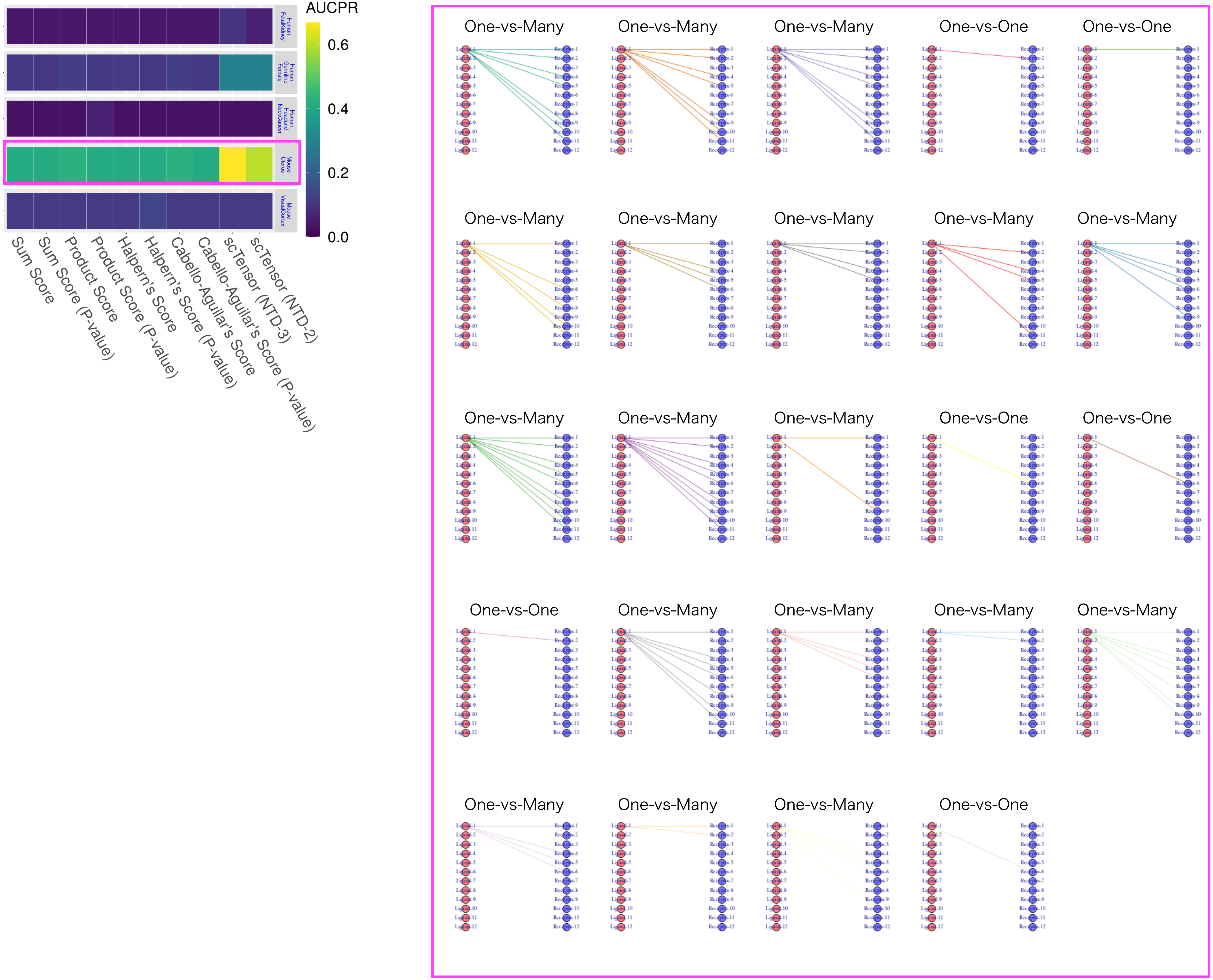

AUCPR
0.6
0.4
0.2
0.0
Human FetalKidney
Human Germline Female
Human HeadandNeckCancer
Mouse Uterus
Mouse VisualCortex
Sum Score
Sum Score (P-value)
Product Score
Product Score (P-value)
Halpern's Score
Halpern's Score (P-value)
Cabello-Aguilar's Score
Cabello-Aguilar's Score (P-value)
scTensor (NTD-3)
scTensor (NTD-2)
One-vs-Many
One-vs-Many
One-vs-Many
One-vs-One
One-vs-One
One-vs-Many
One-vs-Many
One-vs-Many
One-vs-Many
One-vs-Many
One-vs-Many
One-vs-Many
One-vs-Many
One-vs-One
One-vs-One
One-vs-One
One-vs-Many
One-vs-Many
One-vs-Many
One-vs-Many
One-vs-Many
One-vs-Many
One-vs-Many
One-vs-One

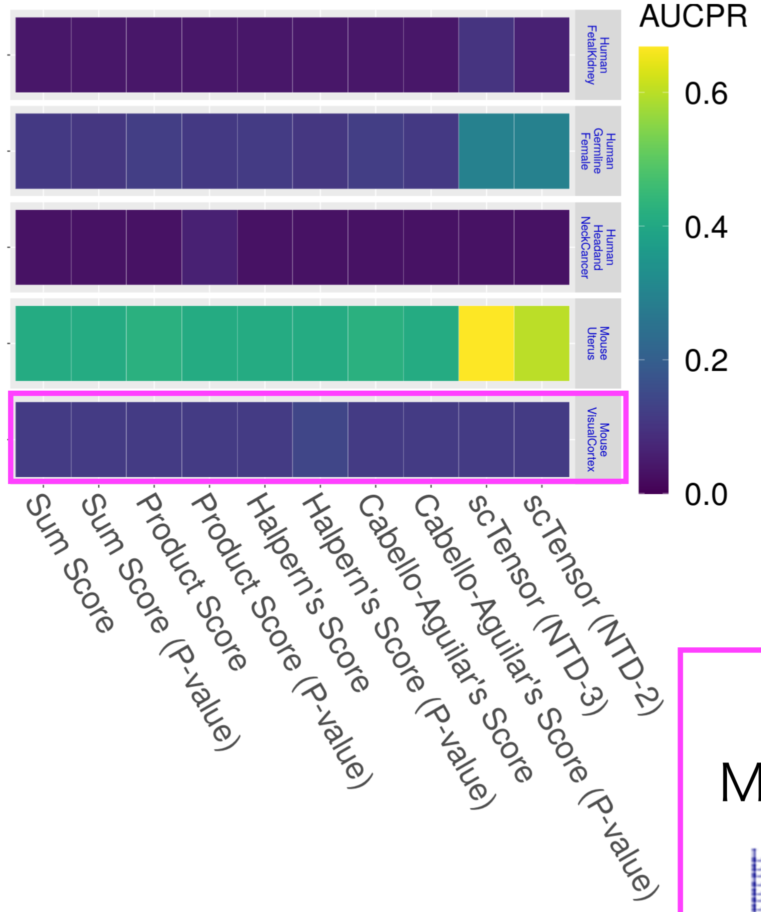
AUCPR
0.6
0.4
0.2
0.0
Human FetalKidney
Human Germline Female
Human Headand NeckCancer
Mouse Uterus
Mouse VisualCortex
Sum Score
Sum Score (P-value)
Product Score
Product Score (P-value)
Halpern's Score
Halpern's Score (P-value)
Cabello-Aguilar's Score
Cabello-Aguilar's Score (P-value)
scTensor (NTD-3)
scTensor (NTD-2)

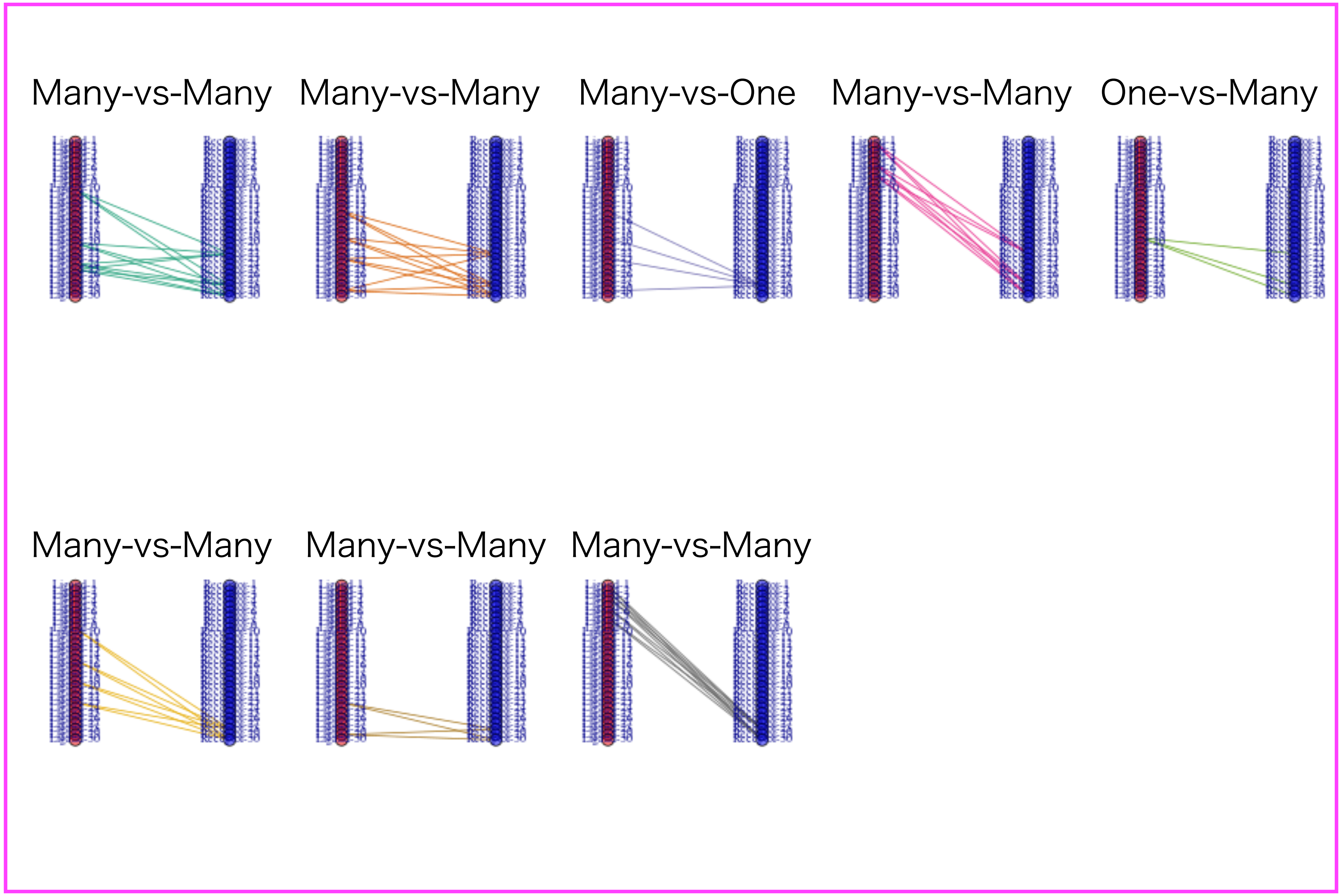
Many-vs-Many
Many-vs-Many
Many-vs-One
Many-vs-Many
One-vs-Many
Many-vs-Many
Many-vs-Many
Many-vs-Many
