## Supplementary material for "scTensor detects many-to-many cell–cell interactions from single cell RNA-sequencing data": https://zenodo.org/record/7412280/files/03_AUCROC.pdf?download=1

### Simulated Datasets

**E2 (Summary)** The value ranges 0 to 1 (the closer to 1, the better)

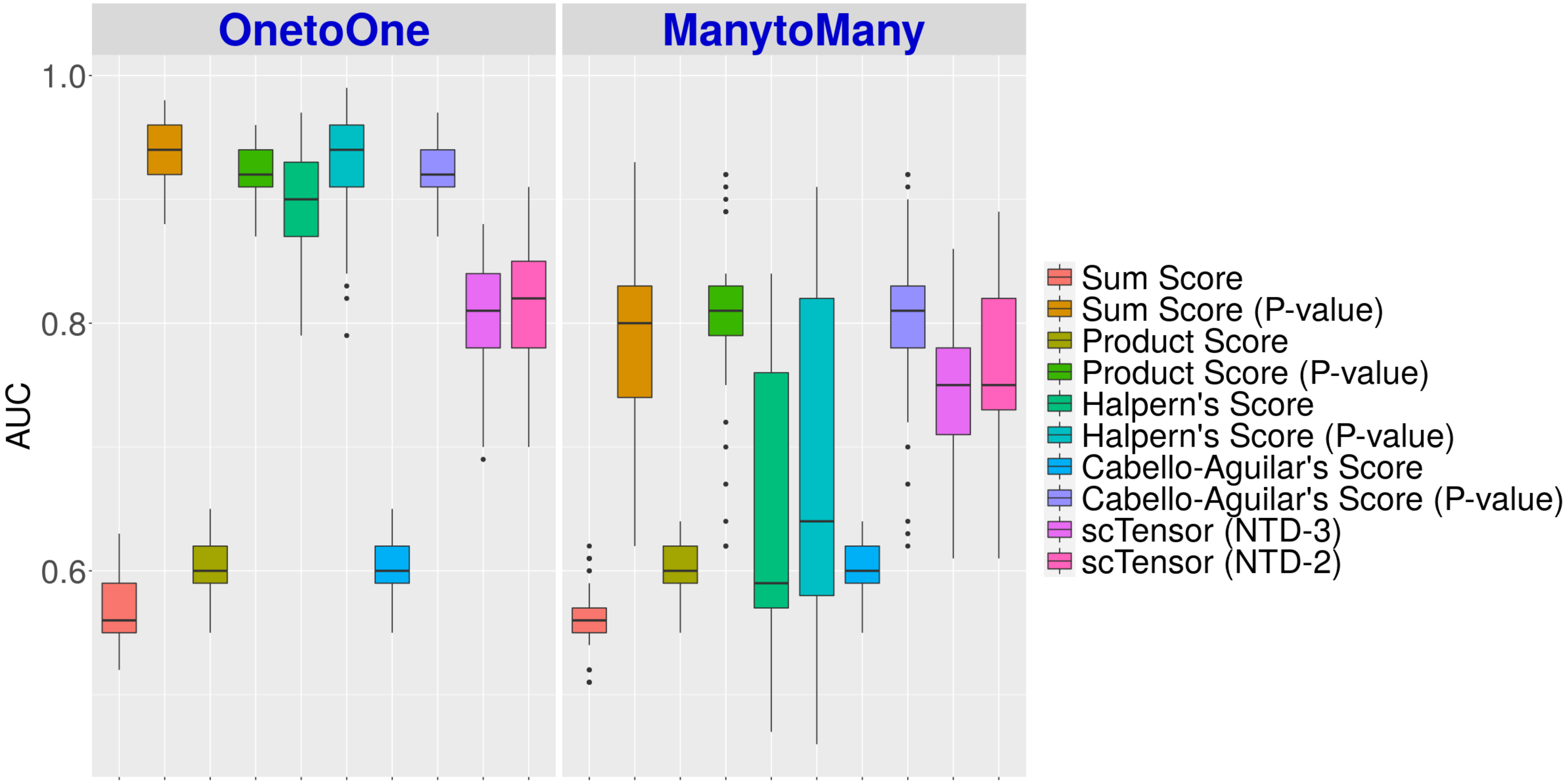

E5 (Summary) The value ranges 0 to 1 (the closer to 1, the better)

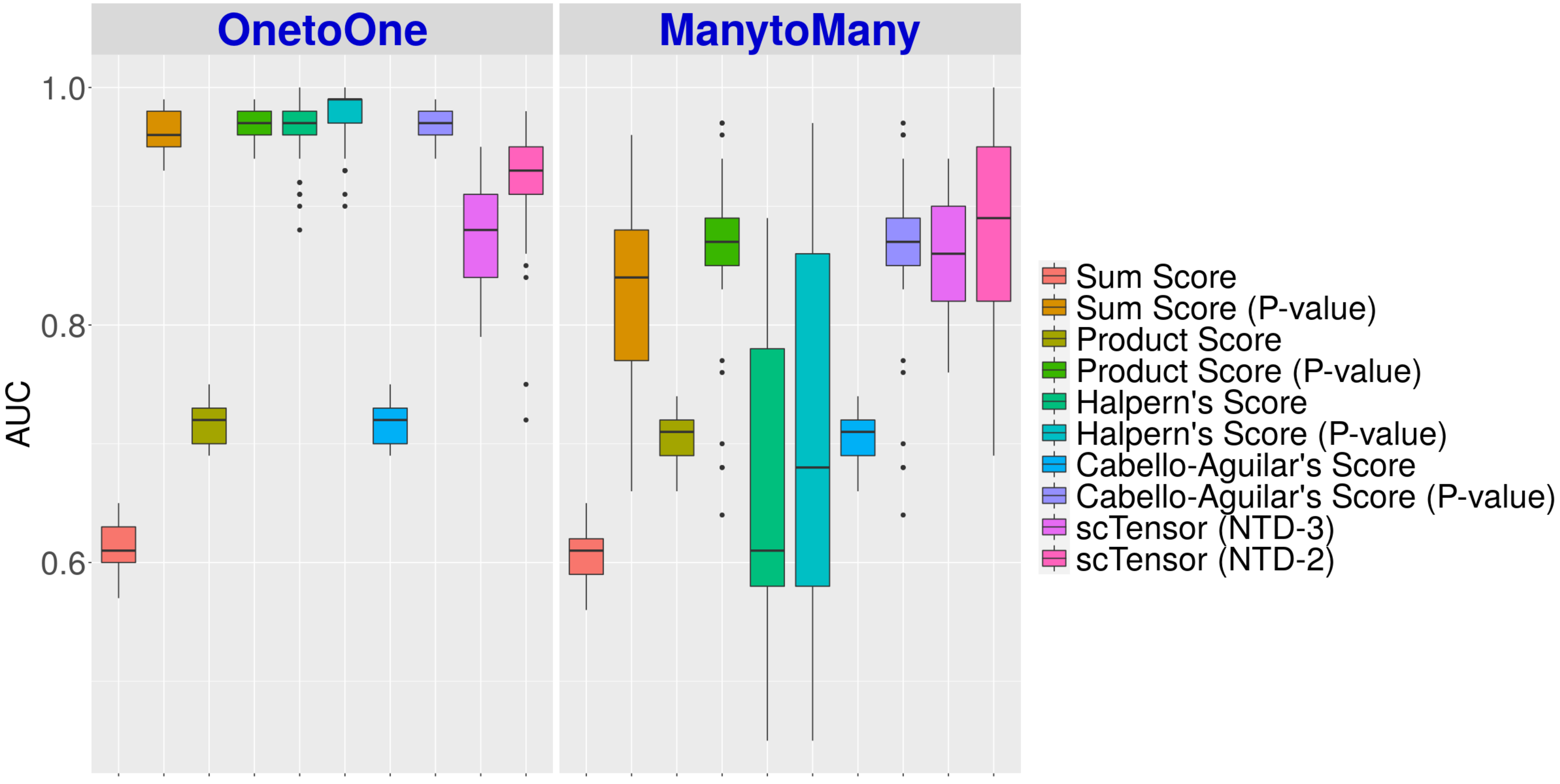

E10 (Summary) The value ranges 0 to 1 (the closer to 1, the better)

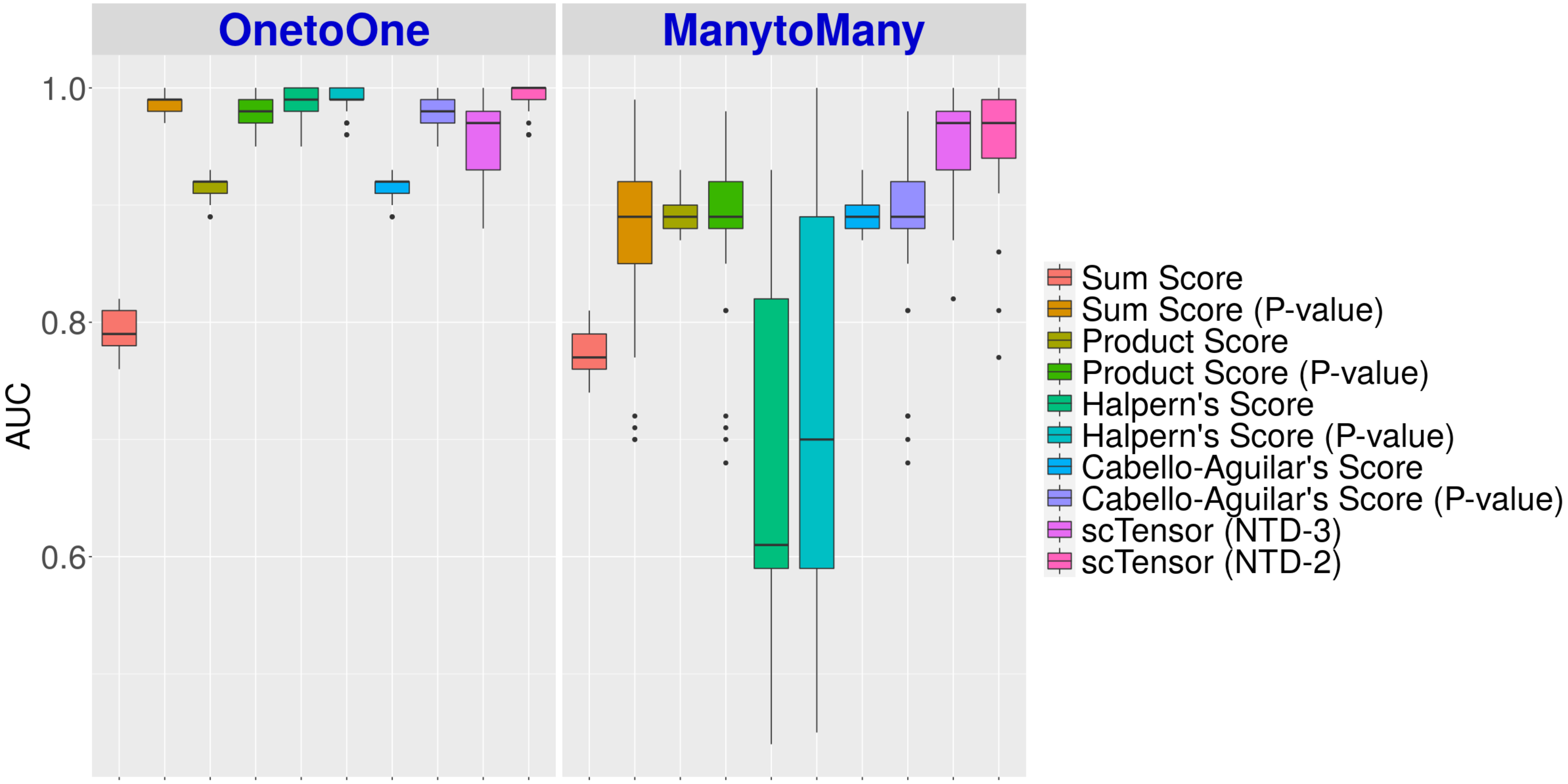

### E2 (Details)

The value ranges 0 to 1 (the closer to 1, the better)

#### Sum Score

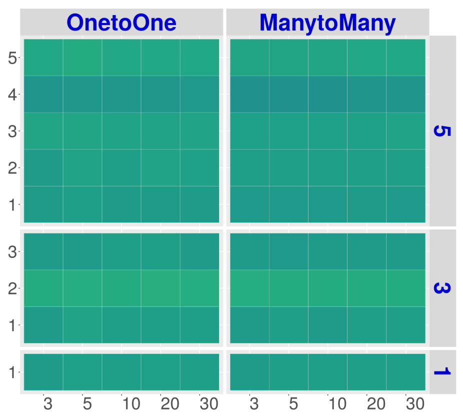


#### Product Score

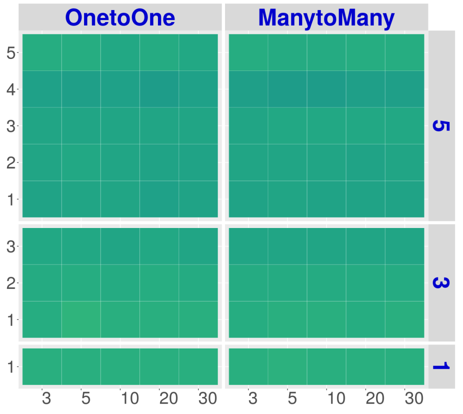


#### Halpern's Score

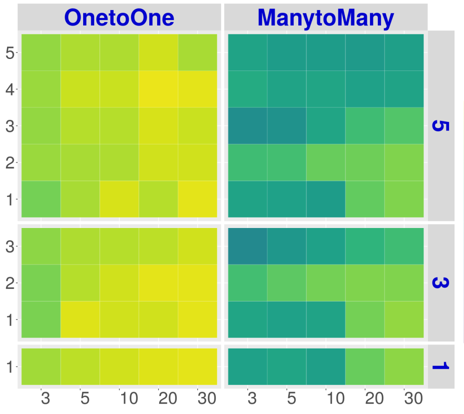


#### Cabello-Aguilar's Score

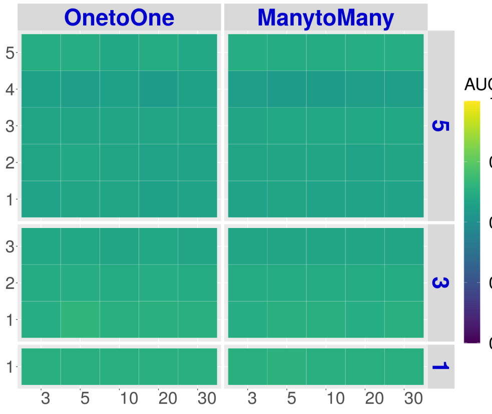


#### scTensor (NTD-3)

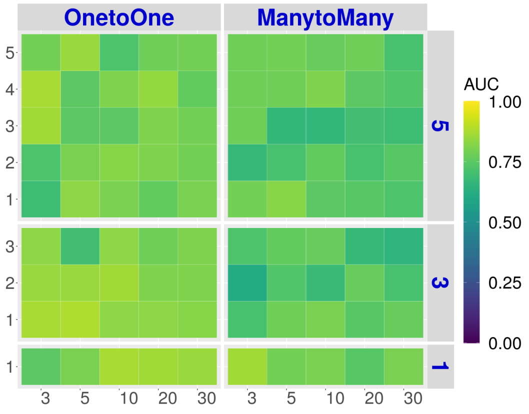


#### Sum Score (P-value)

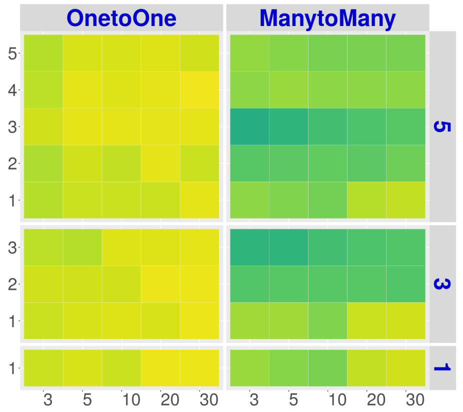


#### Product Score (P-value)

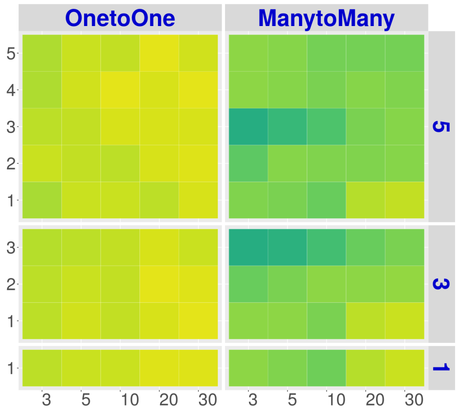


#### Halpern's Score (P-value)

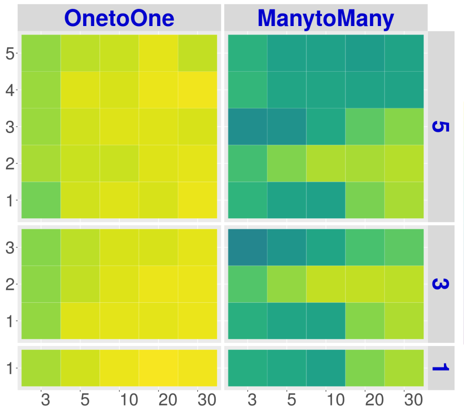


#### Cabello-Aguilar's Score (P-value)

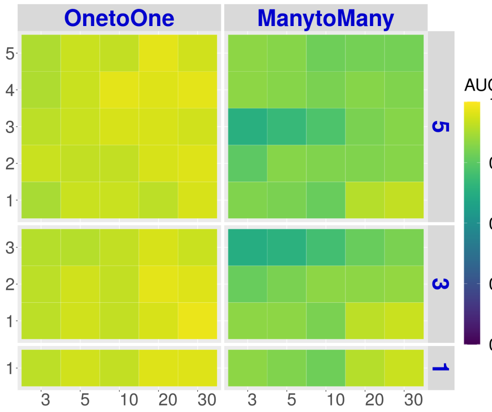


#### scTensor (NTD-2)

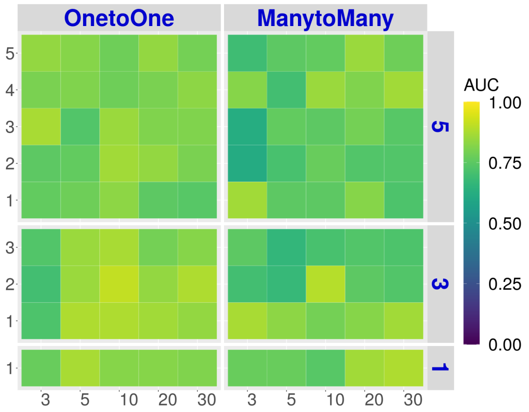

#### E5 (Details)

The value ranges 0 to 1 (the closer to 1, the better)

##### Sum Score

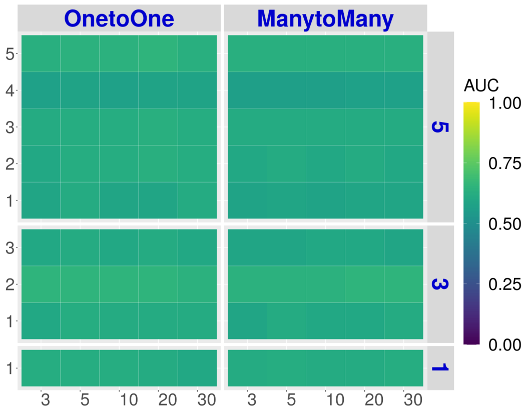


##### Product Score

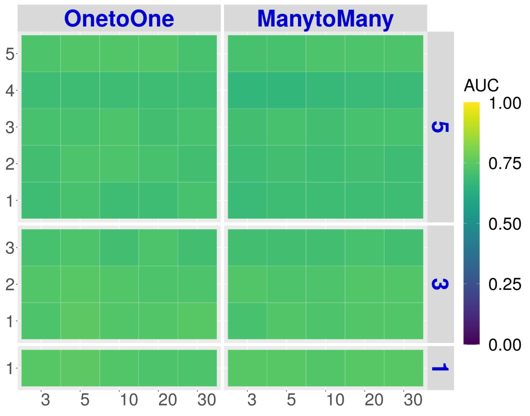


##### Halpern's Score

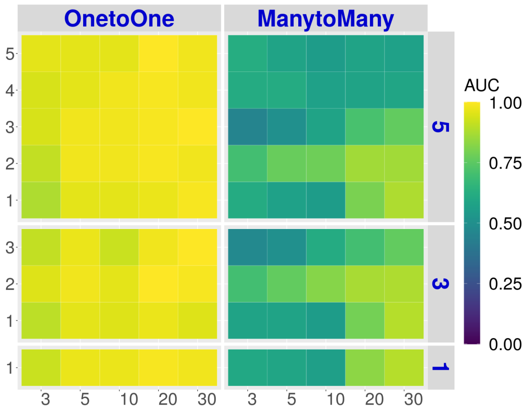


##### Cabello-Aguilar's Score

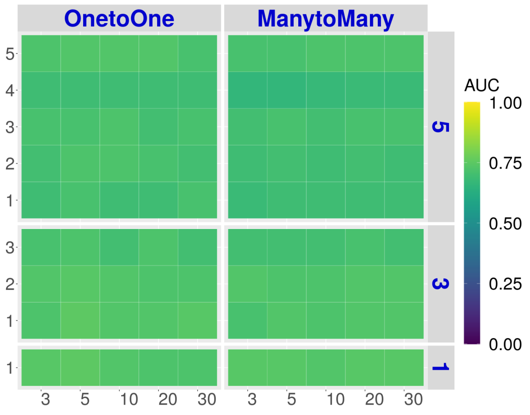


##### scTensor (NTD-3)

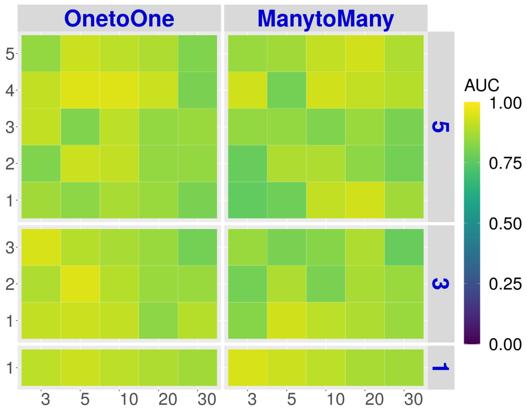


##### Sum Score (P-value)

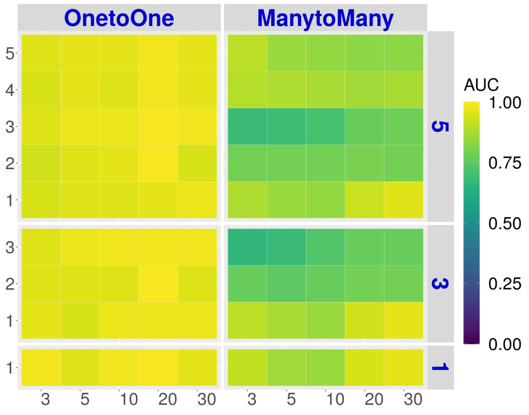


##### Product Score (P-value)

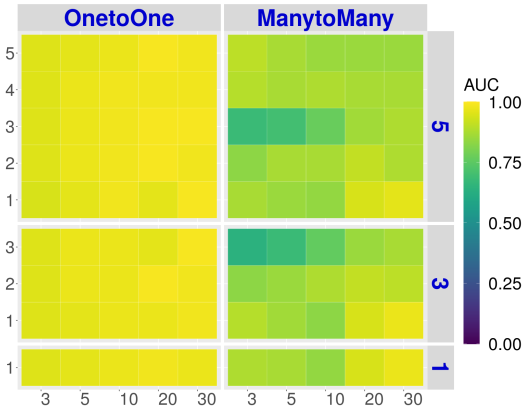


##### Halpern's Score (P-value)

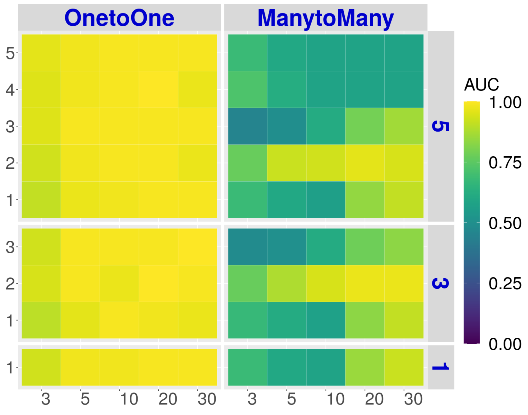


##### Cabello-Aguilar's Score (P-value)

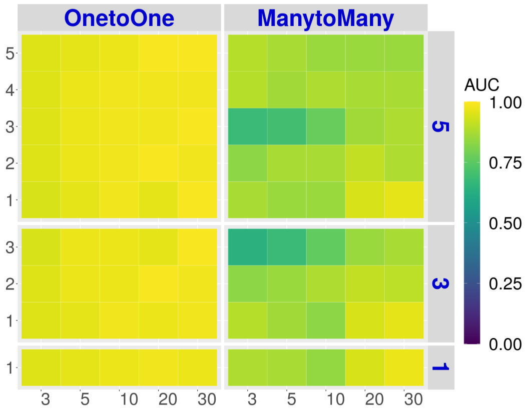


##### scTensor (NTD-2)

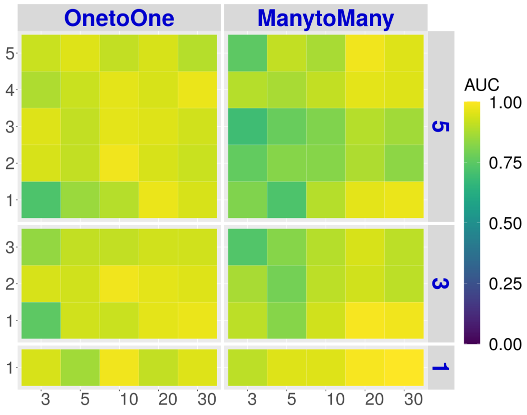

**E10 (Details)** The value ranges 0 to 1 (the closer to 1, the better)

#### Sum Score

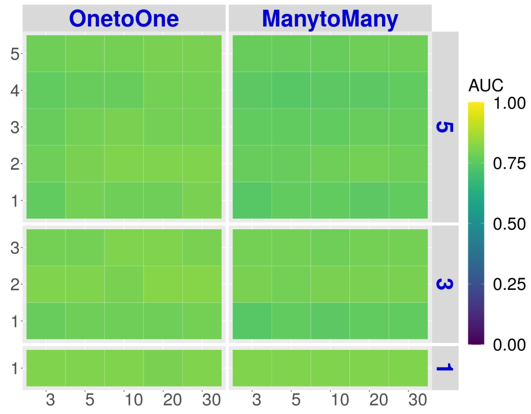


#### Product Score

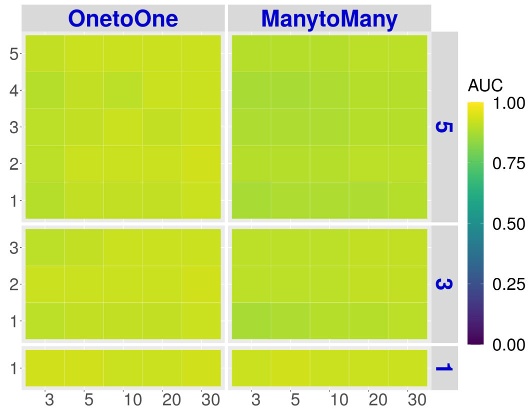


#### Halpern's Score

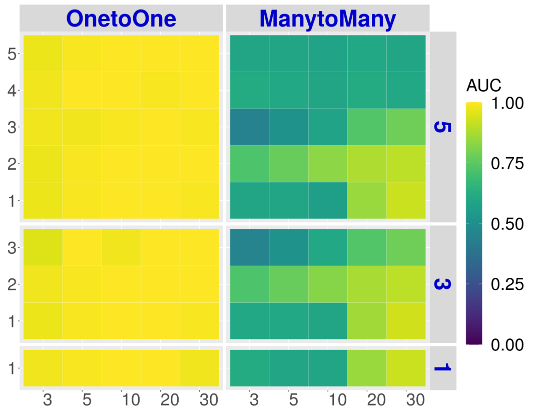


#### Cabello-Aguilar's Score

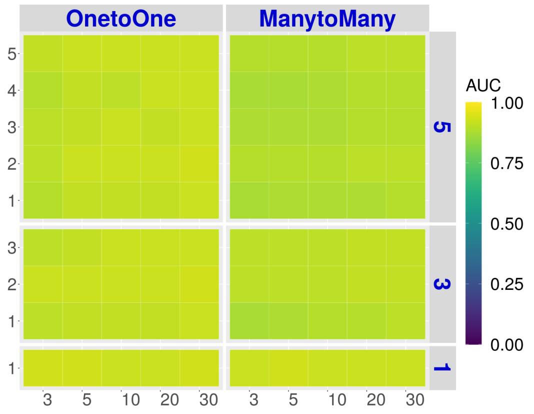


#### scTensor (NTD-3)

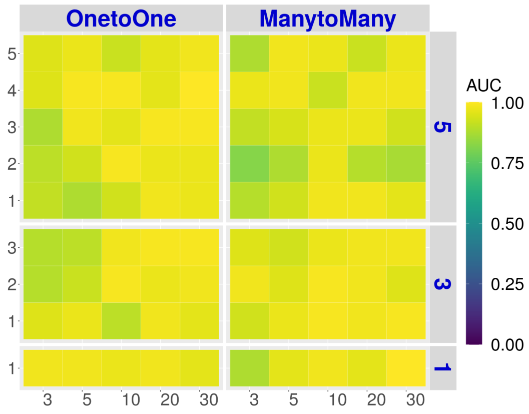


#### Sum Score (P-value)

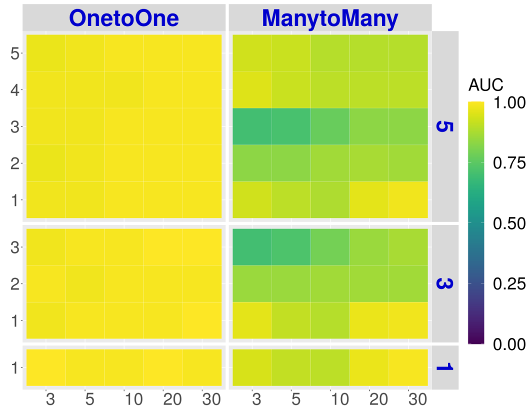


#### Product Score (P-value)

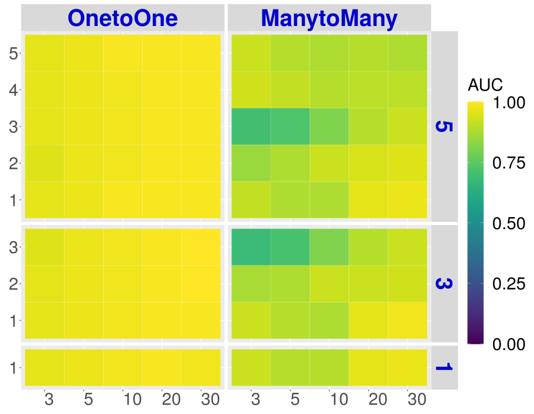


#### Halpern's Score (P-value)

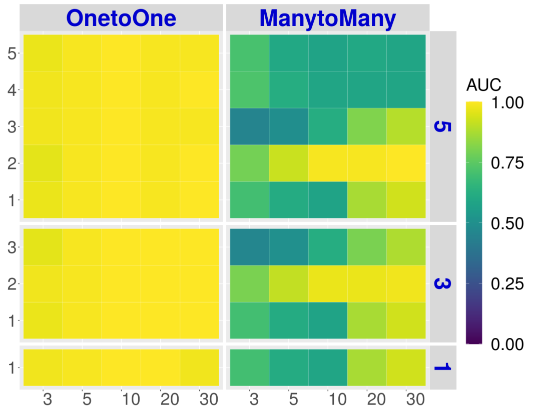


#### Cabello-Aguilar's Score (P-value)

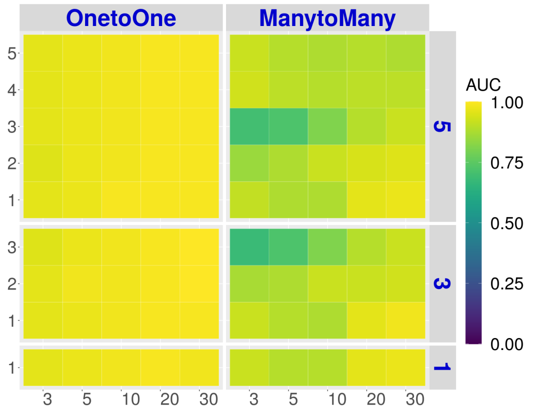


#### scTensor (NTD-2)

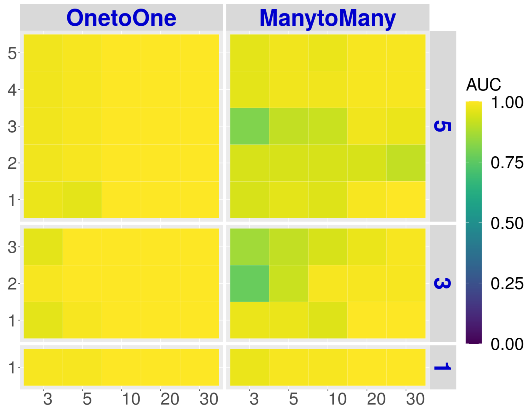

### Real Datasets

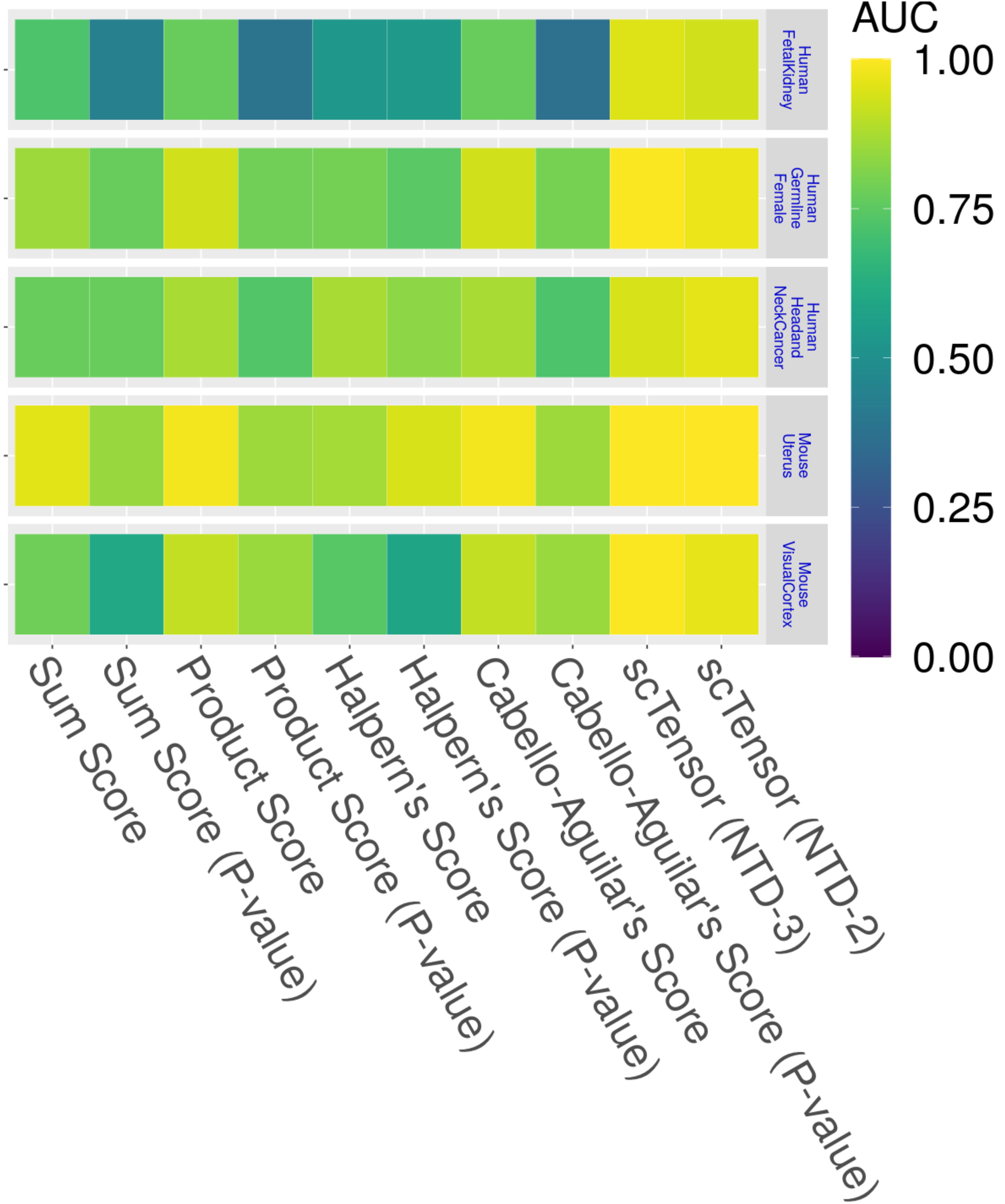
AUC
1.00
0.75
0.50
0.25
0.00
Human FetalKidney
Human Germline Female
Human Headand NeckCancer
Mouse Uterus
Mouse VisualCortex
Sum Score
Sum Score (P-value)
Product Score
Product Score (P-value)
Halpern's Score
Halpern's Score (P-value)
Cabello-Aguilar's Score
Cabello-Aguilar's Score (P-value)
scTensor (NTD-3)
scTensor (NTD-2)
