## Supplementary material for "scTensor detects many-to-many cell–cell interactions from single cell RNA-sequencing data": https://zenodo.org/record/7412280/files/04_AUCPR.pdf?download=1

### Simulated Datasets

**E2 (Summary)** The value ranges 0 to 1 (the closer to 1, the better)

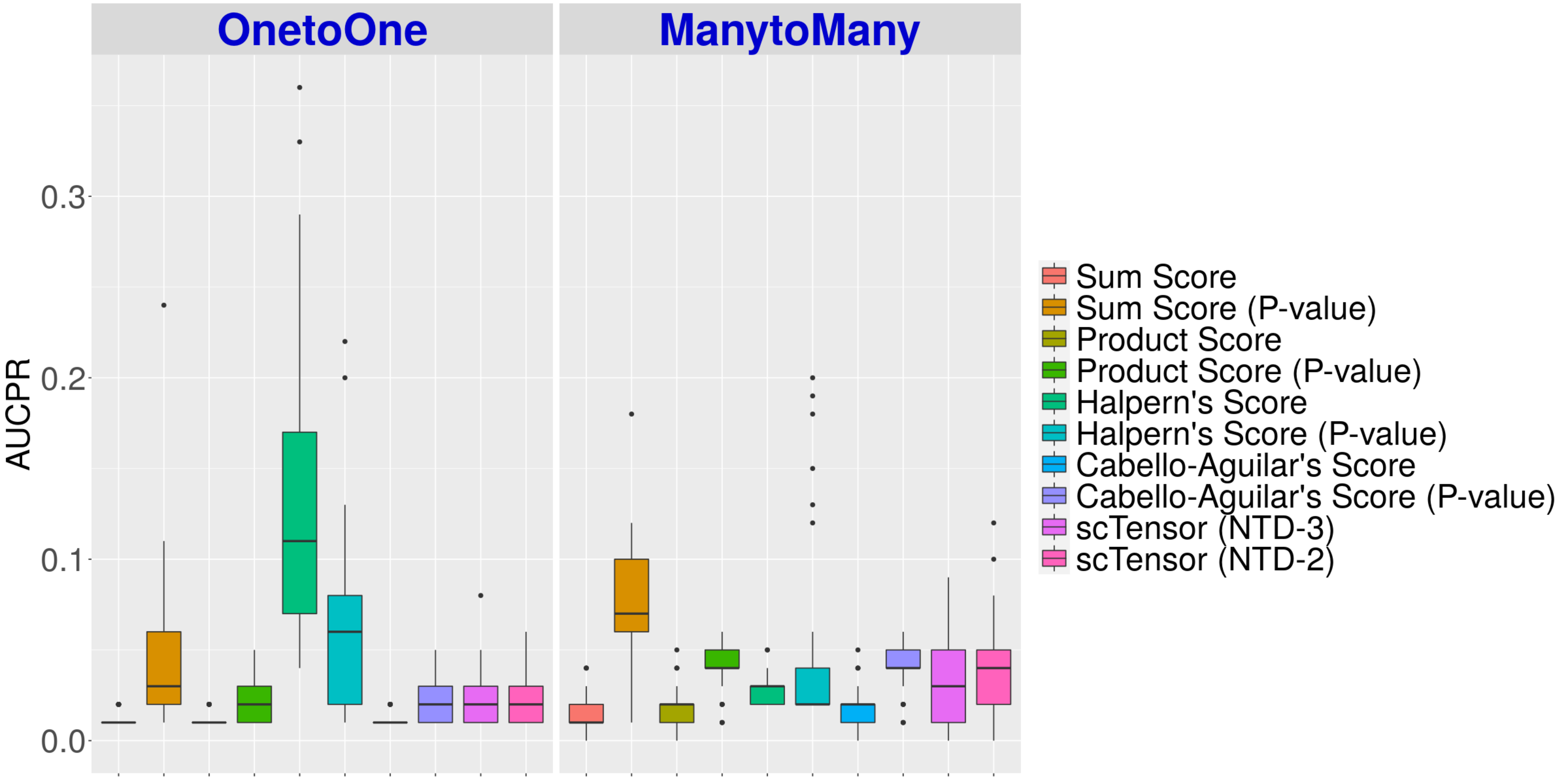

**E5 (Summary)** The value ranges 0 to 1 (the closer to 1, the better)

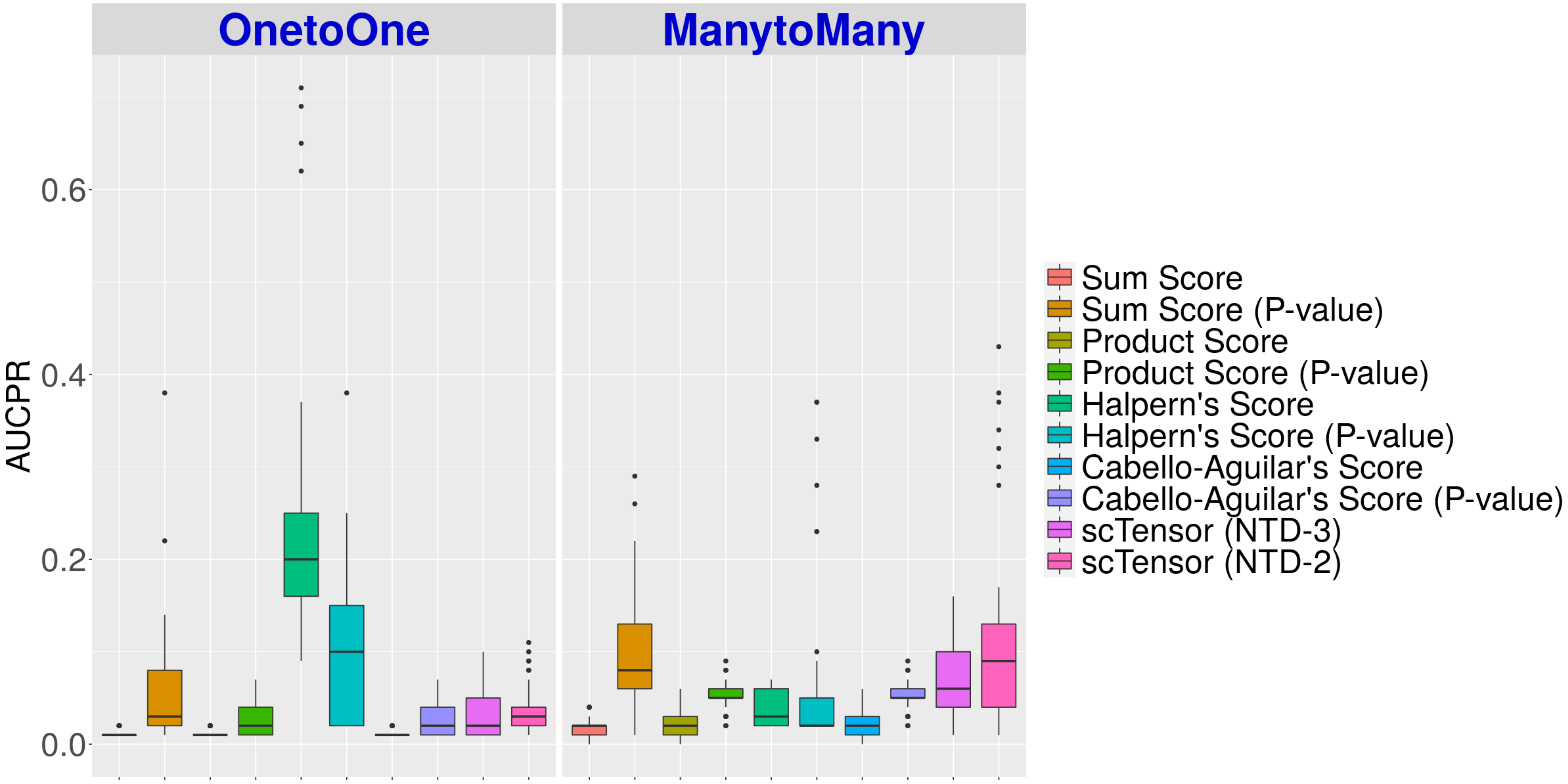

**E10 (Summary)** The value ranges 0 to 1 (the closer to 1, the better)

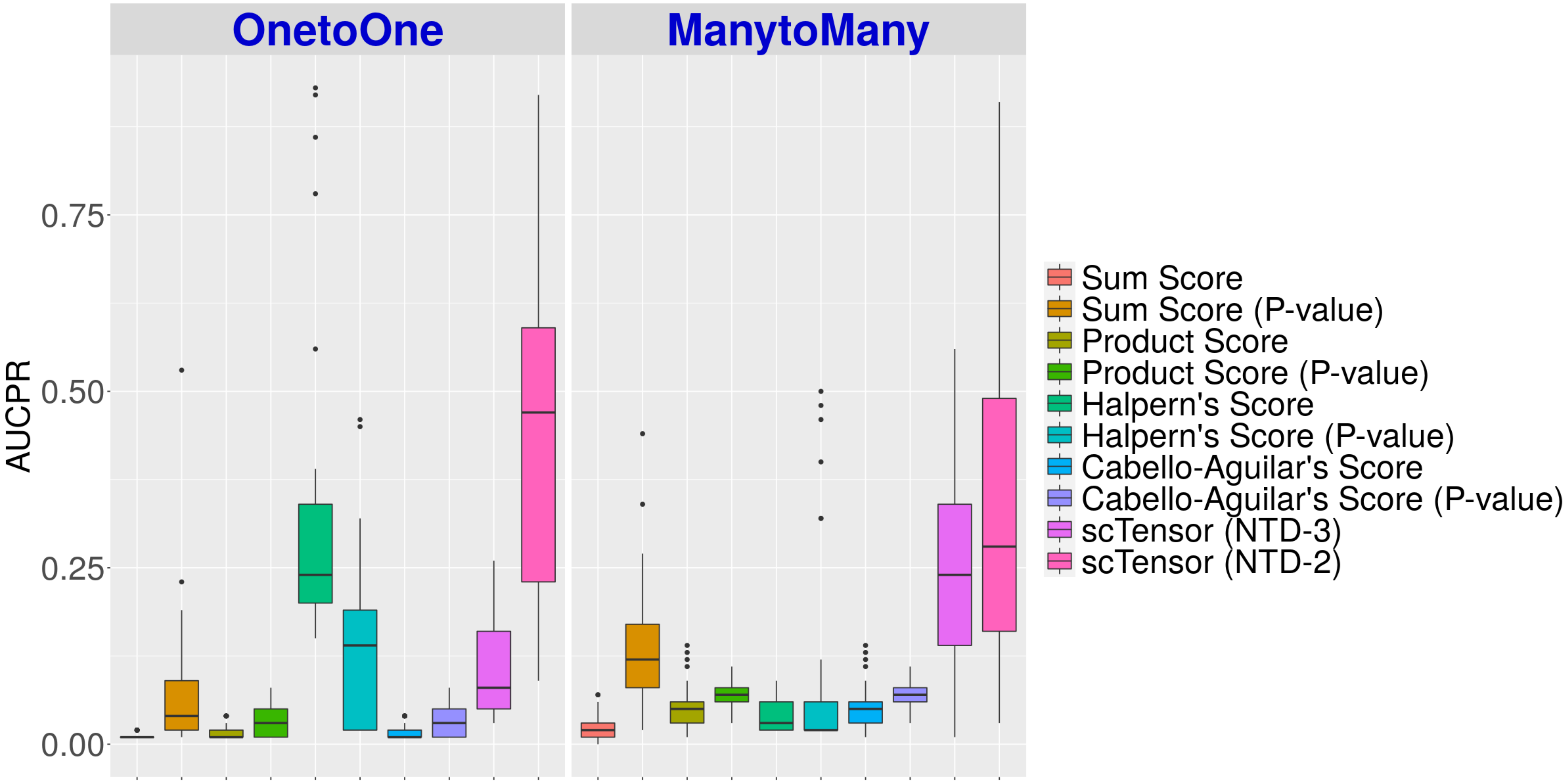

**E2 (Details)** The value ranges 0 to 1 (the closer to 1, the better)

#### Sum Score

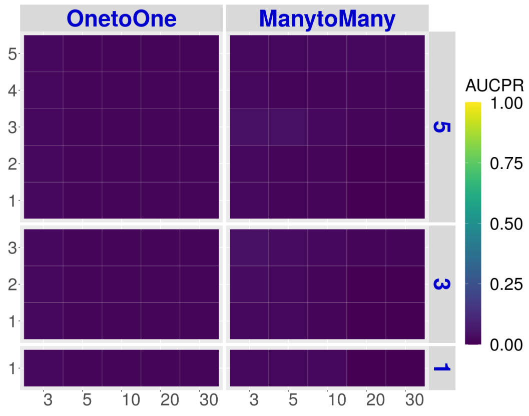


#### Product Score

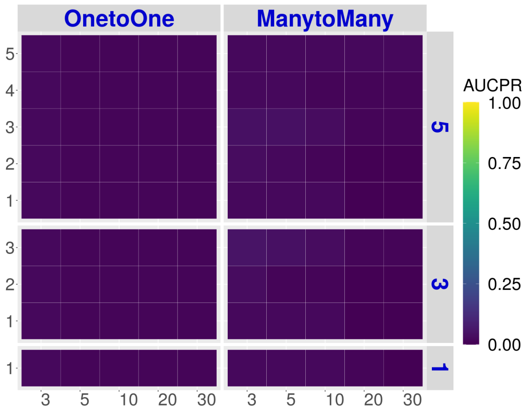


#### Halpern's Score

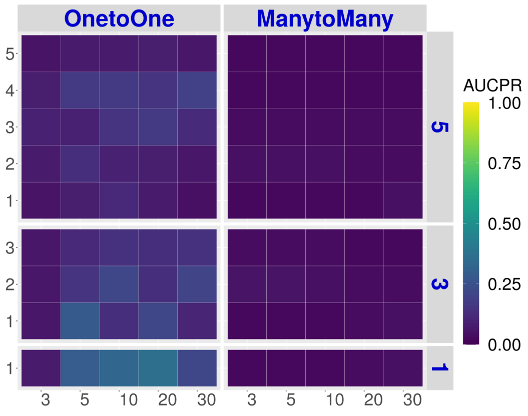


#### Cabello-Aguilar's Score

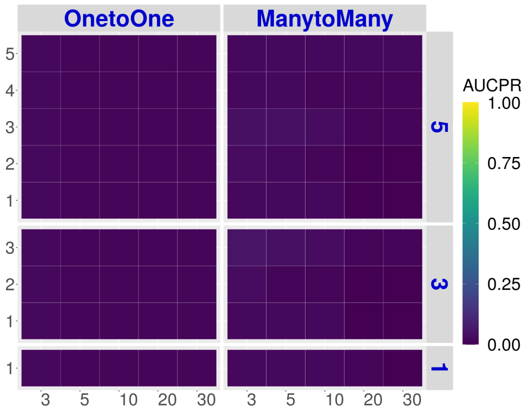


#### scTensor (NTD-3)

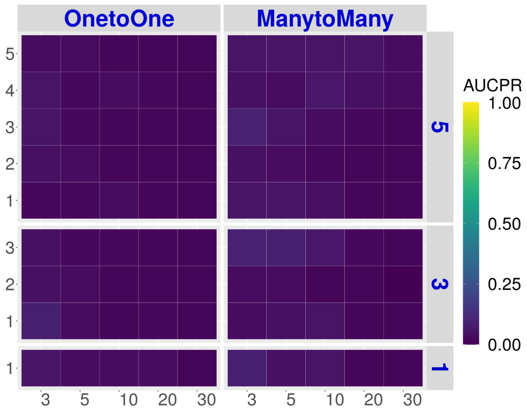


#### Sum Score (P-value)

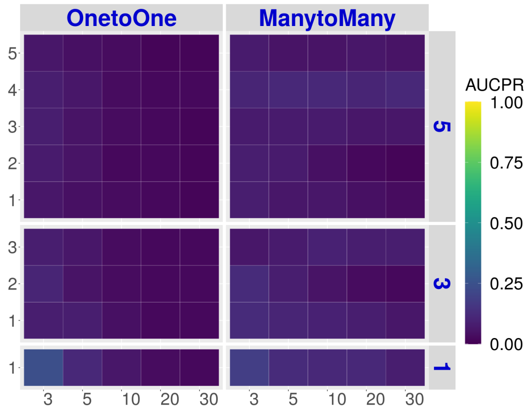


#### Product Score (P-value)

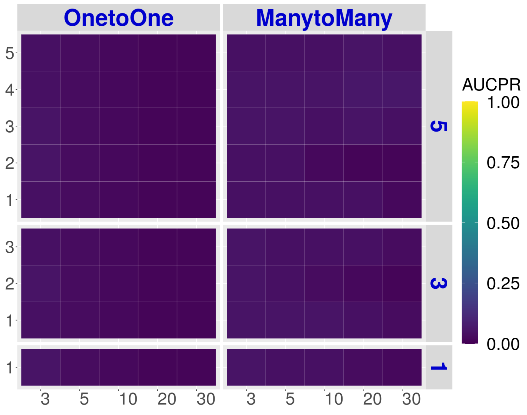


#### Halpern's Score (P-value)

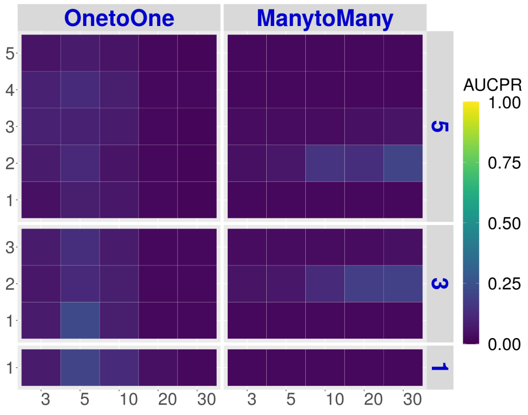


#### Cabello-Aguilar's Score (P-value)

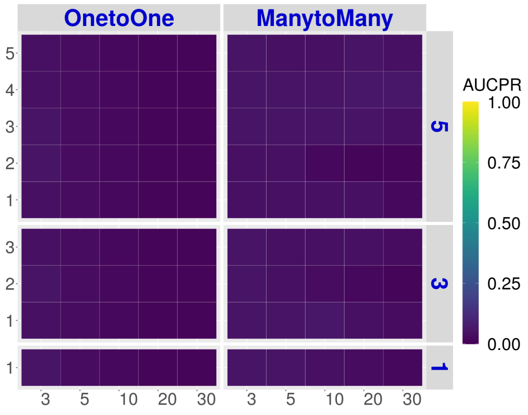


#### scTensor (NTD-2)

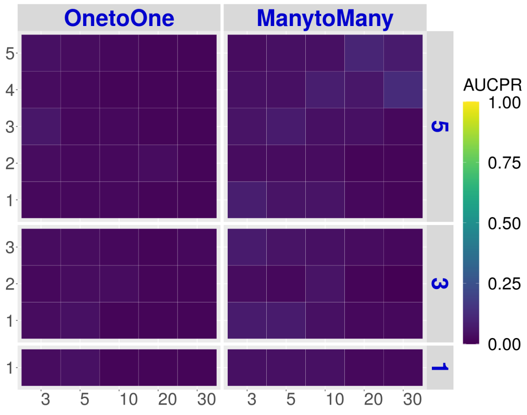

### E5 (Details)

The value ranges 0 to 1 (the closer to 1, the better)

#### Sum Score

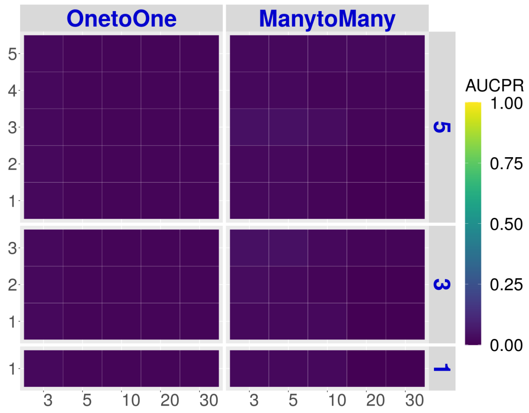


#### Product Score

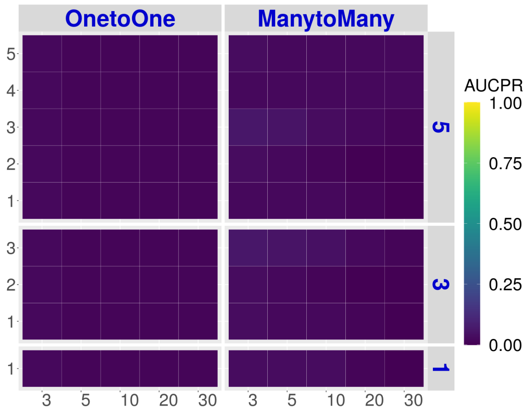


#### Halpern's Score

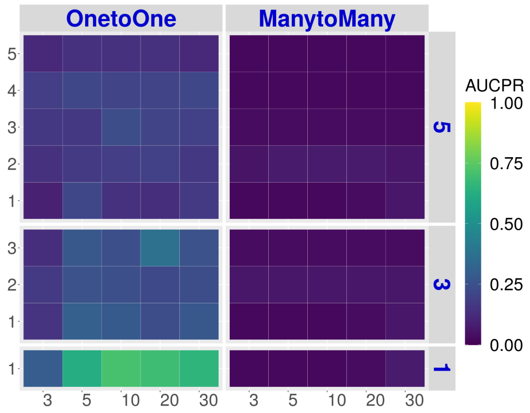


#### Cabello-Aguilar's Score

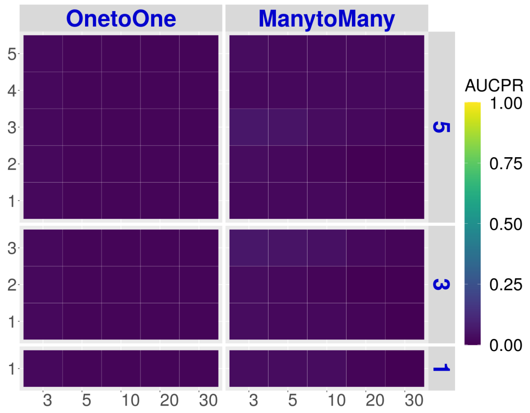


#### scTensor (NTD-3)

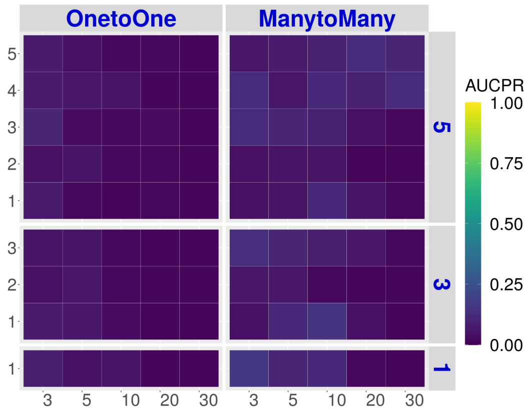


#### Sum Score (P-value)

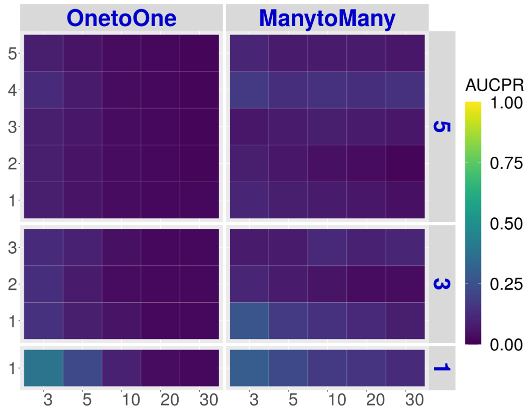


#### Product Score (P-value)

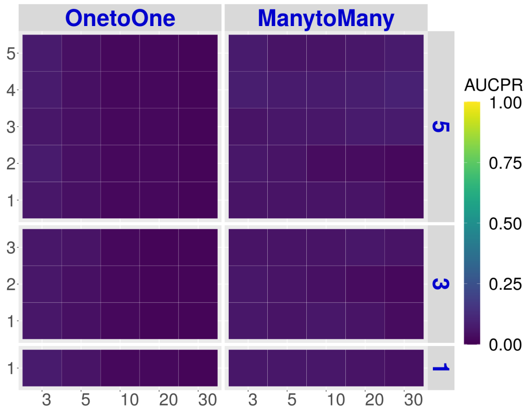


#### Halpern's Score (P-value)

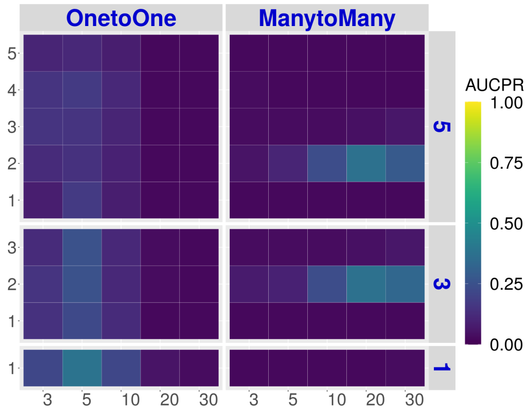


#### Cabello-Aguilar's Score (P-value)

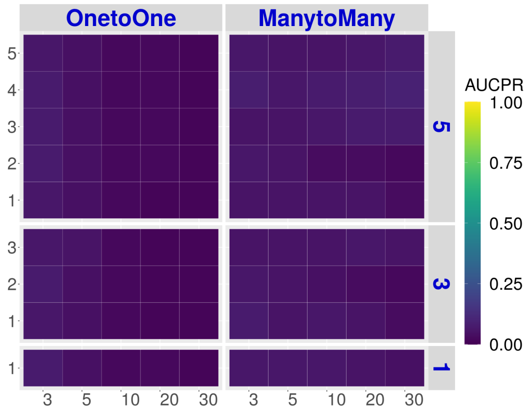


#### scTensor (NTD-2)

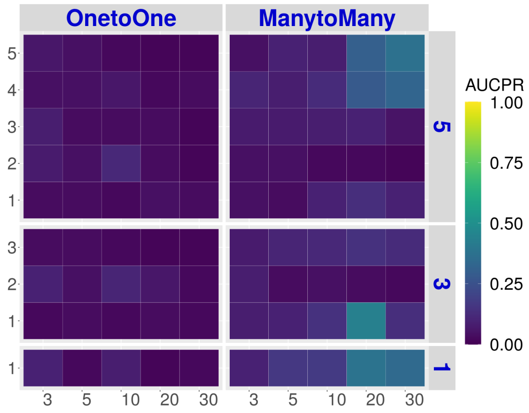

### E10 (Details)

The value ranges 0 to 1 (the closer to 1, the better)

#### Sum Score

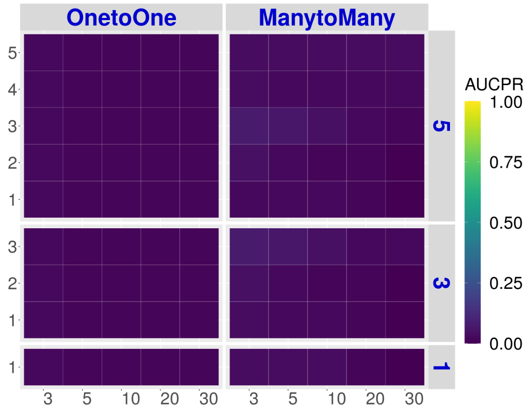


#### Product Score

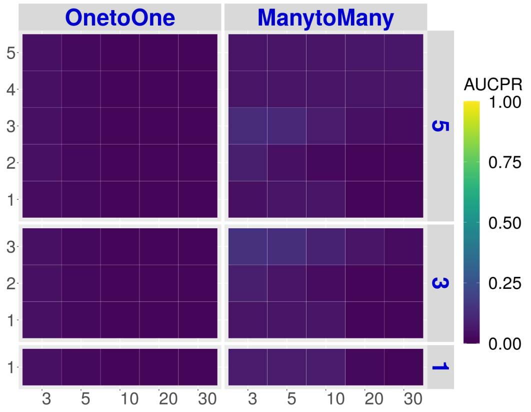


#### Halpern's Score

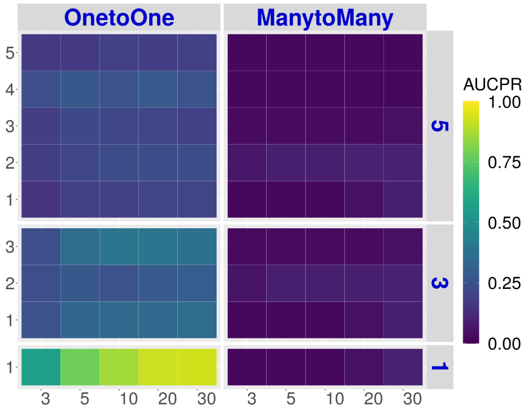


#### Cabello-Aguilar's Score

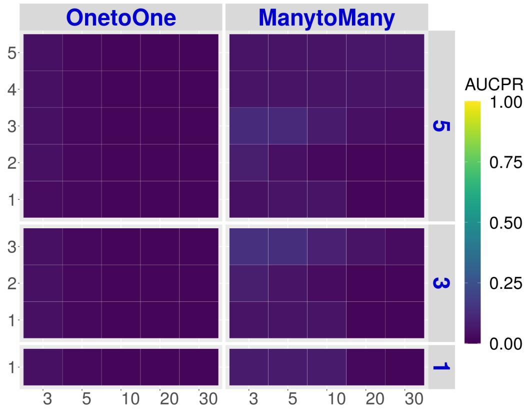


#### scTensor (NTD-3)

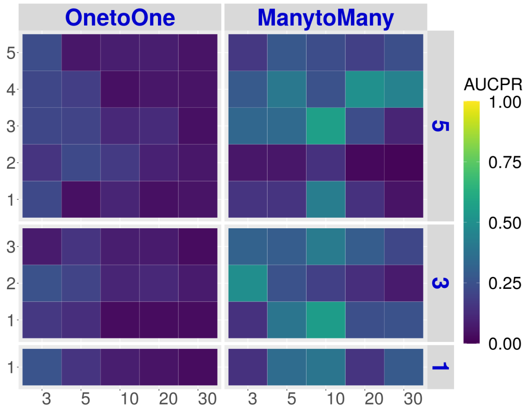


#### Sum Score (P-value)

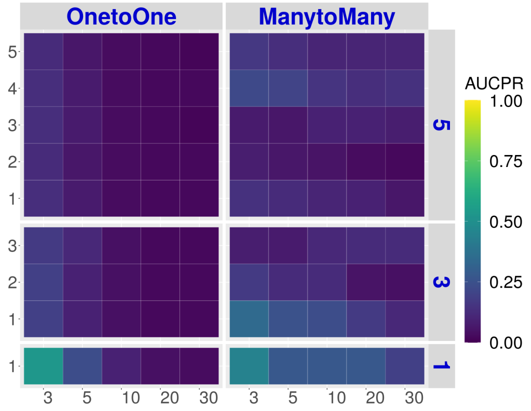


#### Product Score (P-value)

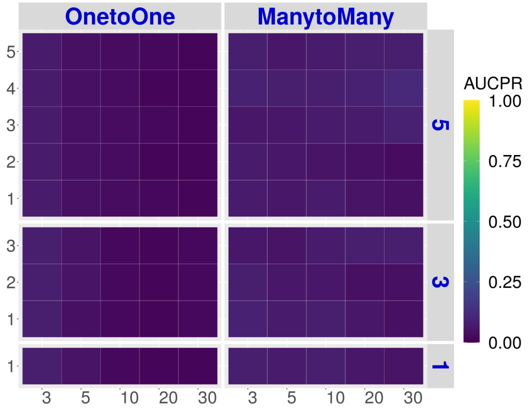


#### Halpern's Score (P-value)

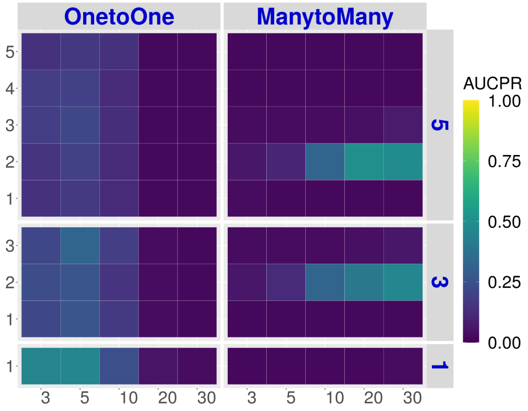


#### Cabello-Aguilar's Score (P-value)

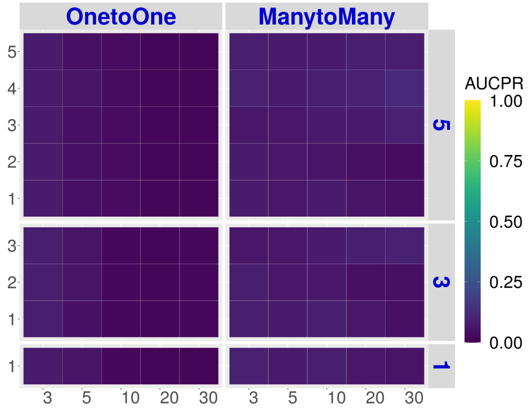


#### scTensor (NTD-2)

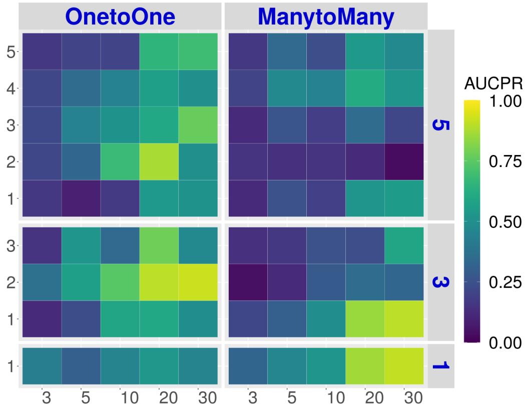

### Real Datasets

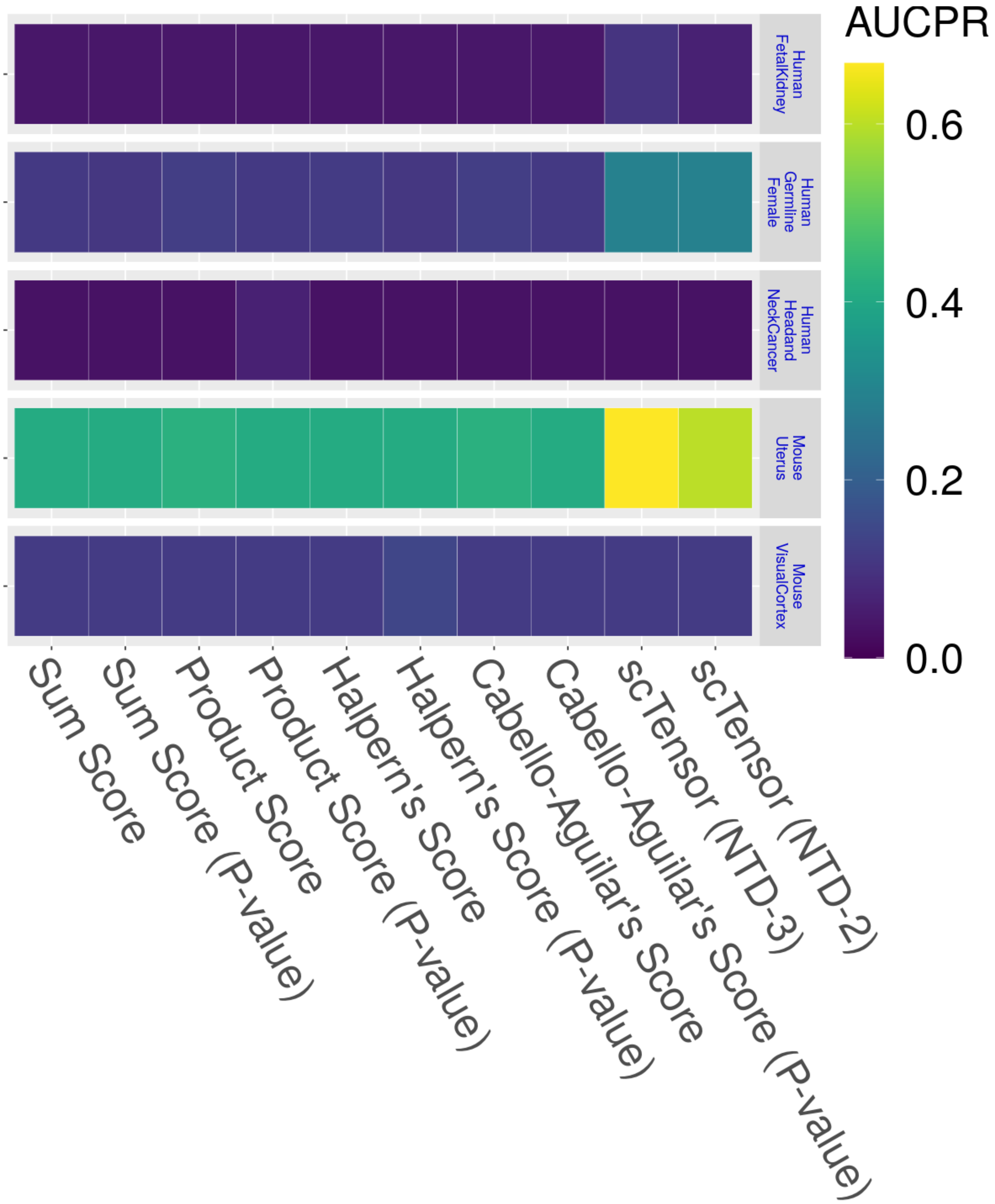

AUCPR
0.6
0.4
0.2
0.0
Human FetalKidney
Human Germline Female
Human Headand NeckCancer
Mouse Uterus
Mouse VisualCortex
Sum Score
Sum Score (P-value)
Product Score
Product Score (P-value)
Halpern's Score
Halpern's Score (P-value)
Cabello-Aguilar's Score
Cabello-Aguilar's Score (P-value)
scTensor (NTD-3)
scTensor (NTD-2)
