## Supplementary material for "scTensor detects many-to-many cell–cell interactions from single cell RNA-sequencing data": https://zenodo.org/record/7412280/files/05_Memory.pdf?download=1

### Simulated Datasets

**E2 (Summary)** The larger, the worse

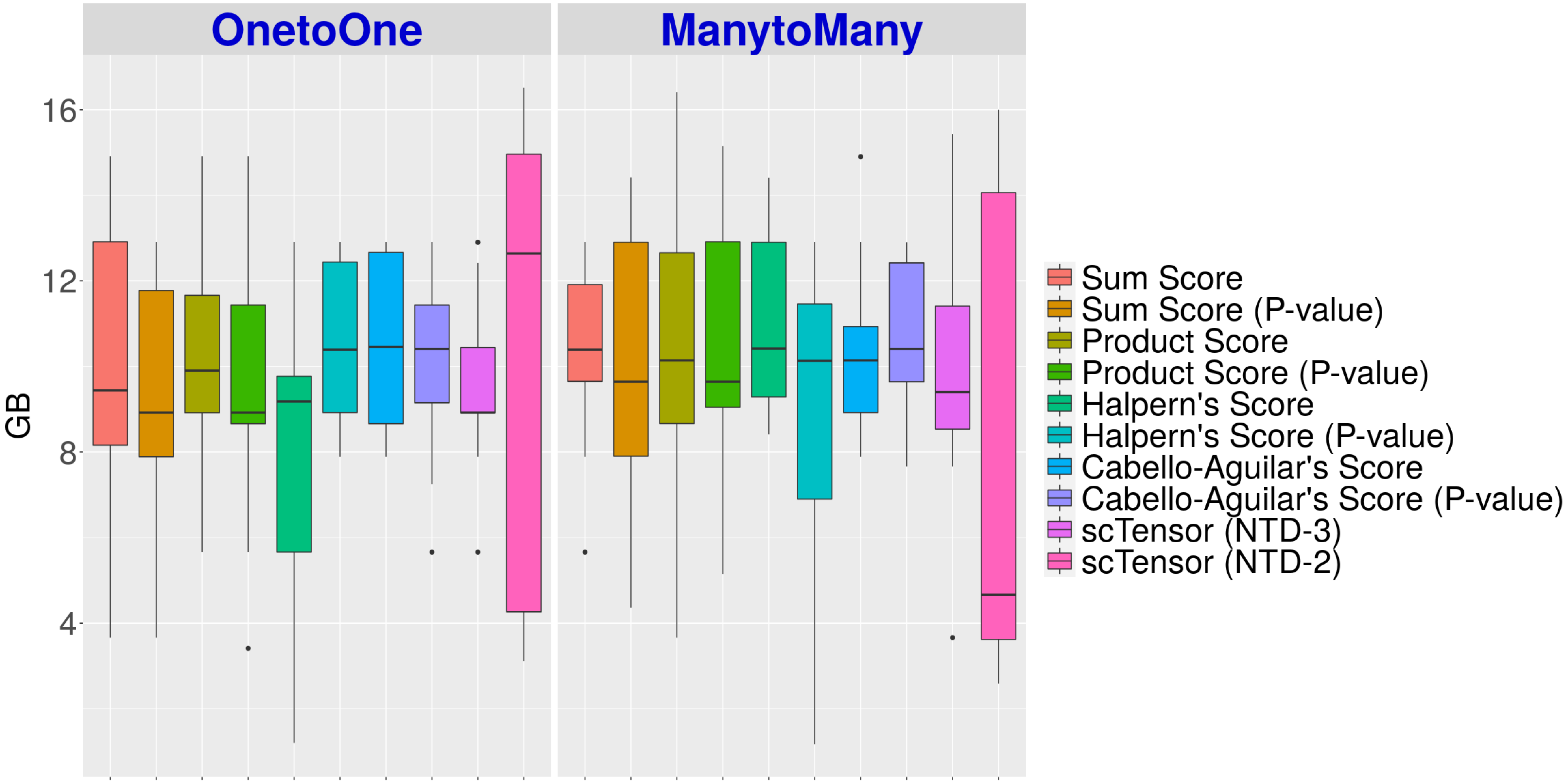

**E5 (Summary)** The larger, the worse

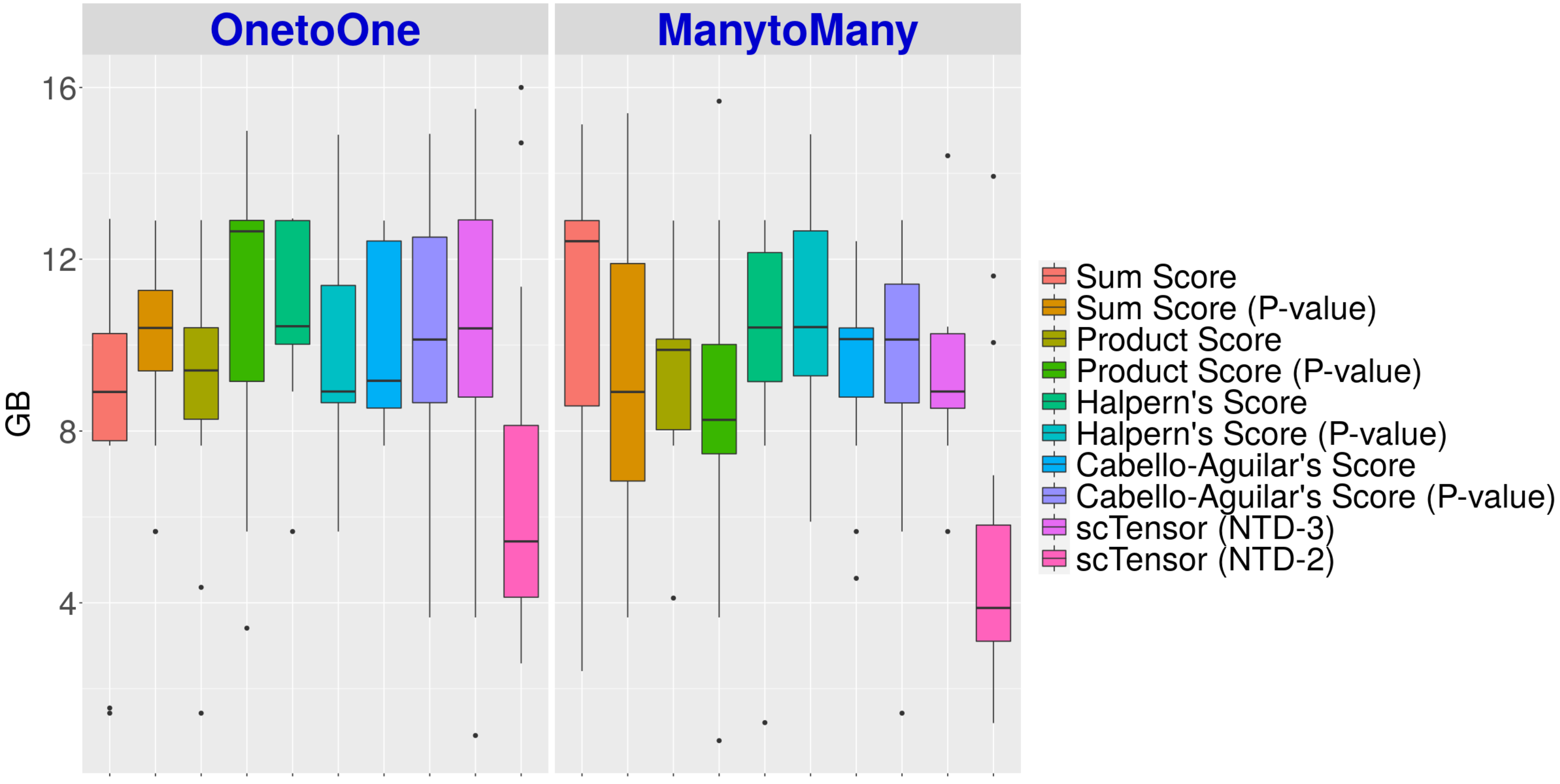

E10 (Summary) The larger, the worse

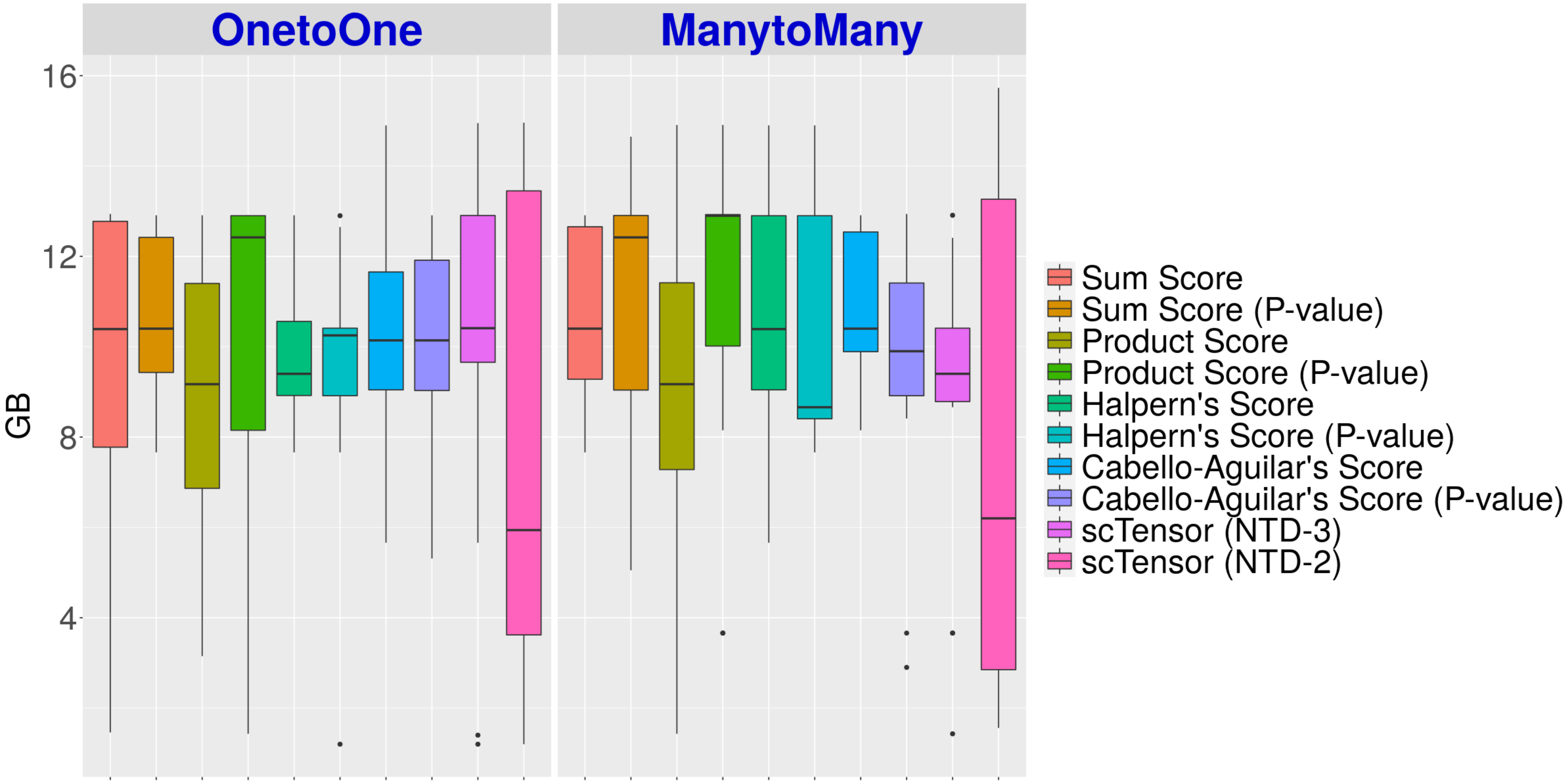

### E2 (Details)

The larger, the worse

#### Sum Score

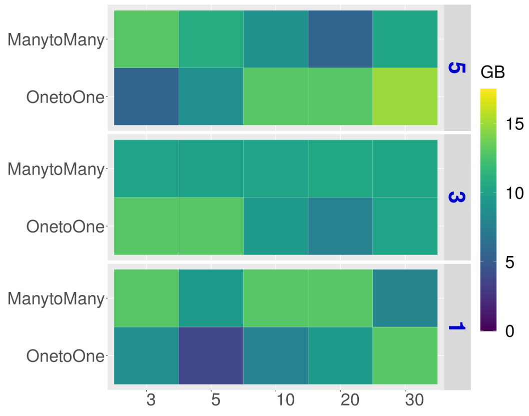


#### Product Score

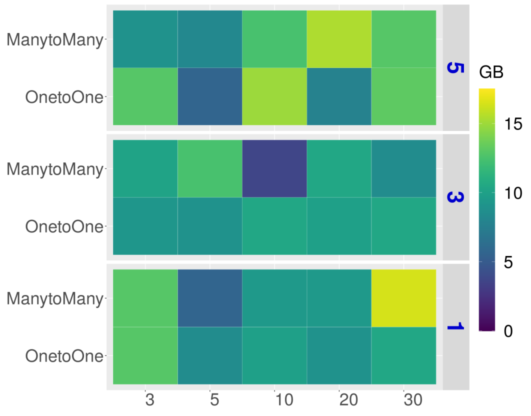


#### Halpern's Score

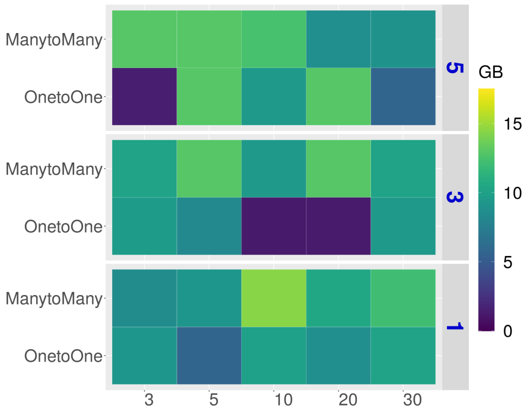


#### Cabello-Aguilar's Score

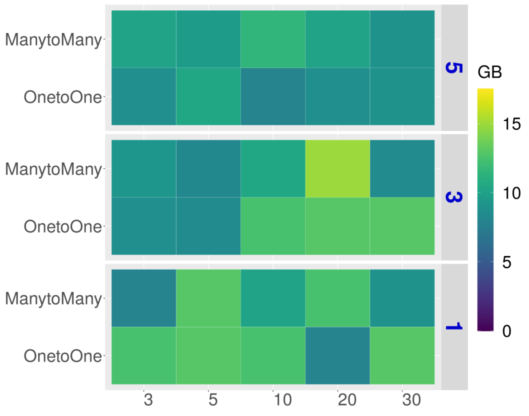


#### scTensor (NTD-3)

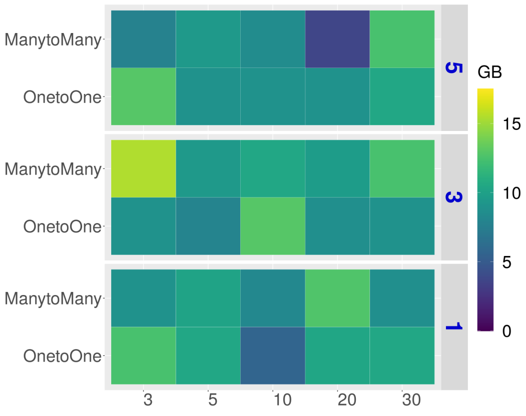


#### Sum Score (P-value)

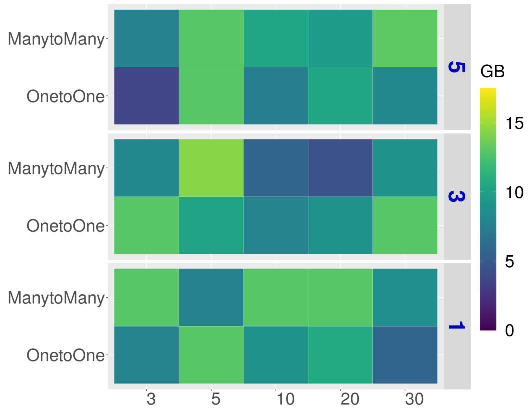


#### Product Score (P-value)

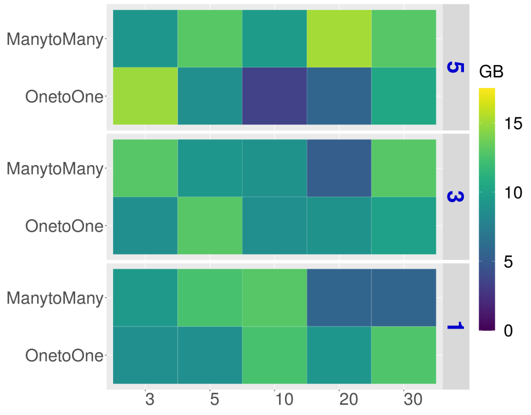


#### Halpern's Score (P-value)

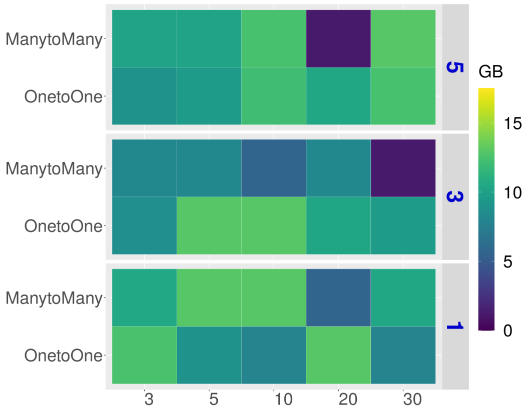


#### Cabello-Aguilar's Score (P-value)

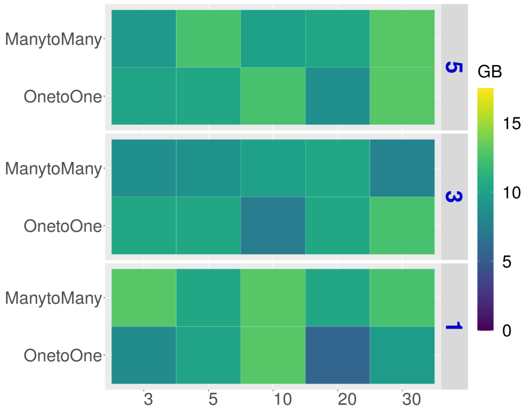


#### scTensor (NTD-2)

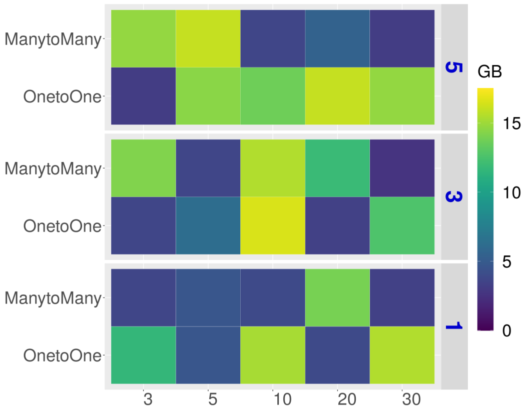

### E5 (Details)

The larger, the worse

#### Sum Score

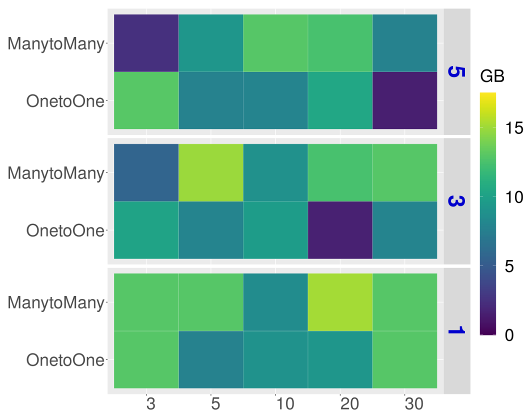


#### Product Score

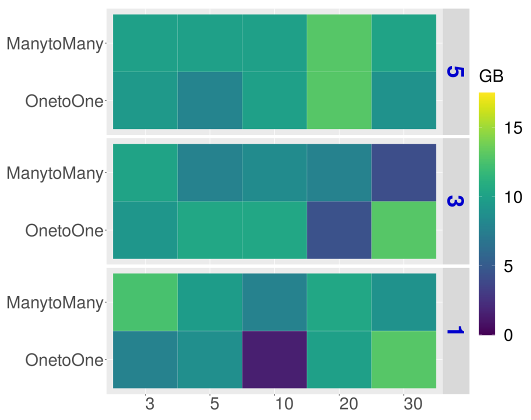


#### Halpern's Score

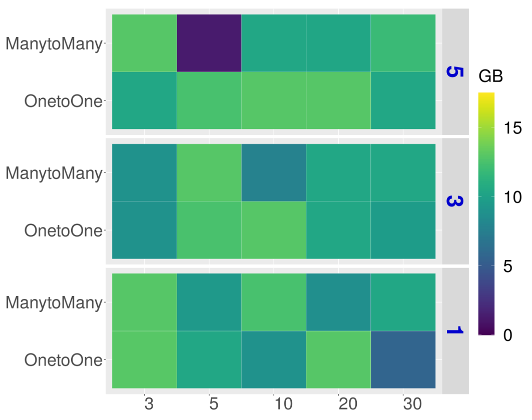


#### Cabello-Aguilar's Score

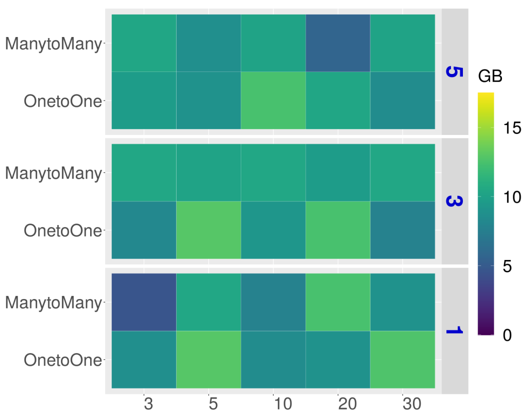


#### scTensor (NTD-3)

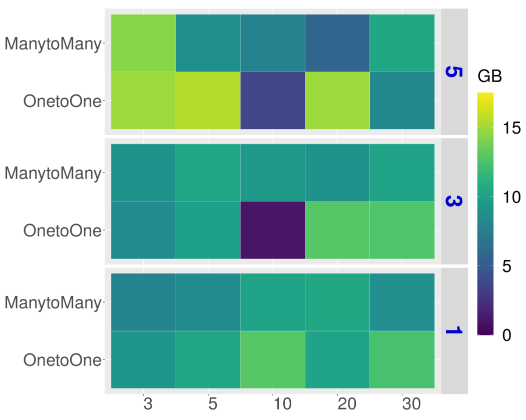


#### Sum Score (P-value)

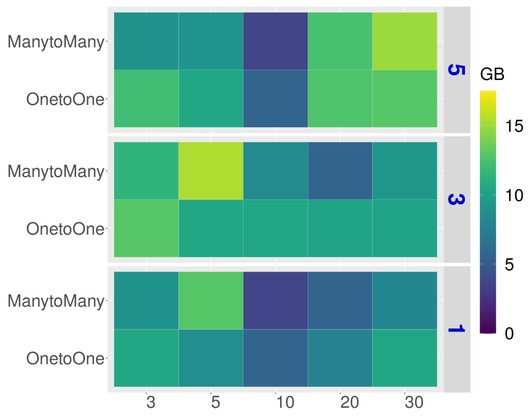


#### Product Score (P-value)

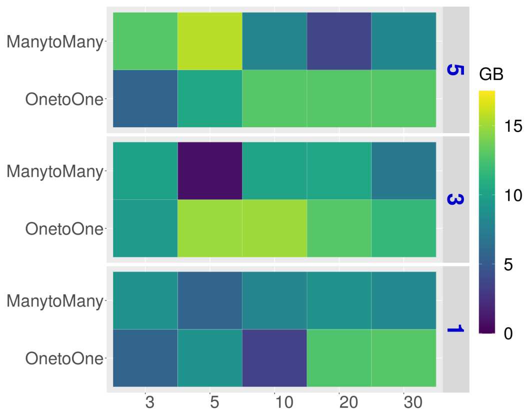


#### Halpern's Score (P-value)

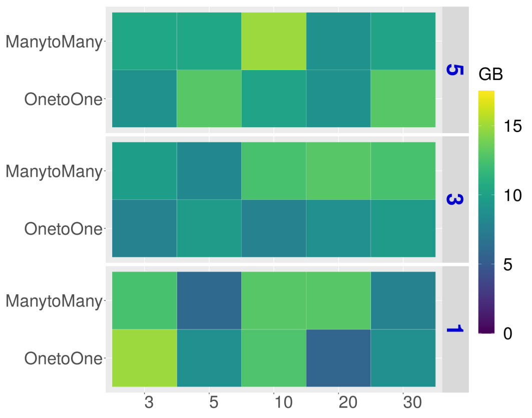


#### Cabello-Aguilar's Score (P-value)

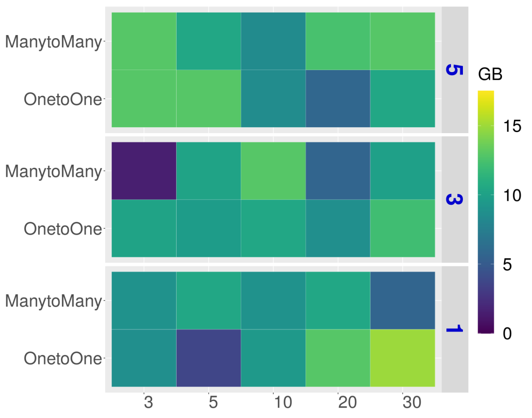


#### scTensor (NTD-2)

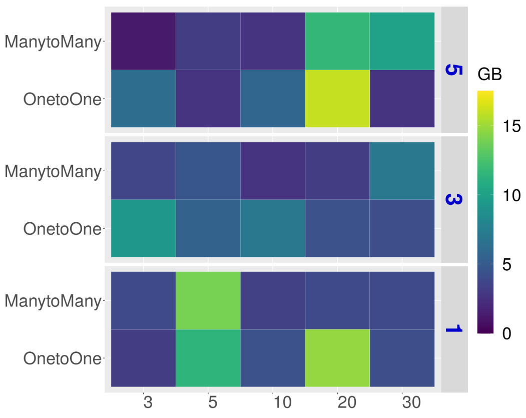

### E10 (Details)

The larger, the worse

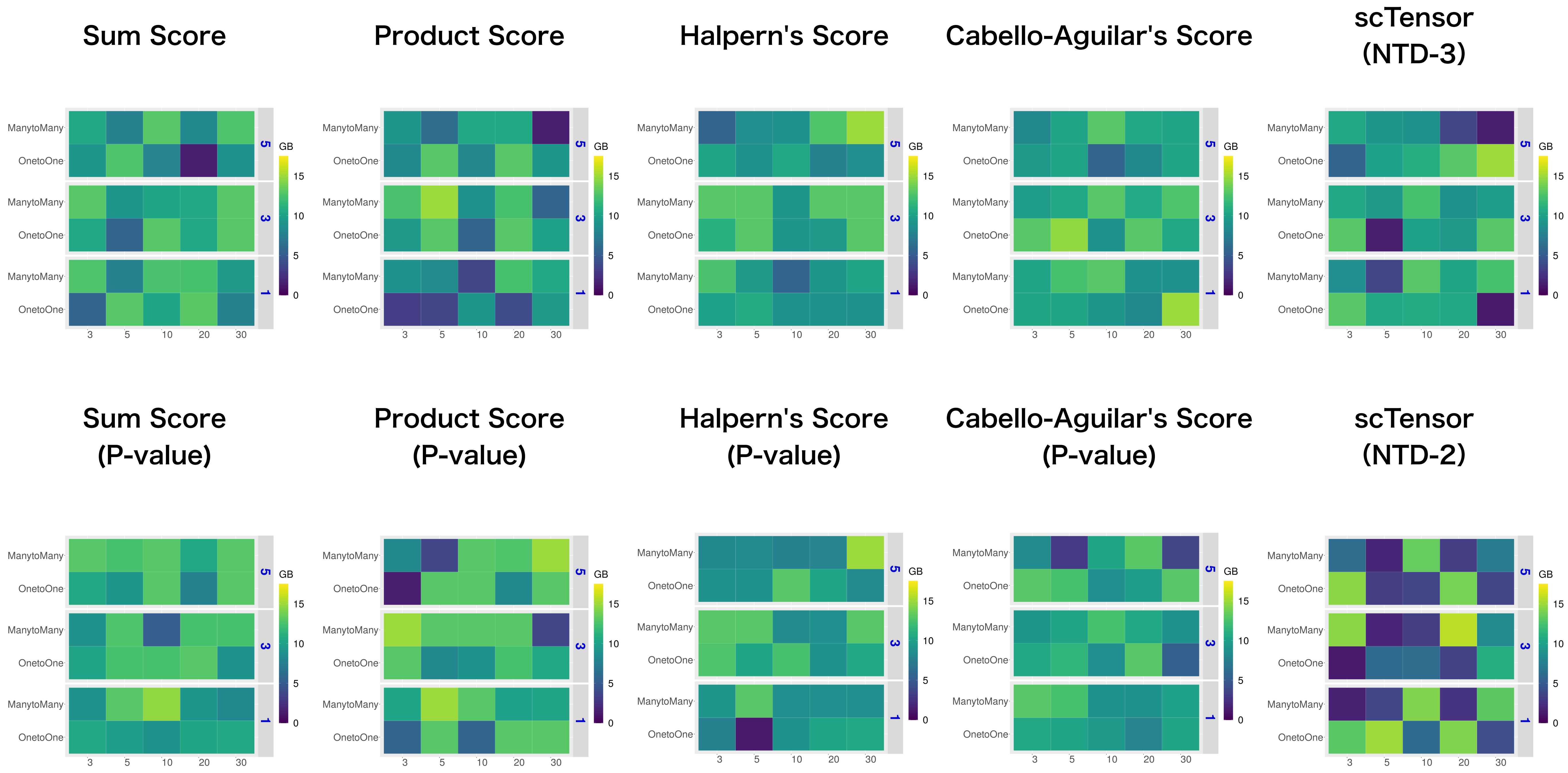

### Real Datasets

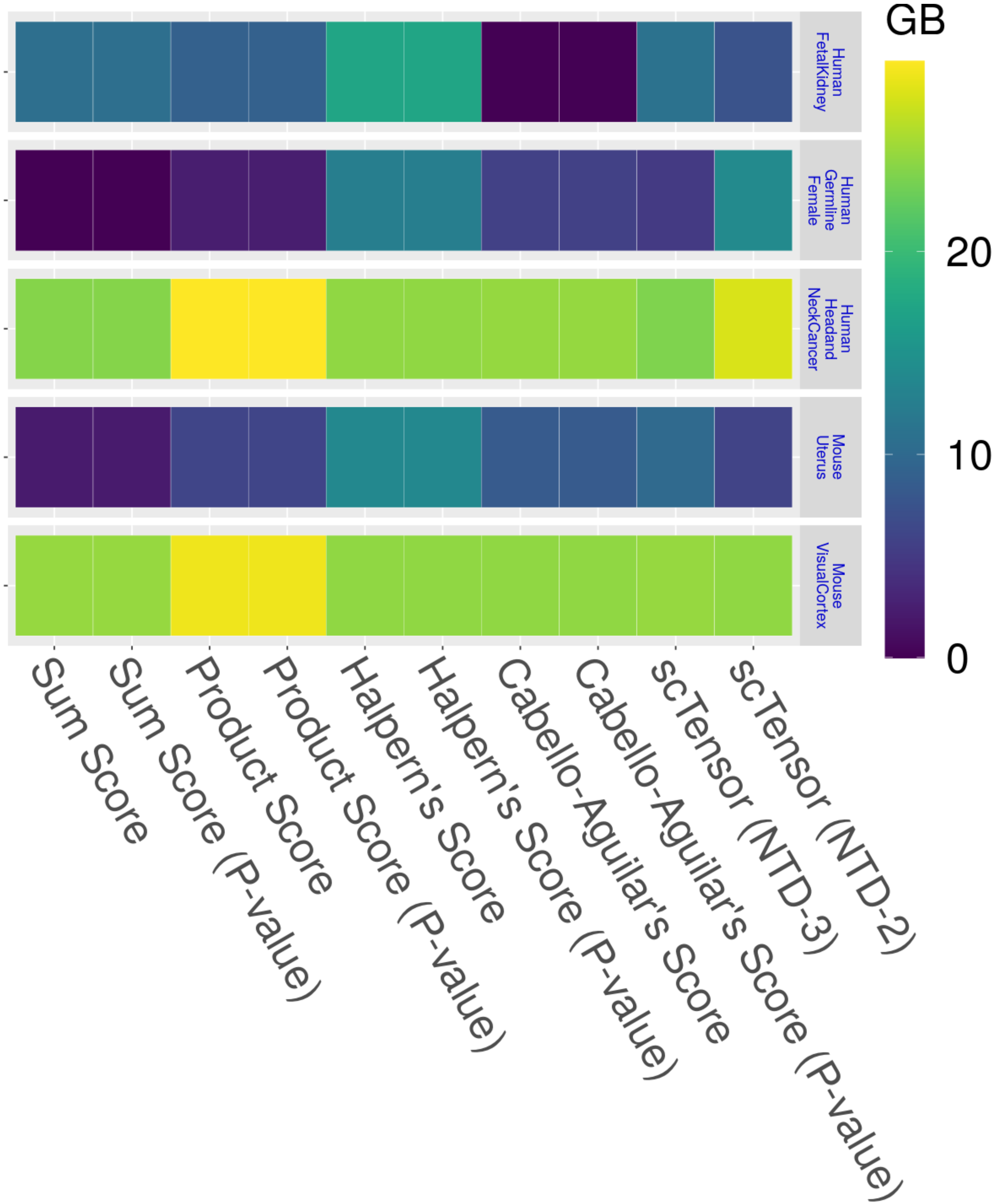

GB
20
10
0
Human FetalKidney
Human Germline Female
Human Headand NeckCancer
Mouse Uterus
Mouse VisualCortex
Sum Score
Sum Score (P-value)
Product Score
Product Score (P-value)
Halpern's Score
Halpern's Score (P-value)
Cabello-Aguilar's Score
Cabello-Aguilar's Score (P-value)
scTensor (NTD-3)
scTensor (NTD-2)
