## Supplementary material for "scTensor detects many-to-many cell–cell interactions from single cell RNA-sequencing data": https://zenodo.org/record/7412280/files/07_F-measure.pdf?download=1

### Simulated Datasets

**E2 (Summary)** The value ranges 0 to 1 (the closer to 1, the better)

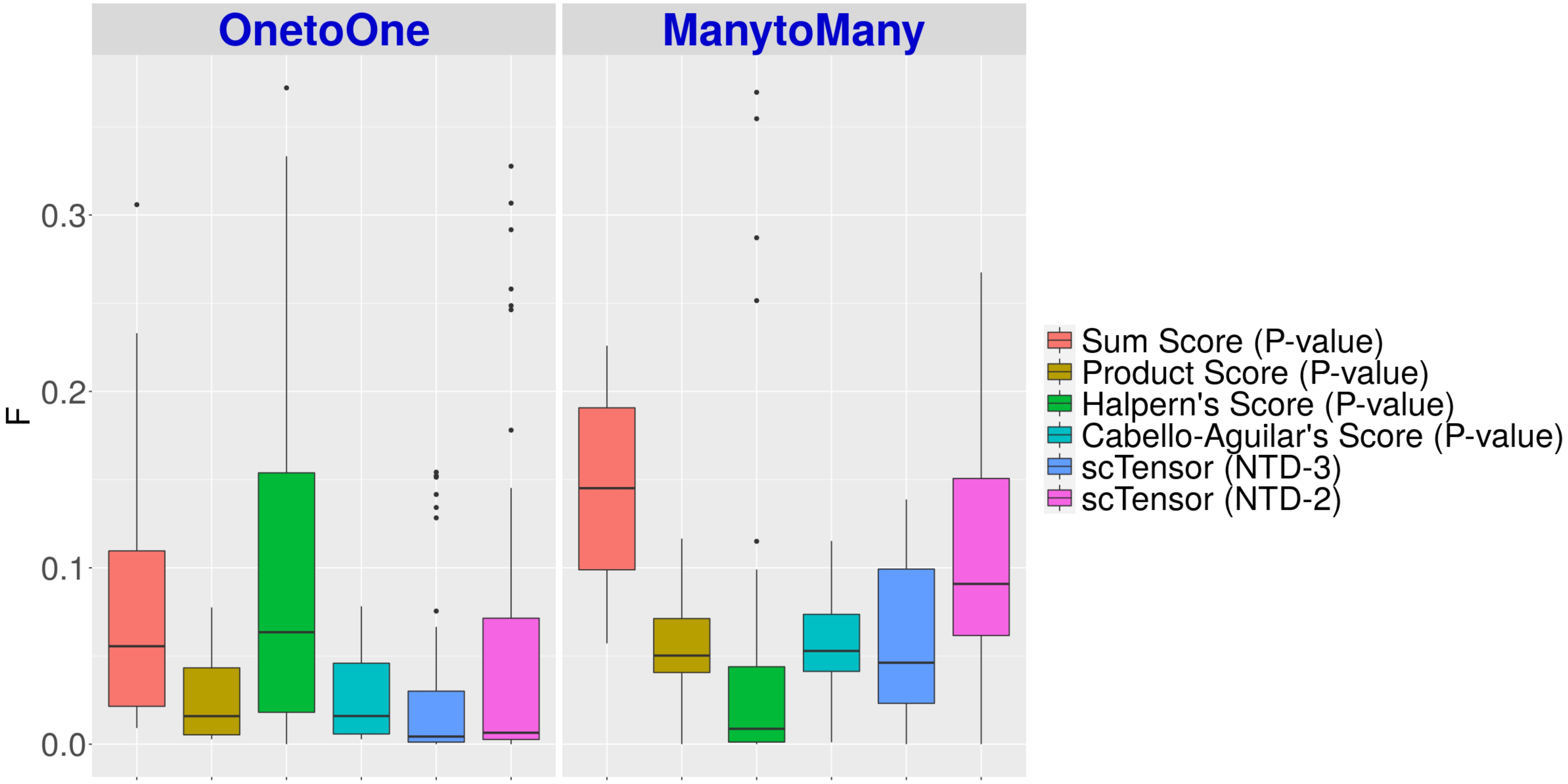

E5 (Summary) The value ranges 0 to 1 (the closer to 1, the better)

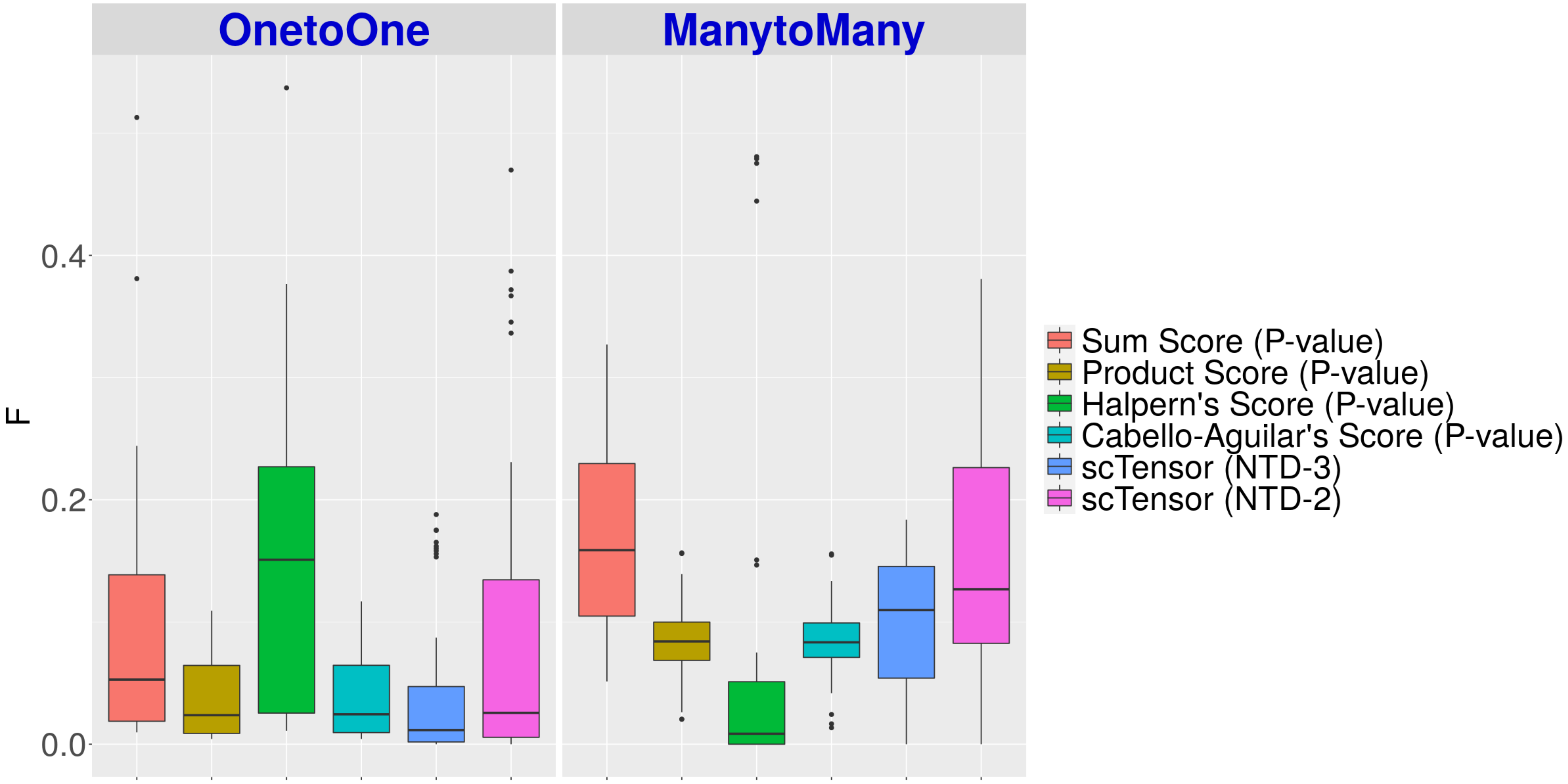

E10 (Summary) The value ranges 0 to 1 (the closer to 1, the better)

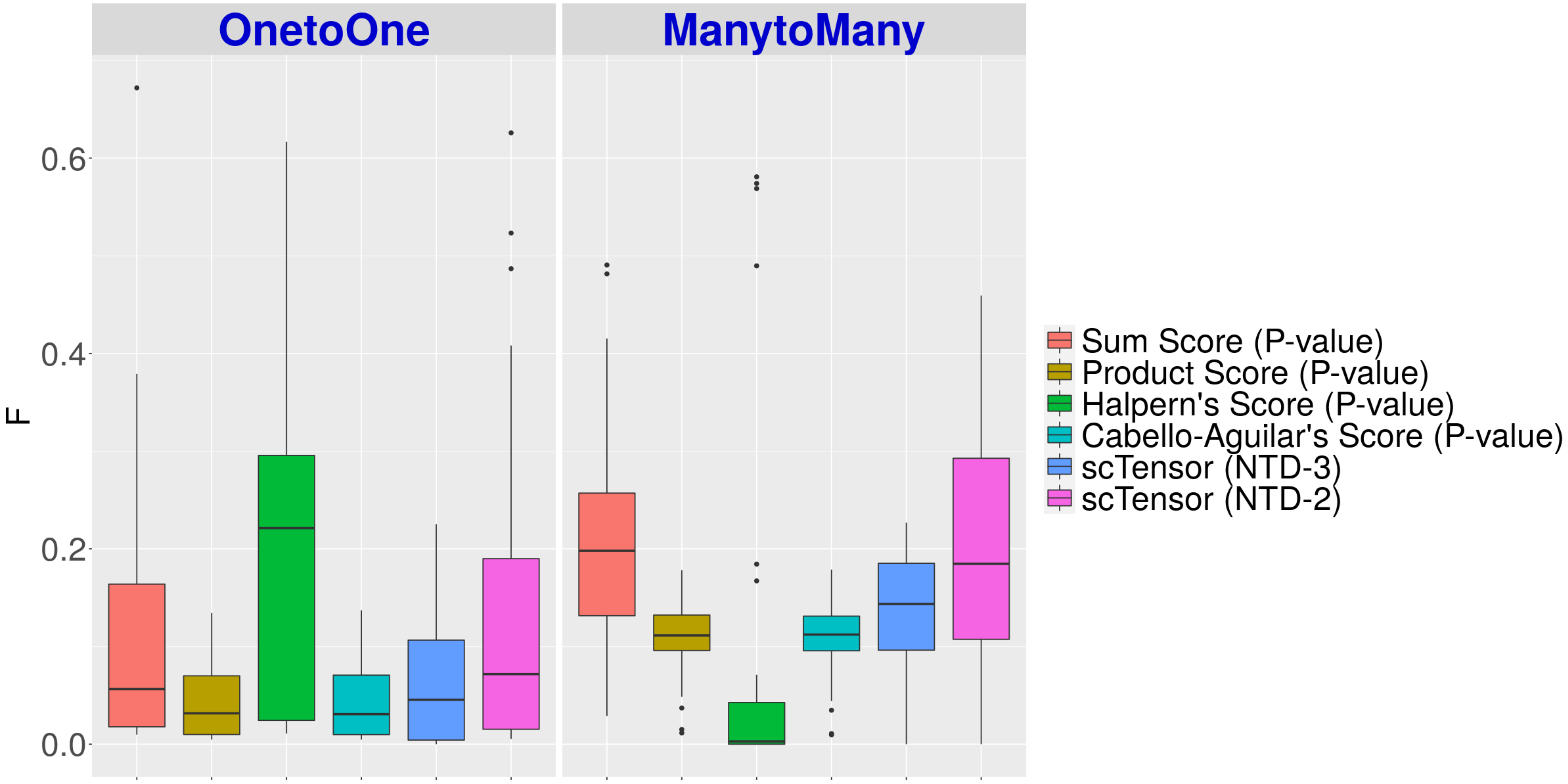

#### E2 (Details)

The value ranges 0 to 1 (the closer to 1, the better)

These may only work in certain cases
but overall these do not work well.

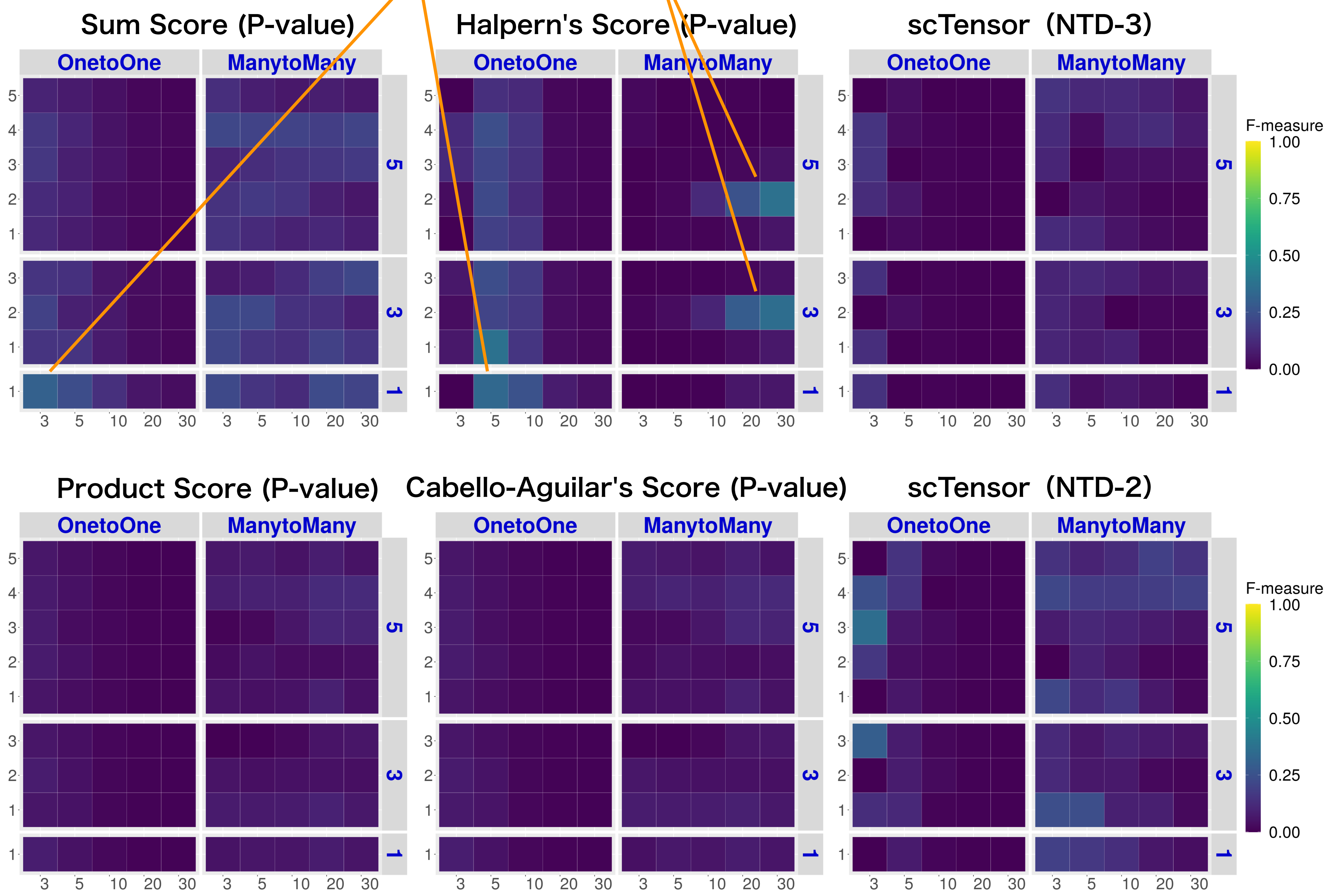


Stable in both 1vs1
and many vs many

### E5 (Details)

The value ranges 0 to 1 (the closer to 1, the better)

These may only work in certain cases
but overall these do not work well.

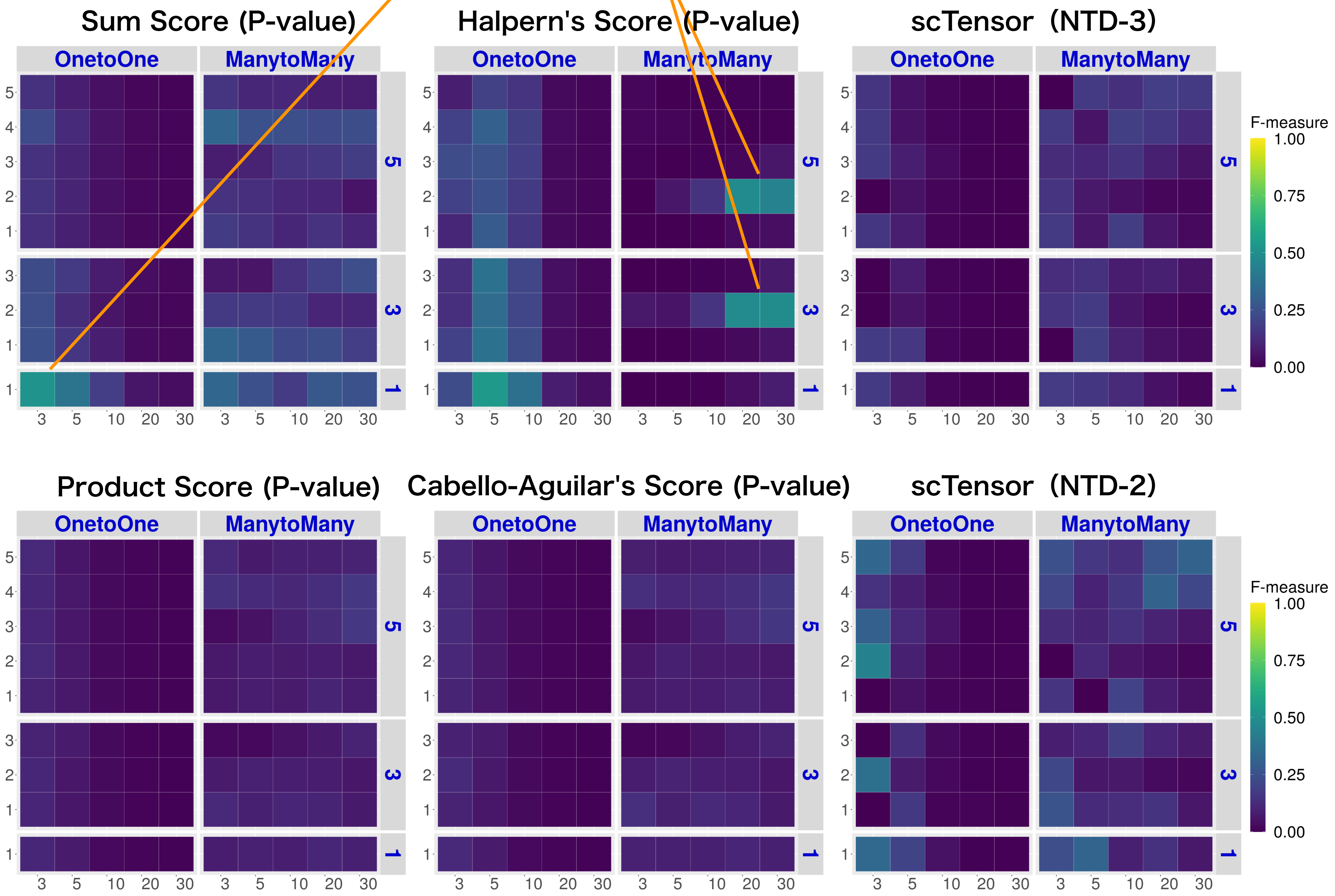


Stable in both 1vs1
and many vs many

### E10 (Details)

The value ranges 0 to 1 (the closer to 1, the better)

These may only work in certain cases
but overall these do not work well.

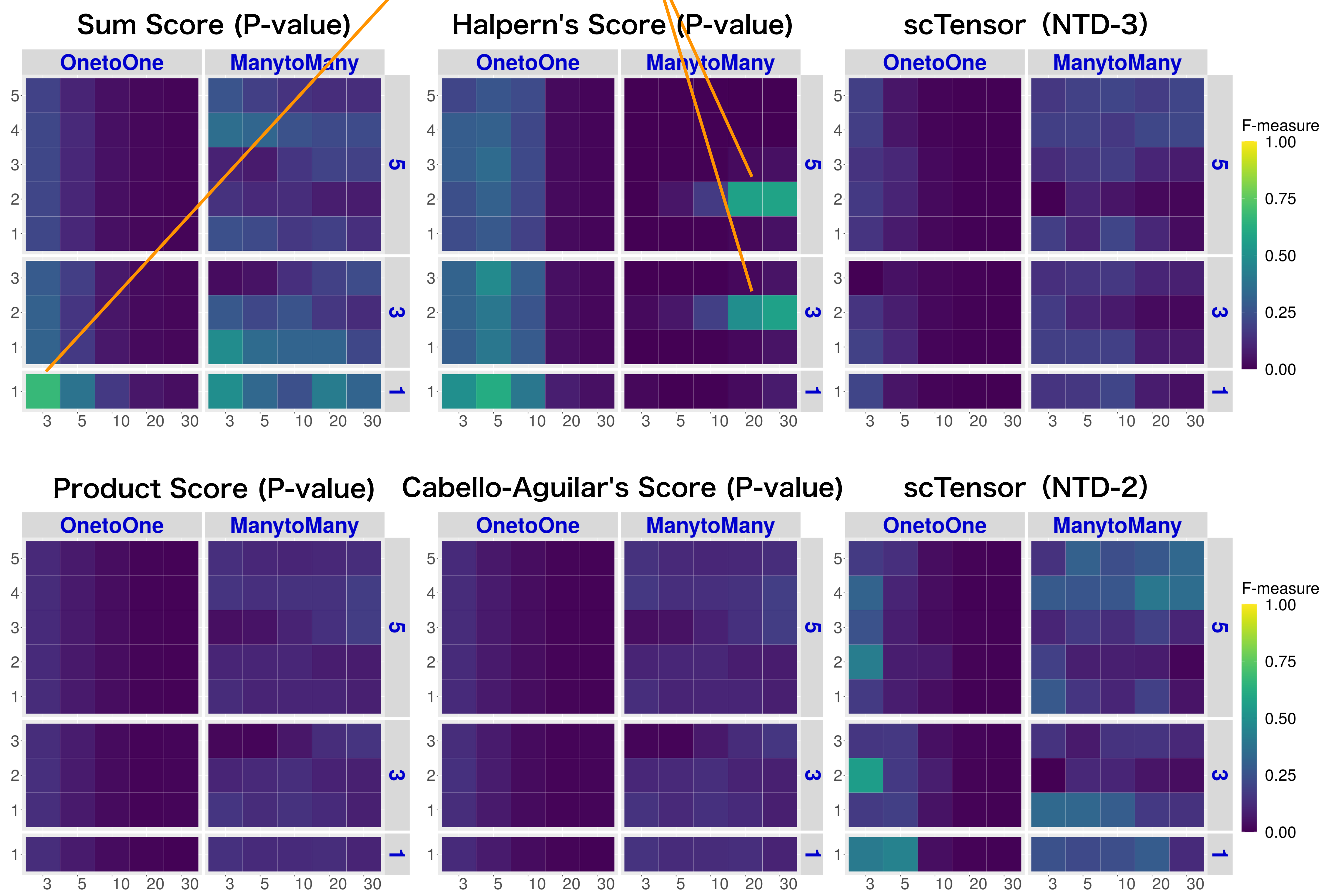


Stable in both 1vs1
and many vs many

### Real Datasets

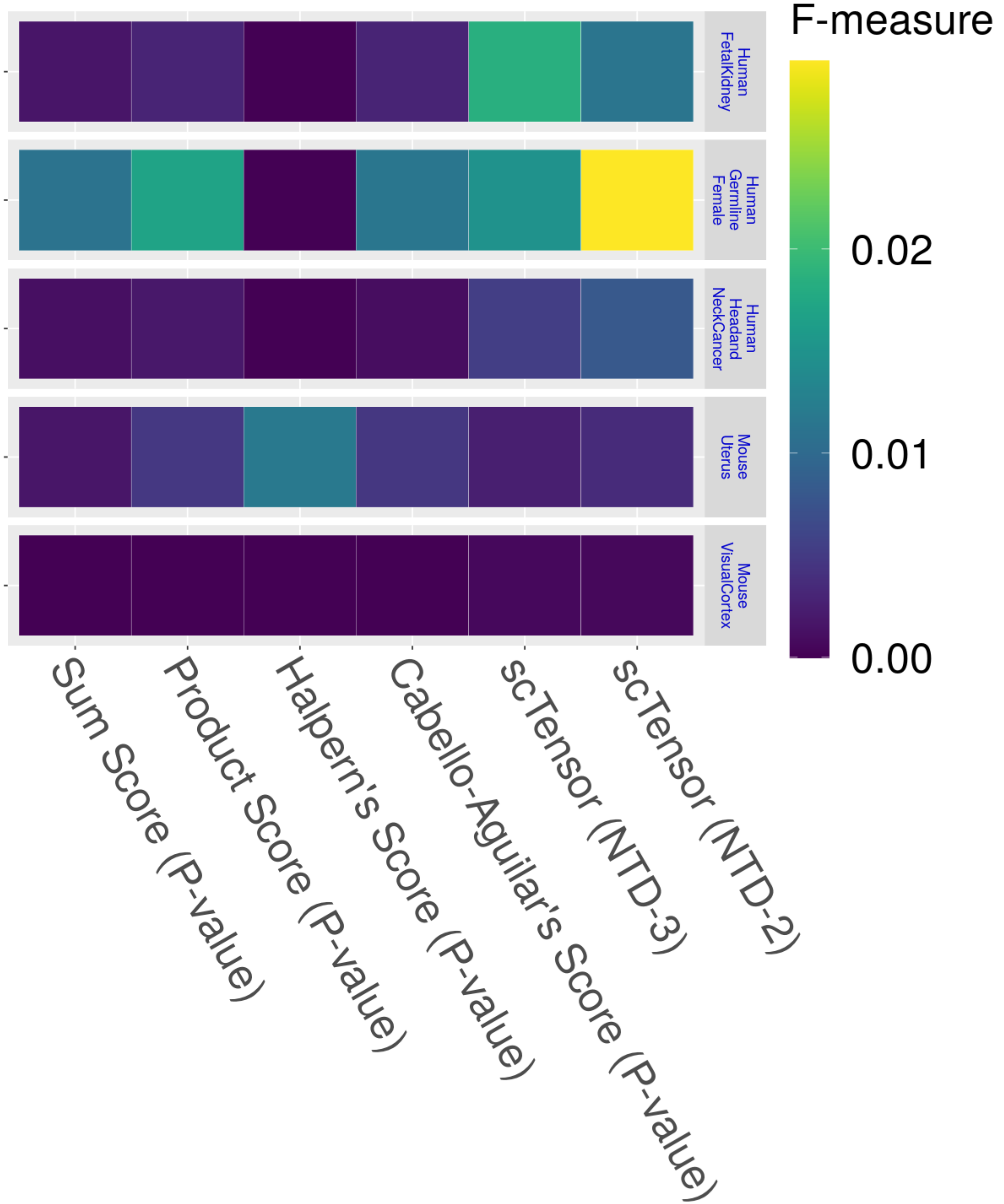
F-measure
0.02
0.01
0.00
Human FetalKidney
Human Germline Female
Human Headand NeckCancer
Mouse Uterus
Mouse VisualCortex
Sum Score (P-value)
Product Score (P-value)
Halpern's Score (P-value)
Cabello-Aguilar's Score (P-value)
scTensor (NTD-3)
scTensor (NTD-2)
