## Supplementary figures and images for "scTensor detects many-to-many cell–cell interactions from single cell RNA-sequencing data"

### https://zenodo.org/record/7412280/files/09_PR.pdf?download=1

#### Simulated Datasets

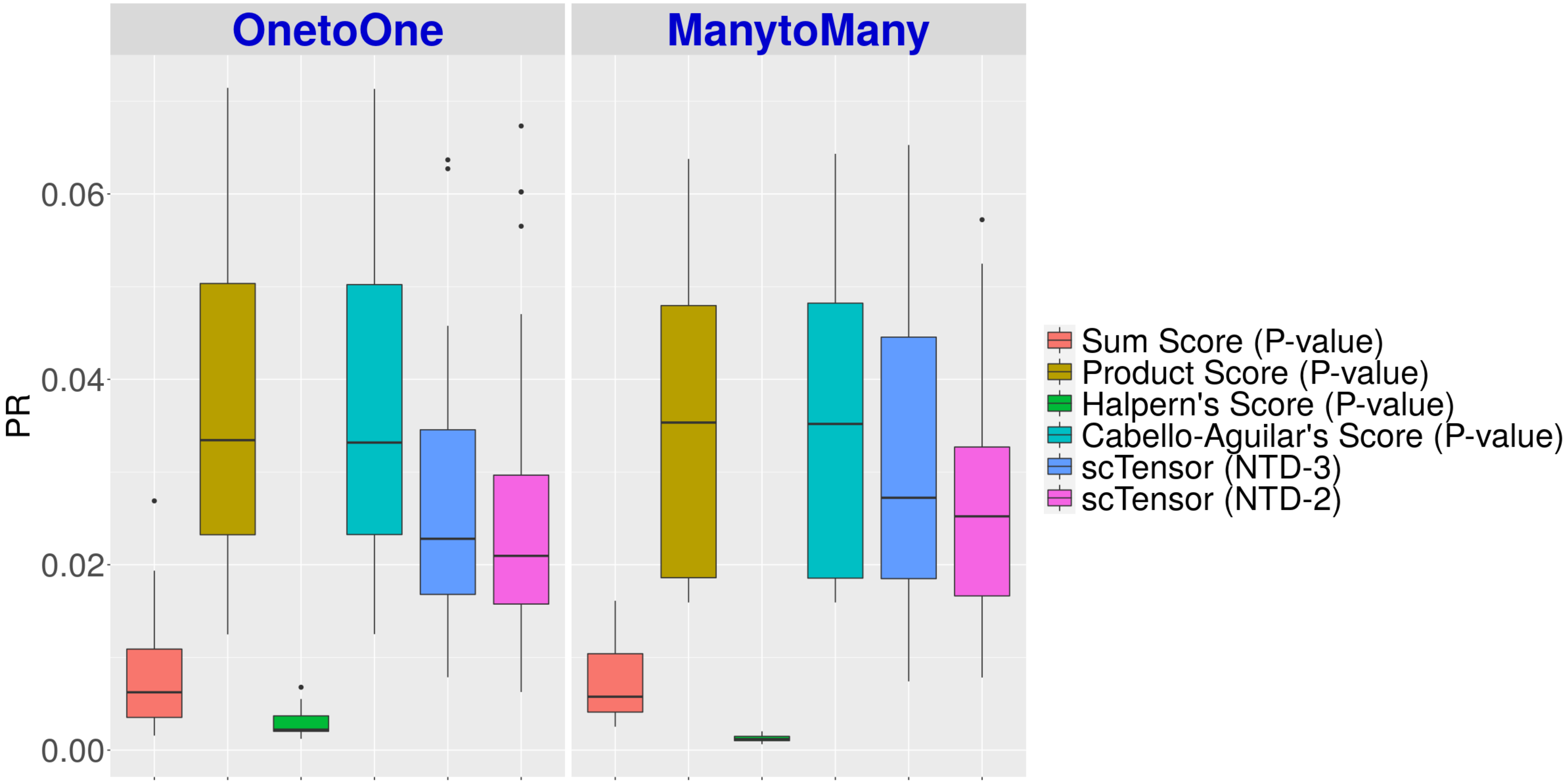

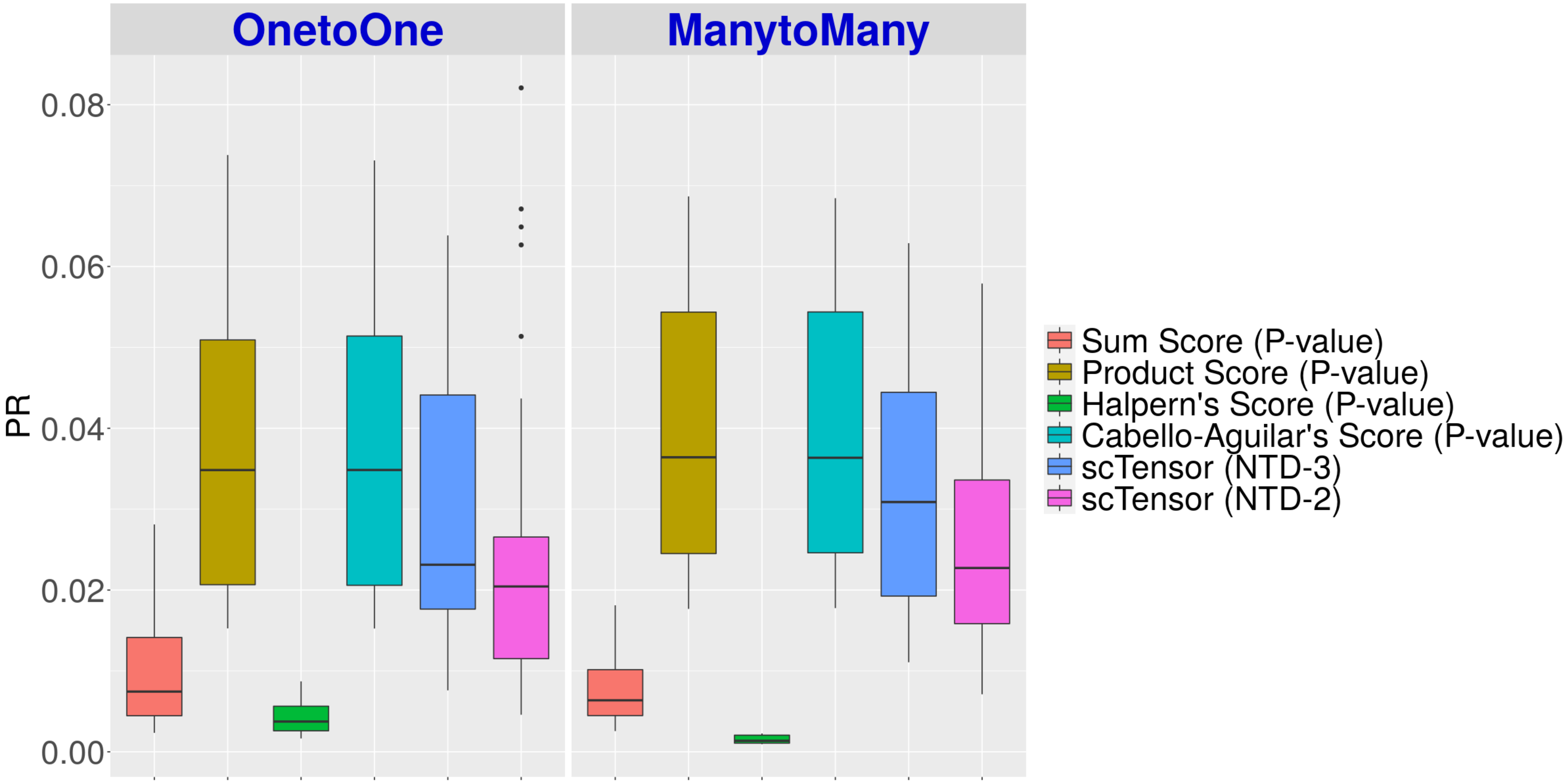

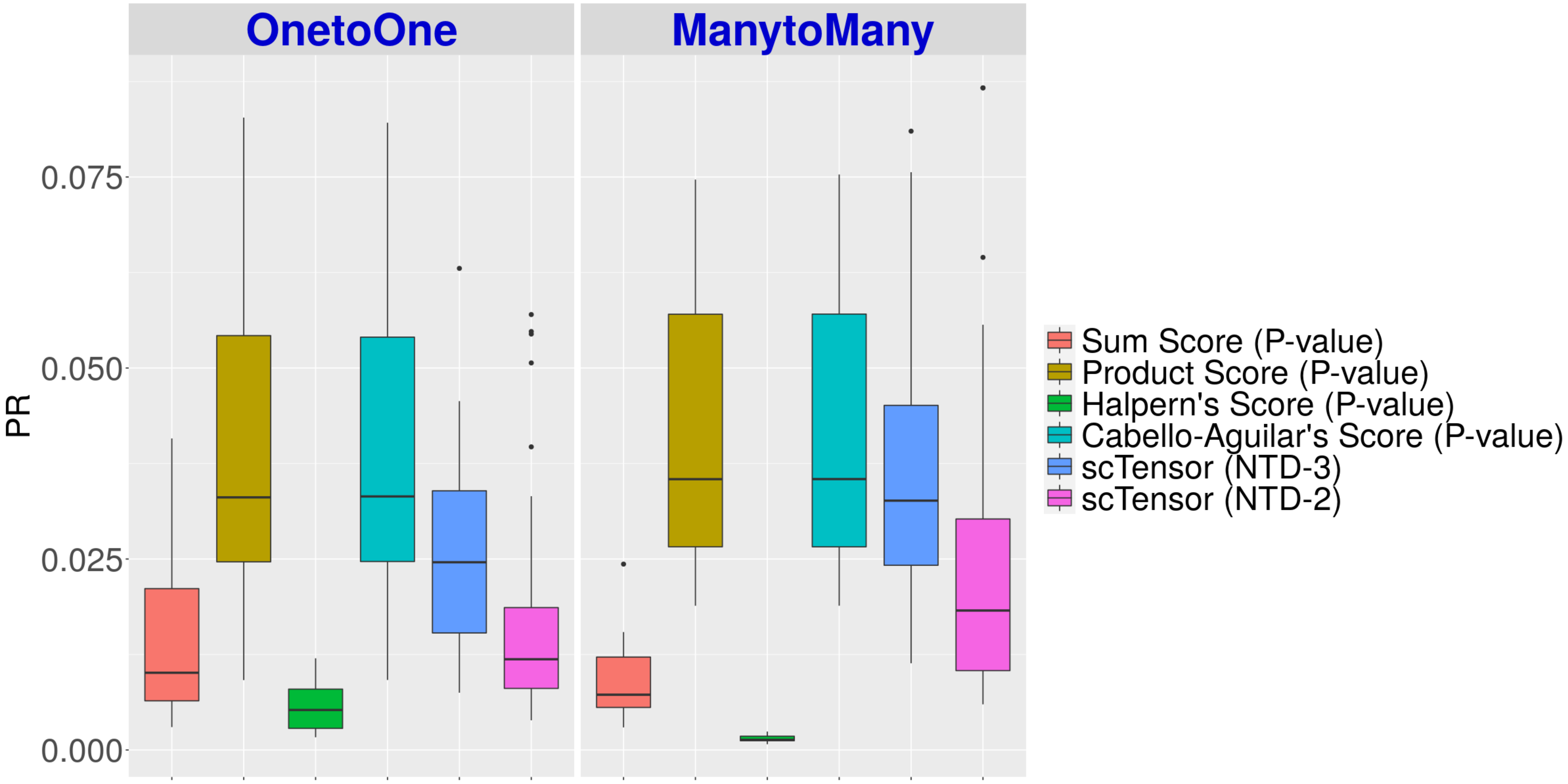

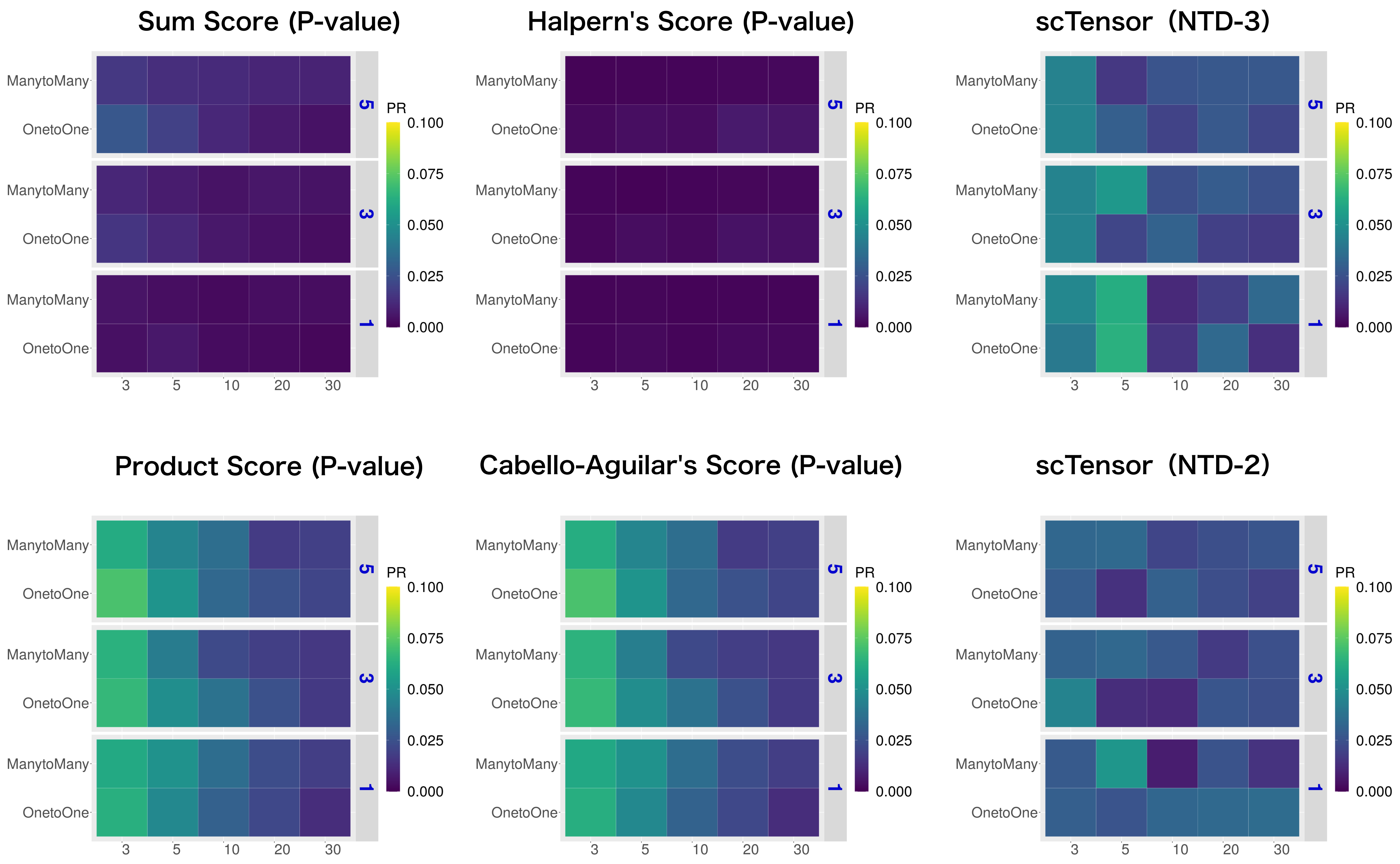

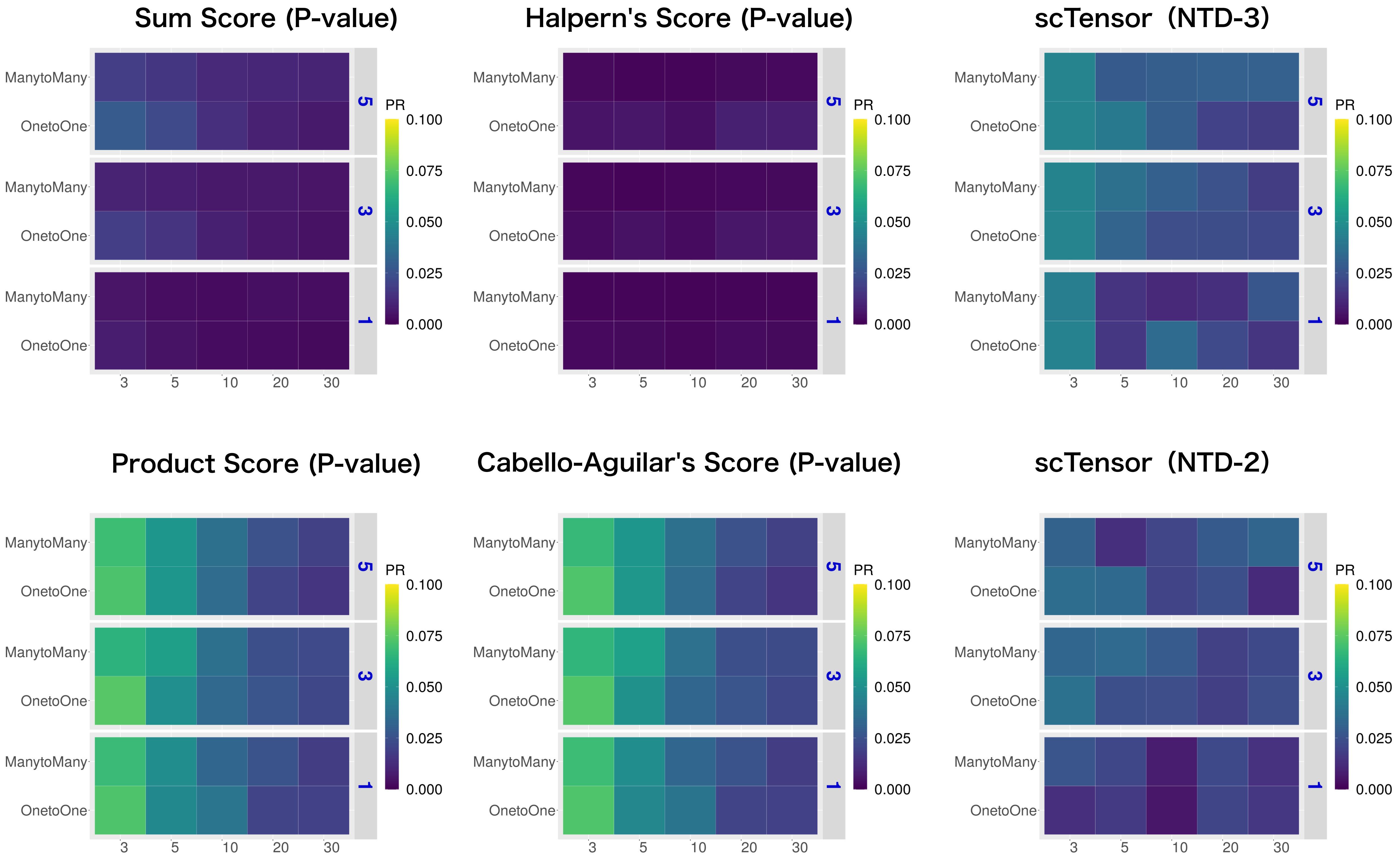

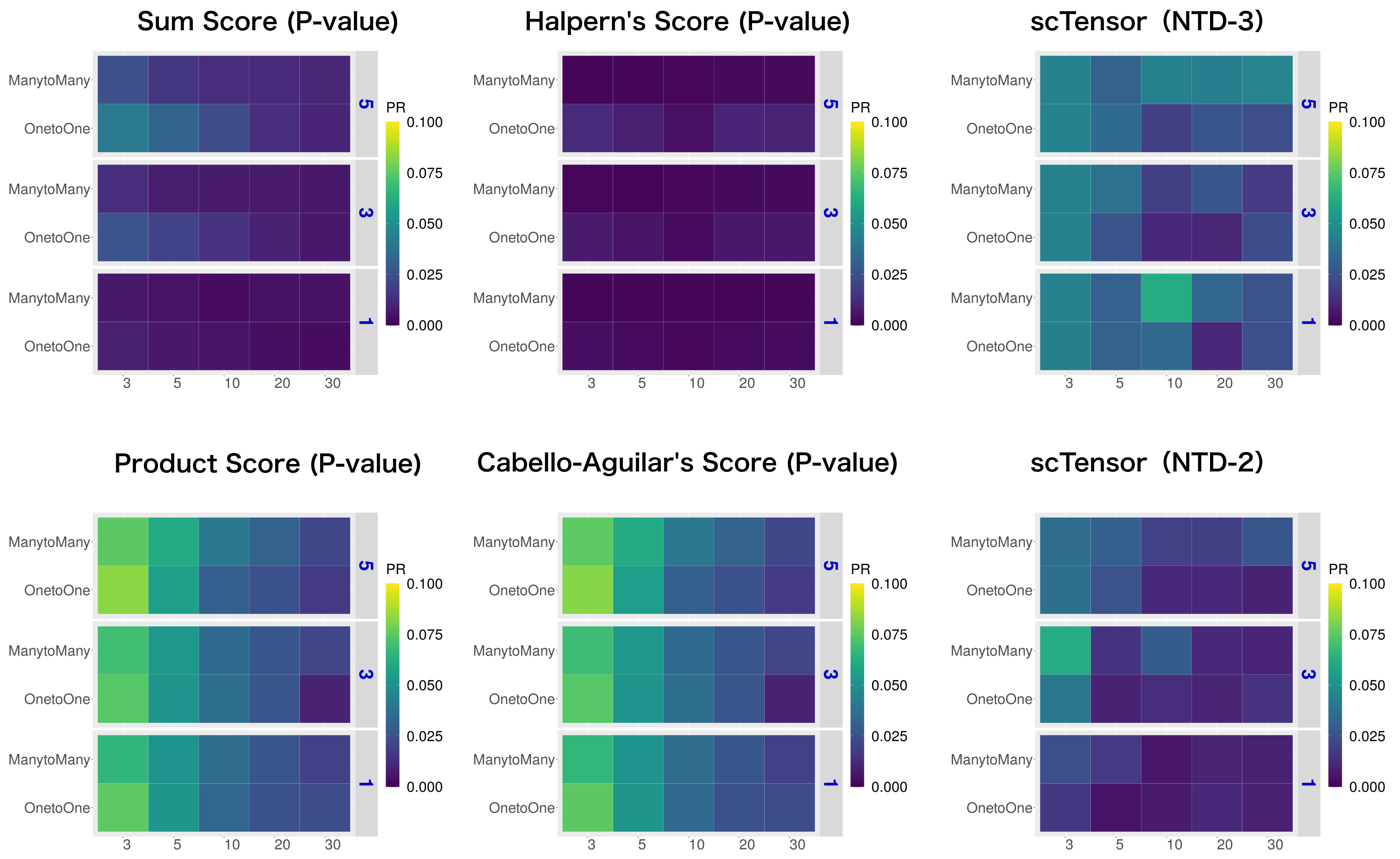

#### Real Datasets

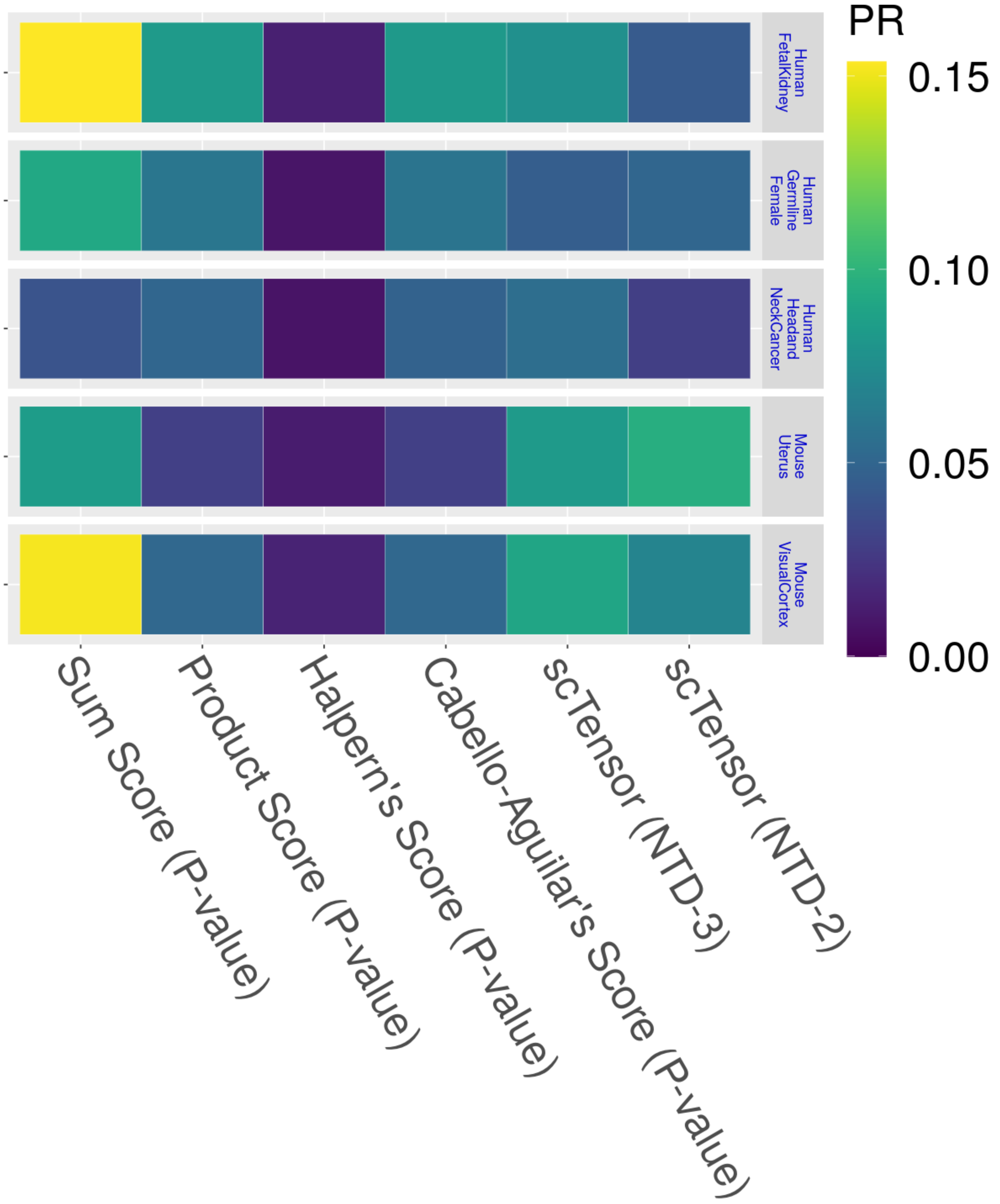

### https://zenodo.org/record/7412280/files/12_TR.pdf?download=1

#### Simulated Datasets

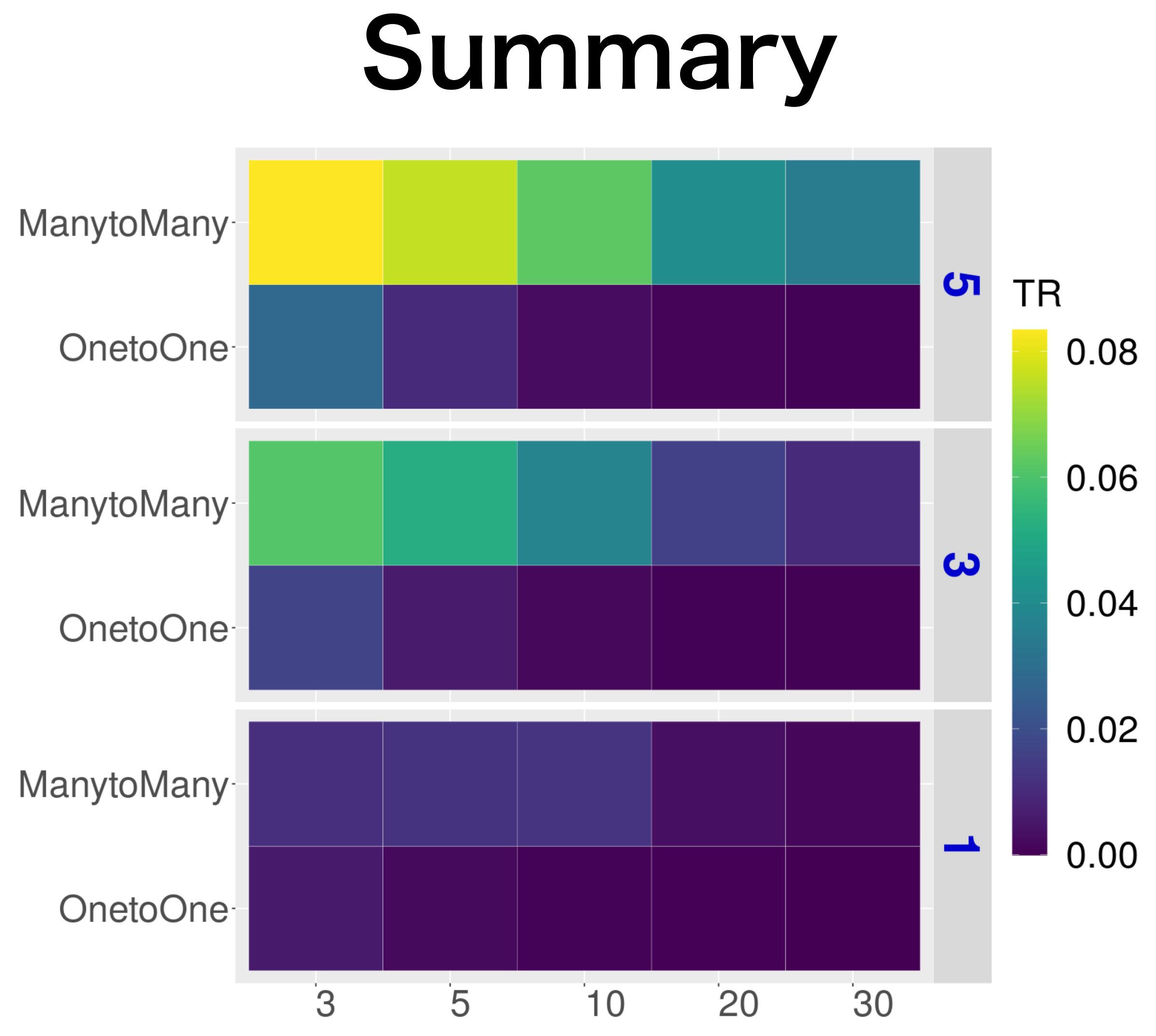

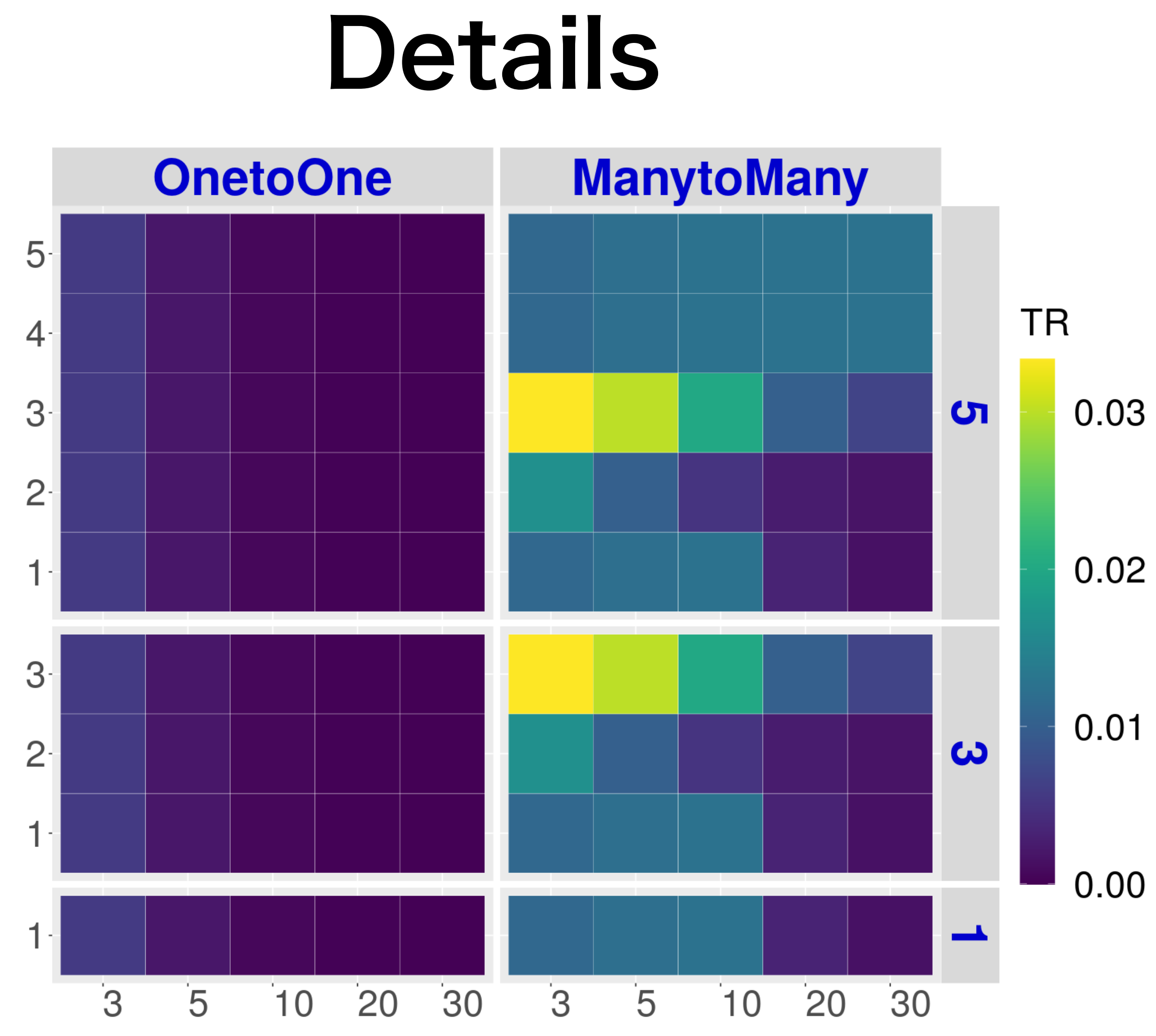

#### Real Datasets

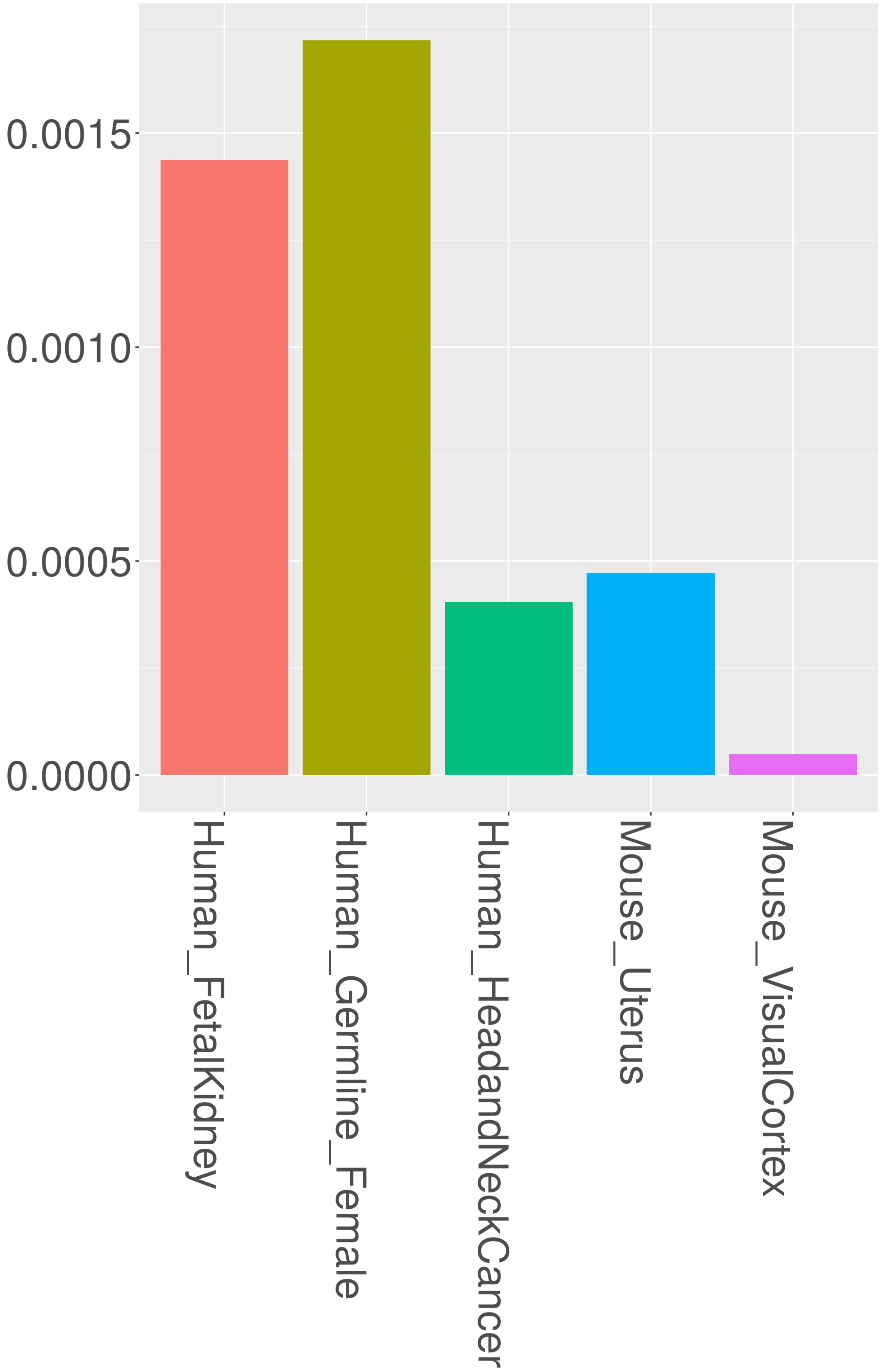
