## Supplementary material for "scTensor detects many-to-many cell–cell interactions from single cell RNA-sequencing data": https://zenodo.org/record/7412280/files/11_FNR.pdf?download=1

### Simulated Datasets

**E2 (Summary)** The value ranges 0 to 1 (the closer to 1, the worse)

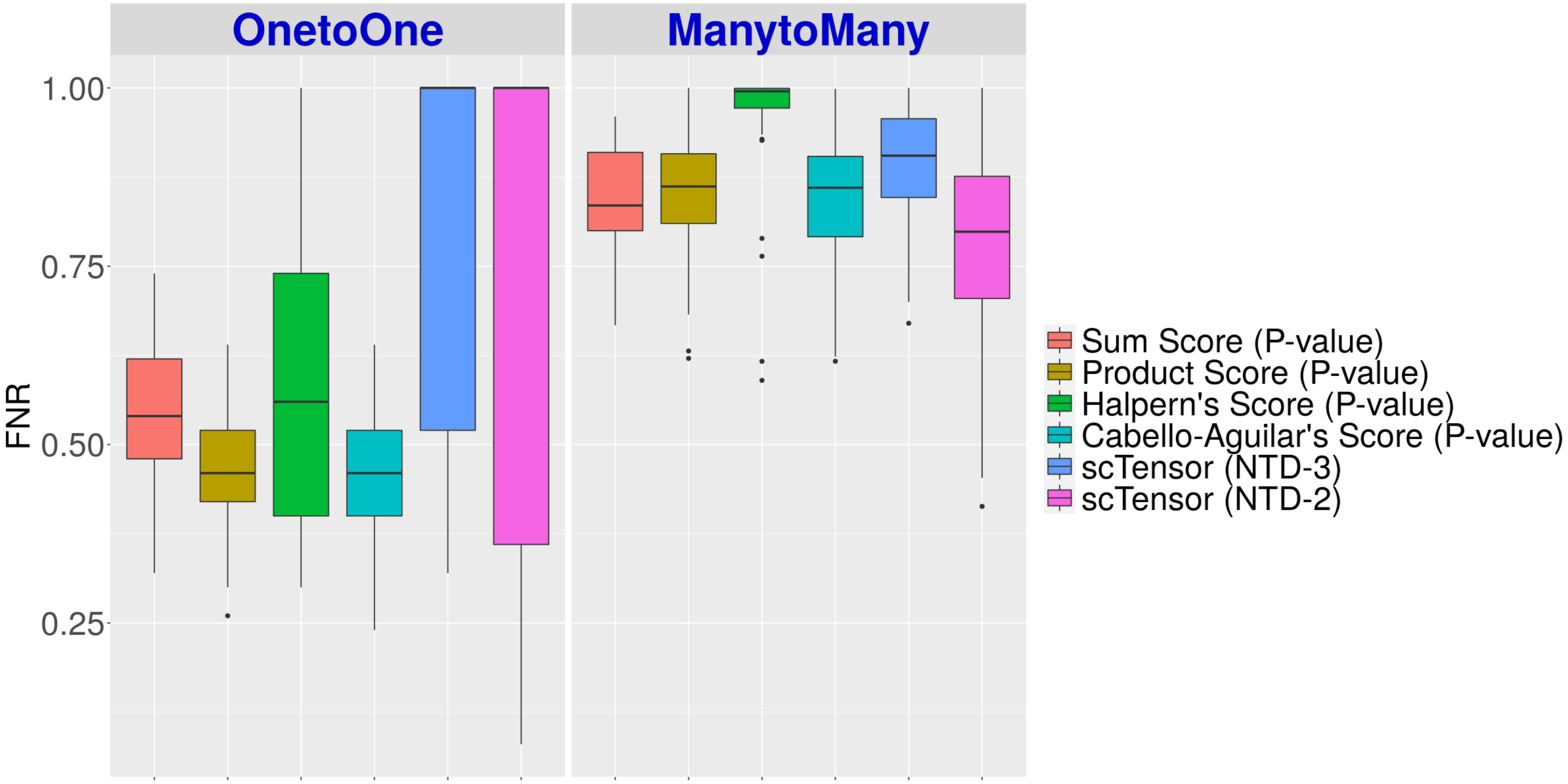

**E5 (Summary)** The value ranges 0 to 1 (the closer to 1, the worse)

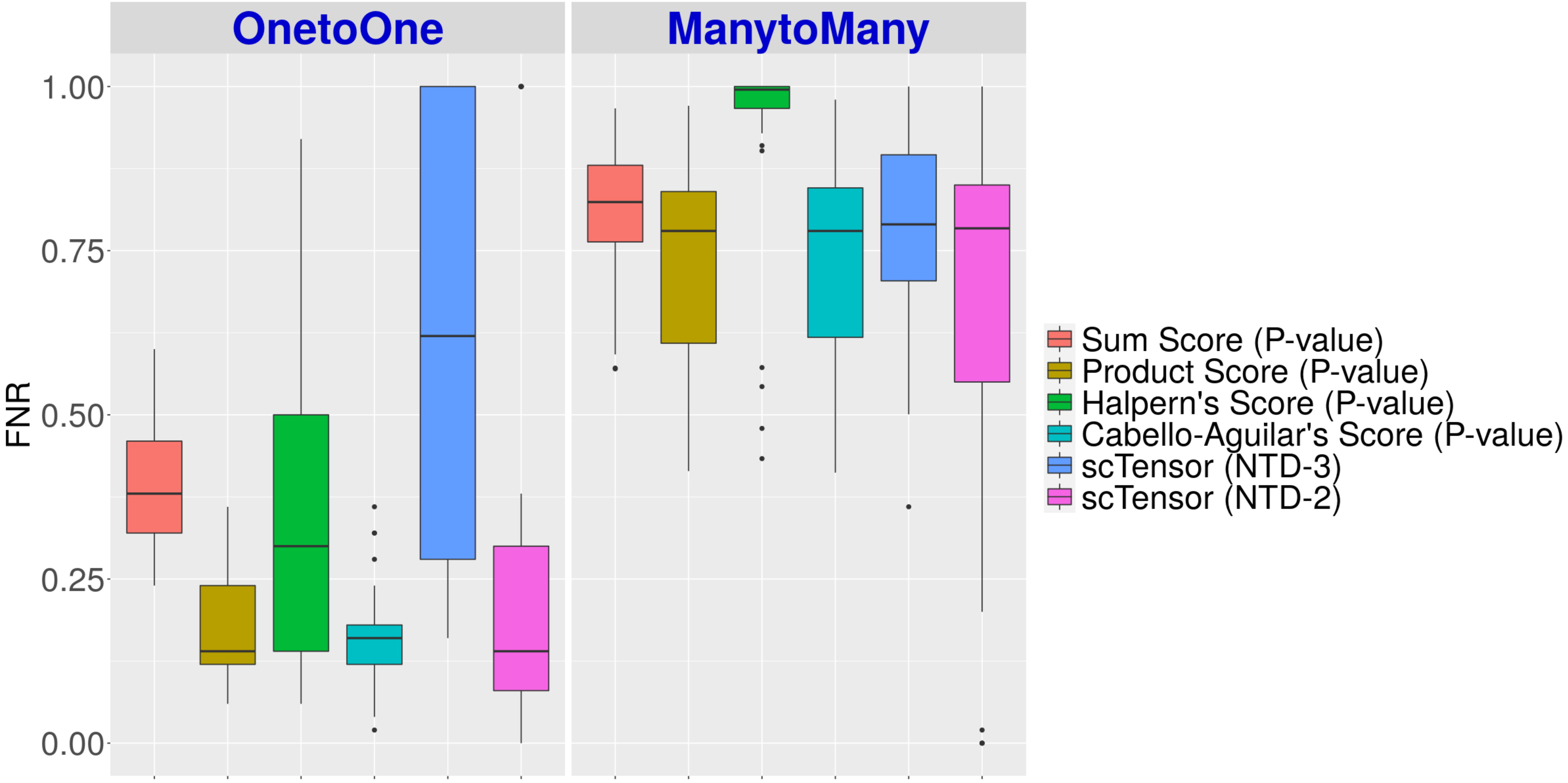

E10 (Summary) The value ranges 0 to 1 (the closer to 1, the worse)

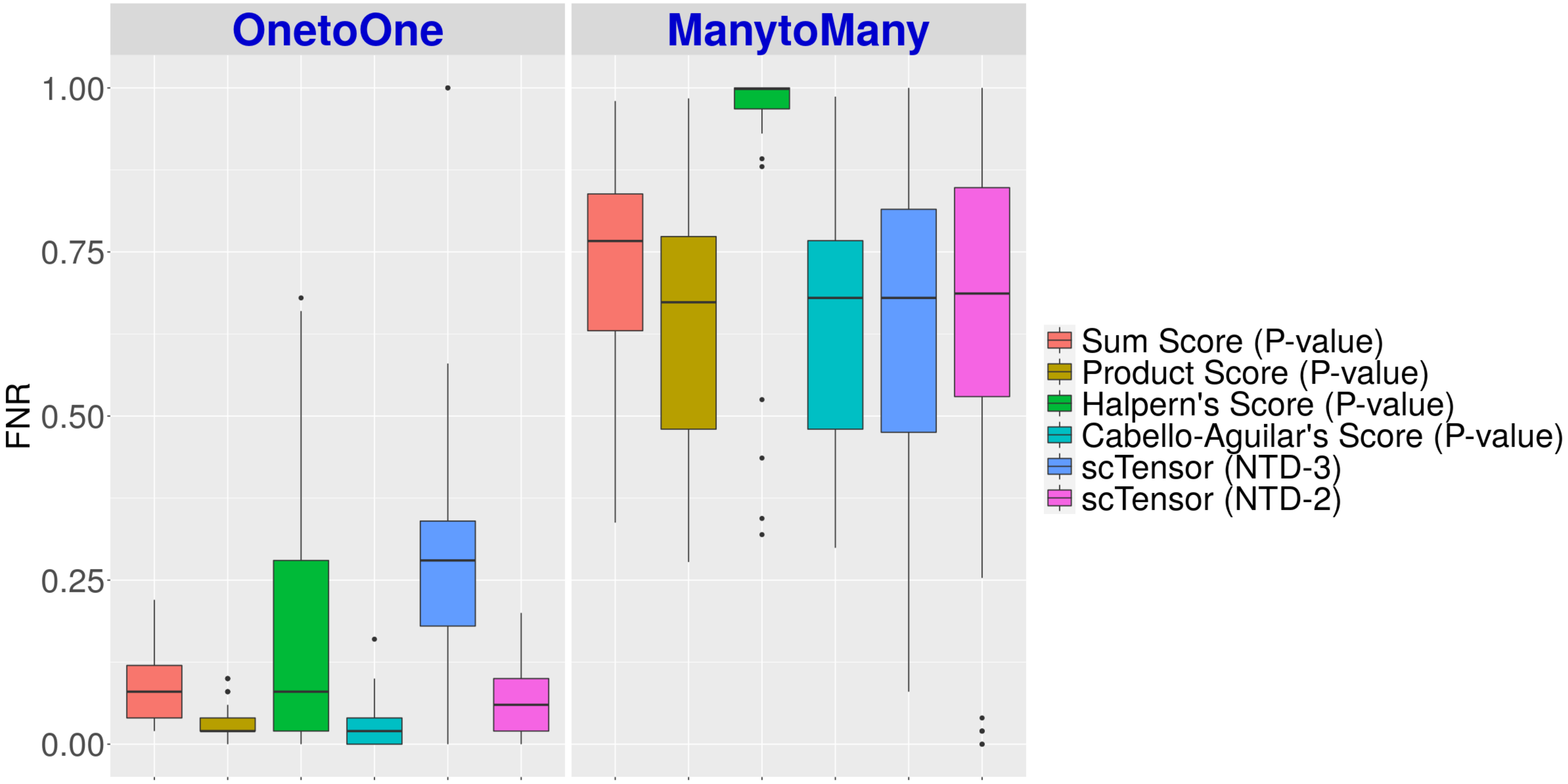

**E2 (Details)** The value ranges 0 to 1 (the closer to 1, the worse)

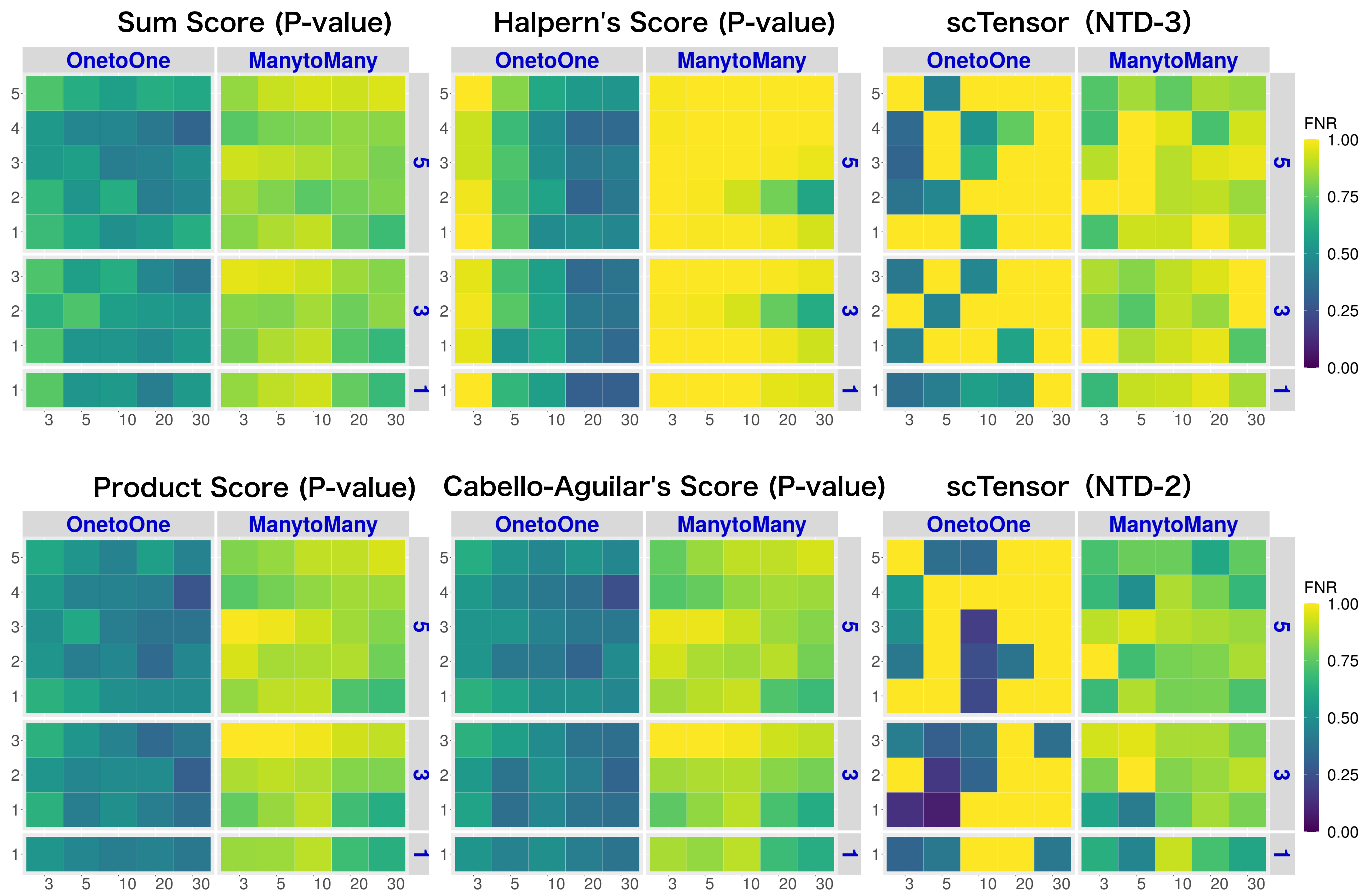

#### E5 (Details)

The value ranges 0 to 1 (the closer to 1, the worse)

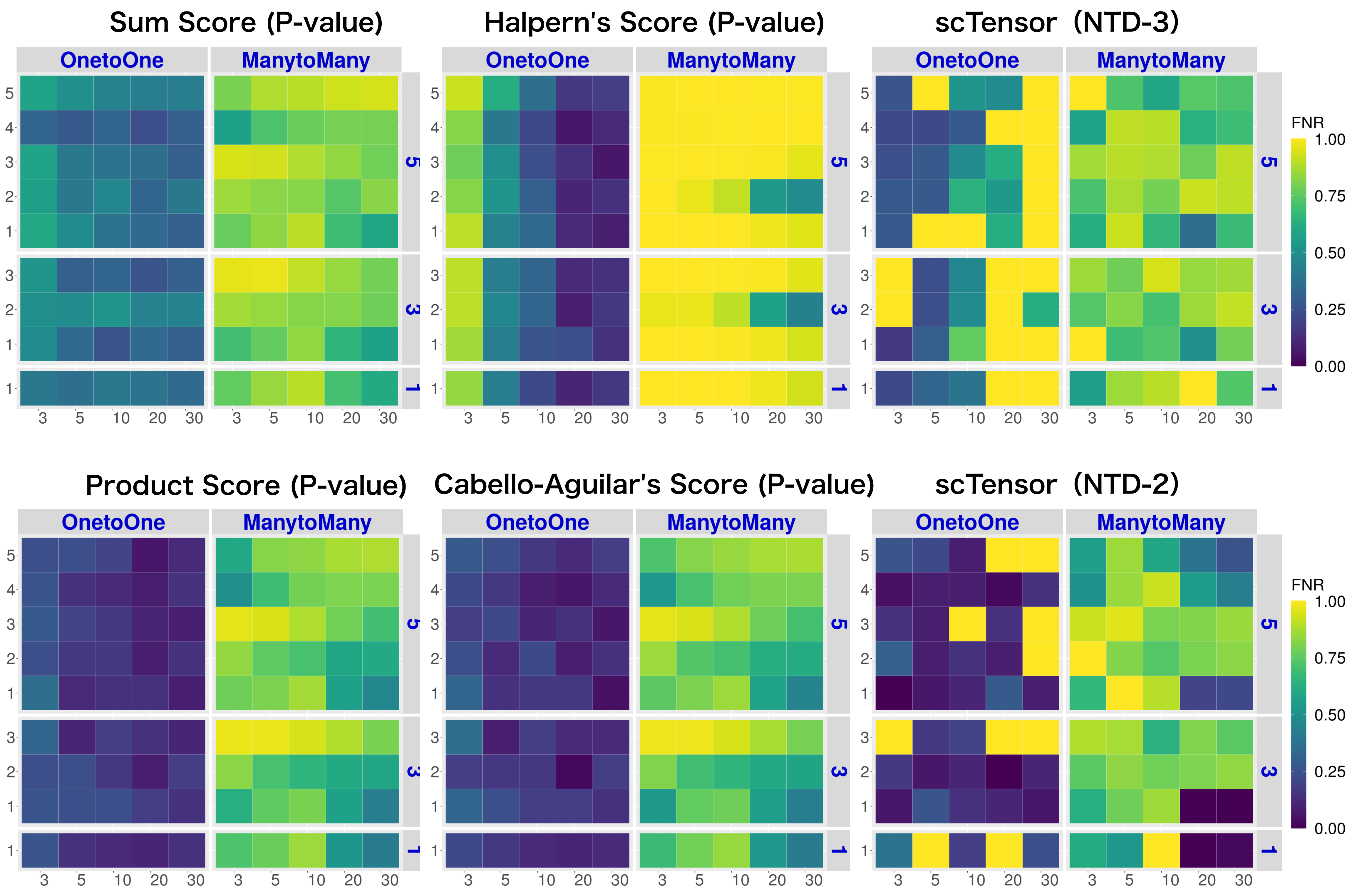

### E10 (Details)

The value ranges 0 to 1 (the closer to 1, the worse)

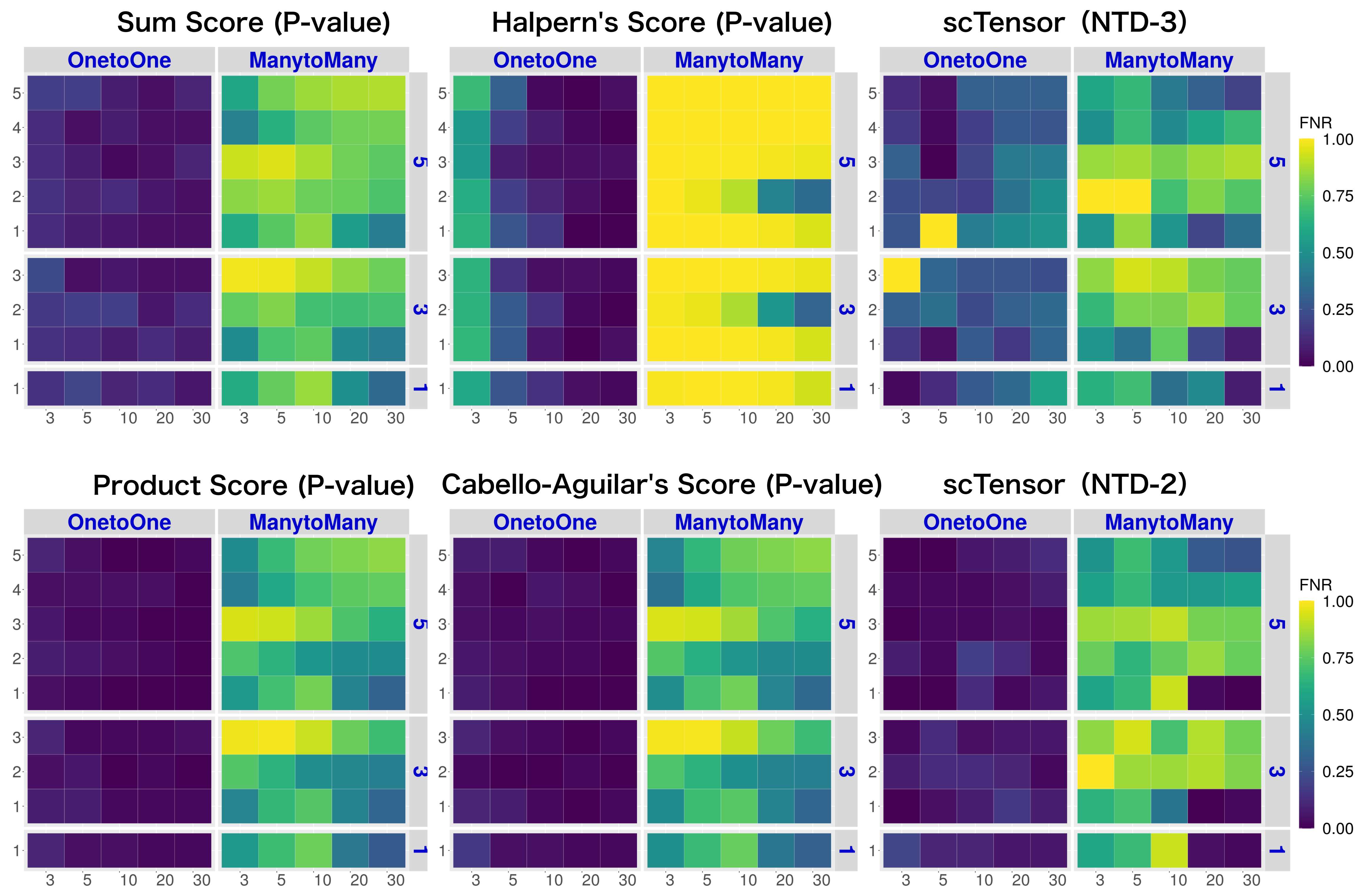

### Real Datasets

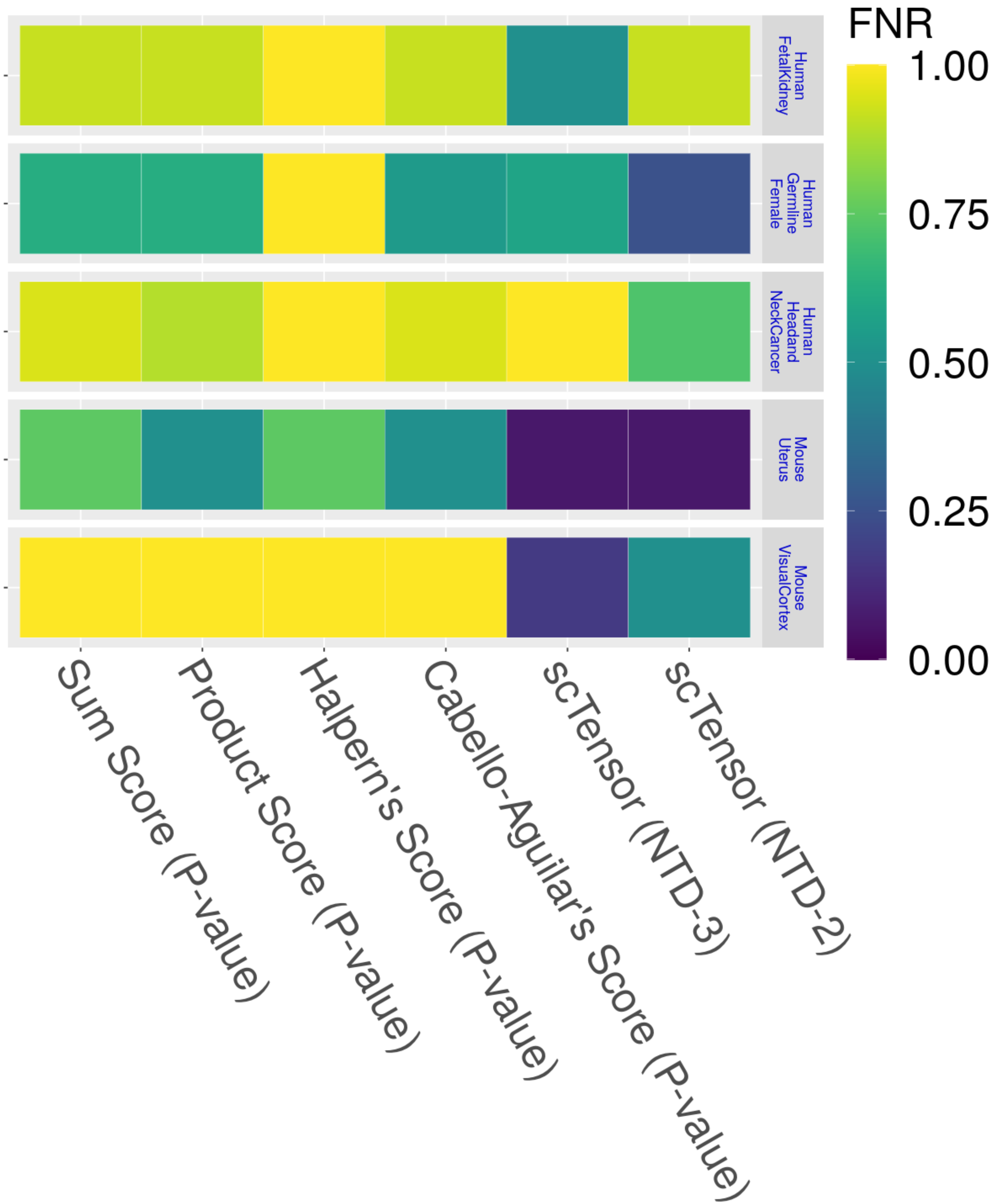

FNR
1.00
0.75
0.50
0.25
0.00
Human FetalKidney
Human Germline Female
Human Headand NeckCancer
Mouse Uterus
Mouse VisualCortex
Sum Score (P-value)
Product Score (P-value)
Halpern's Score (P-value)
Cabello-Aguilar's Score (P-value)
scTensor (NTD-3)
scTensor (NTD-2)
